# Elucidation of independently modulated genes in *Streptococcus pyogenes* reveals carbon sources that control its expression of hemolytic toxins

**DOI:** 10.1101/2022.08.04.502797

**Authors:** Yujiro Hirose, Saugat Poudel, Anand V. Sastry, Kevin Rychel, Richard Szubin, Daniel Zielinski, Hyun Gyu Lim, Nitasha Menon, Helena Bergsten, Satoshi Uchiyama, Tomoki Hanada, Shigetada Kawabata, Bernhard O. Palsson, Victor Nizet

## Abstract

*Streptococcus pyogenes* can cause a wide variety of acute infections throughout the body of its human host. The underlying transcriptional regulatory network (TRN) is responsible for altering the physiological state of the bacterium to adapt to each host environment. Consequently, an in-depth understanding the comprehensive dynamics of its TRN could inform new therapeutic strategies. Here, we compiled 116 existing high-quality RNA-seq data sets of *S. pyogenes* serotype M1, and estimated the TRN structure in a top-down fashion by performing independent component analysis (ICA). The algorithm computed 42 independently modulated sets of genes (iModulons). Four iModulons contained *nga-ifs-slo* virulence-related operon, which allowed us to identify carbon sources that control its expression. In particular, dextrin utilization upregulated *nga-ifs-slo* operon by activation of two-component regulatory system CovRS-related iModulons, and changed bacterial hemolytic activity compared to glucose or maltose utilization. Finally, we show that the iModulon-based TRN structure can be used to simplify interpretation of noisy bacterial transcriptome at the infection site.

## Introduction

*Streptococcus pyogenes* is a significant human pathogen responsible for over 700 million infections and at least 517,000 deaths annually worldwide^1^. This organism causes diverse diseases, ranging from superficial pharyngitis and impetigo to life-threatening invasive diseases, such as necrotizing fasciitis and streptococcal toxic shock syndrome^2^. As adaptation to many different host environments is reflected in its versatile pathogenicity, a comprehensive understanding of the dynamics of *S. pyogenes* gene regulation may be useful in guiding new therapeutic strategies.

An underlying transcriptional regulatory network (TRN) alters the physiological state of a bacterium to adapt to unique challenges presented by each host environment^3–7^. *S. pyogenes* regulons have been determined based on direct molecular methods, including transcriptomics performed on single gene knockout mutants^3,5,6,8–21^(PMIDs are listed in Supplementary_Data_3), and chromatin immunoprecipitation sequencing (ChIP-seq)^22,23^. At least 13 *S. pyogenes* two-component regulatory systems and 30 transcriptional regulators are known^24^. Despite these extensive and worthwhile efforts, it is often difficult to predict transcriptomic data from regulon annotations^25,26^. With many more potential interactions among each regulatory factor, comprehensive interpretation of the *S. pyogenes* global TRN has been difficult.

We previously reported an independent component analysis (ICA)-based framework that decomposes a compendium of RNA-sequencing (RNA-seq) expression profiles to determine the underlying regulatory structure of a bacterial transcriptome^27–29^. ICA computes independently modulated sets of genes (termed iModulons) and their corresponding activity levels under each experimental condition. The iModulons can be interpreted as data-driven regulons, since they rely on observed expression changes instead of predicted transcription factor binding sites. Therefore, it is possible that some iModulons reveal sets of genes never known to move coordinately. Moreover, since the number of iModulons is substantially fewer than the number of genes, they provide a helpful framework to more easily interpret the bacterial transcriptome. Here we apply ICA for the first time to elucidate the iModulon architecture of the major human pathogen *S. pyogenes*.

Among over 200 serotypes of *S. pyogenes*, serotype M1 is the most frequently identified from streptococcal pharyngitis^30^ and invasive diseases worldwide^31^. Todd-Hewitt medium supplemented with yeast extract (THY) is the most commonly used growth medium in which transcriptome profiling of *S. pyogenes* has been performed. To elucidate the TRN features of *S. pyogenes* serotype M1, we compiled 116 high-quality RNA-seq data sets of *S. pyogenes* serotype M1 cultured in THY and conducted ICA-based decomposition. ICA computed 42 iModulons that we characterized and analyzed their activities to formulate hypotheses. Users can search and browse all iModulons from this data set on iModulonDB.org, and view or interrogate them using interactive dashboards^32^.

Drilling down, we focused in this study on 4 iModulons that each include the *nga-ifs-slo* operon encoding an NAD-glycohydrolase, immunity factor, and the pore-forming cytolytic toxin streptolysin O that contribute to enhanced virulence of *S. pyogenes* ^33,34^. Comparing iModulon activities across 116 all samples allowed us to formulate hypotheses and identify carbon sources that control *nga-ifs-slo* expression. Although maltose and dextrin both consist solely of glucose molecules, dextrin utilization upregulated the *nga-ifs-slo* operon and changed bacterial hemolytic activity in contrast to glucose or maltose utilization. Furthermore, dextrin utilization induced the activation of iModulons related to the two-component regulatory system CovRS, a global transcriptional control system of *S. pyogenes* virulence phenotypes^2^. In the concluding part of this study, we show that the computed TRN structure from iModulon analyses can simplify the interpretation of noisy bacterial transcriptomes at the infection site using matrix projection. Our composite results suggest that *S. pyogenes*, in the inflammatory environment of the necrotizing fasciitis, senses and responds to both carbohydrate depletion and stresses affecting the CovRS system.

## Results

### 42 Biologically meaningful sets of genes are revealed from 116 high quality gene expression profiles by ICA

To extract regulatory signals from transcriptomic data, we used our established protocol^35^ and downloaded RNA-seq data from the public Sequence Read Archive^36^, where 229 RNA-seq samples were available for the M1 serotype of *S. pyogenes*. After quality control, which checks for read quality, alignment, and high replicate correlations, we selected 116 RNA-seq samples for ICA (**Supplementary Data 1**). These samples derived from from *S. pyogenes* propagated in THY under various culture conditions and exhibiting diverse expression states (**Supplementary Fig. 1**). Applying an extended ICA algorithm^27^, we decomposed the expression compendium into 42 iModulons and their activities (**Fig. 1A**) (**Supplementary Data 2**).

**Figure 1:**
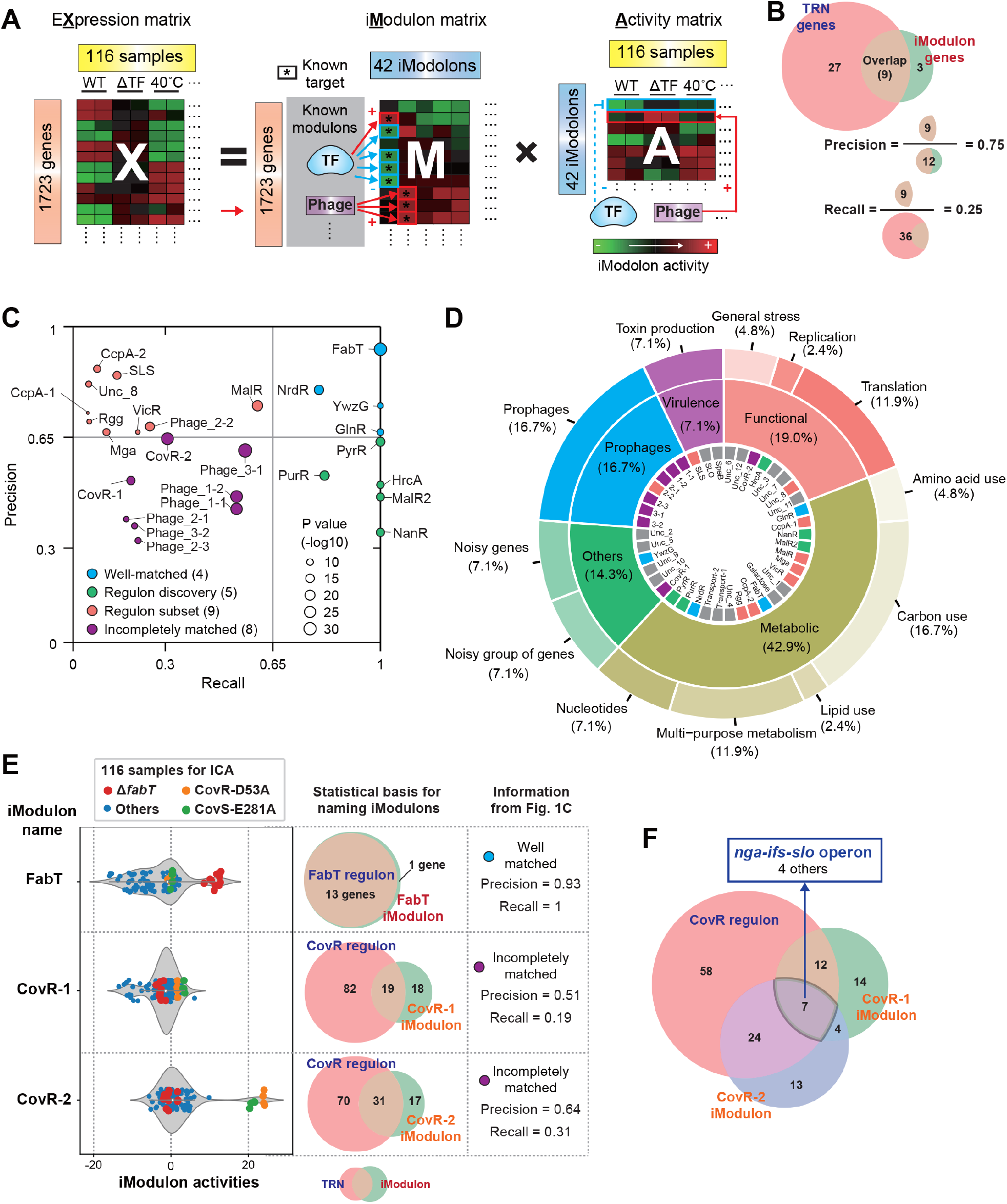
Independent component analysis (ICA) decomposes the compendium of transcriptomic data of *S. pyogenes* serotype M1 to biologically meaningful signals. (A) Schematic illustration of ICA applied to a gene expression compendium. ICA decomposes a transcriptomic matrix (X) into independent components (M) and their condition-specific activities (A). (B) The definitions of precision and recall with example numbers. (C) Scatter plot of precision and recall of the enrichments for the 26 (out of 42) iModulons that were matched to a regulon. The size of the circle indicates the negative log of P-values to the base 10. (D) Donut chart of iModulon functions. The outermost ring lists specific functions and the center ring lists broad functions. The coloring of innermost ring is reflected the colors of 26 regulatory iModulons as indicated in Fig. 1C, and gray indicates iModulons which don’t match to a regulon. (E) Examples to demonstrate the reliability of the annotated Modulons. Violin plot shows that FabT iModulon and CovRS-related iModulon activities (CovR-1 and CovR-2 iModulons) of 116 all samples. Transcriptomes from *fabT* deletion mutant strains (Red) are for the positive control of FabT iModulon. Transcriptomes from strains with CovR-D53A (Orange) and CovS-E281A (Green) are for the positive control of CovR-1 and CovR-2 iModulons. The overlap of iModulons with the known regulons are shown as Venn diagram. iModulons are detailed in Supplementary information 1. The list of the TRN is shown in Supplementary Data 3. The list of genes in each iModulons is shown in Supplementary Data 4. The imodulondb.org, where users can search and browse all iModulons from this data set and view them with interactive dashboards. (F) Overlapping genes among CovR regulon, CovR-1 iModulon, and CovR-2 iModulon.

Regulons are sets of co-regulated genes classified informally through scientific consensus on a variety of experimental results in the literature, while iModulons are derived solely from the measured transcriptome through an unbiased method^27^. Nevertheless, the known regulon structure of the *S. pyogenes* TRN described in RegPrecise^37^, PHASTER^38^, or previous reports (See **Supplementary Data 3**, or Materials and Methods for more details) is largely captured by the iModulons. Overall, 26 of the 42 iModulons were successfully mapped to a known regulator (**Supplementary Data 4**), and we named iModulons based on known regulons that share the highest overlap with the given iModulon, or by shared functionality of genes (e.g., cytolysins, transporter). The details for 42 iModulons are shown in **Supplemental Information 1** and **Supplementary Data 4**. They are also freely available to browse, search, or download on iModulonDB.org^32^.

Precision and recall between iModulons and regulons (**Fig. 1B**) were used to classify iModulons into four groups, namely well-matched, regulon subset, regulon discovery, and incompletely matched groups (**Fig. 1C**). (1) The well-matched group (n = 4) has precision and recall greater than 0.65. (2) The regulon subset (n = 5) exhibits high precision and low recall. They contain only part of their enriched regulon, perhaps because the regulon is very large and only the genes with the most pronounced transcriptional changes are captured. (3) A third group, regulon discovery (n = 5), has low precision but high recall. These iModulons contain some genes that are known to be co-regulated, along with other genes that are co-stimulated by the conditions in the data set. (4) The remaining enriched iModulons are termed incompletely matched (n = 8) because neither their precision nor their recall met the cutoff; however, this grouping still had statistically significant enrichment levels and appropriate activity profiles. The difference in gene membership between these iModulons and their regulons provide excellent targets for discovery.

The iModulons identified with no enrichments suggests that their composite gene sets may be regulated by unexplored transcriptional mechanisms or that there remains some noise due to large variance within the data. Functional categorization of iModulons provides a systems-level perspective on the transcriptome (**Fig. 1D**). Metabolism-related iModulons account for over 40% of the iModulons, while comparatively fewer iModulons deal with toxin production, stress response, replication, translation, and mobile genetic elements like prophages.

### iModulon activities of mutant strains demonstrate the reliability of the annotation of iModulons

All iModulons have a computed activity in every sample (**Supplementary Data 2**), allowing for easy comparisons of iModulon activities across samples (**Supplemental Information 1**). To benchmark the reliability of iModulon annotation, we show the FabT and CovRS-related iModulon activities (CovR-1 and CovR-2 iModulons) of 116 all samples (**Fig. 1E**).

The FabT iModulon overlaps nearly perfectly with the FabT regulon. This iModulon contains all genes known to be regulated by FabT, plus a hypothetical protein (XK27_06930) that this result indicates may be co-regulated with the known regulon. Additionally, transcriptomes from *fabT* deletion mutant strains (**Fig. 1E**, Red)^39^ show high FabT iModulon activities, consistent with the role of FabT as a repressor. Thus, this iModulon captures a known regulon and its expected behavior using transcriptome data alone. Several other iModulons are similarly easily explainable, particularly those in the well-matched group.

The activity of the CovRS transcription factor can be altered with exact point mutations. CovR-D53A or CovS-E281A point mutations induce considerable differential expression of genes (DEGs) in *S. pyogenes* transcriptome (171 or 139, respectively)^40^. Although the CovR-1 and CovR-2 iModulons incompletely overlap with the CovR regulon^6^ (**Supplementary information 1**), strains with CovR-D53A (**Fig. 1E**, Orange) and CovS-E281A (**Fig. 1E**, Green) show higher CovR-1 and CovR-2 iModulon activities compared to other samples. These correlations suggest that ICA results and iModulon annotations reflect previous findings properly and are useful for interpreting RNA-seq results.

Interestingly, the CovR regulon, CovR-1 iModulon, and CovR-2 iModulon overlap in 7 genes that contain *nga-ifs-slo* operon (**Fig. 1F**). Increased expression of the *nga-ifs-slo* operon is linked to enhanced virulence of *S. pyogenes*^33,34^.

### *nga-ifs-slo* operon is in SLO iModulon, MalR2 iModulon, CovR-1 iModulon, and CovR-2 iModulon

To decipher *S. pyogenes* pathogenicity, it is important to resolve the regulatory dynamics underlying two important virulence operons encoding potent cytolytic toxins: the abovementioned *nga-ifs-slo*, which encodes the gene for streptolysin O (SLO), and *sagA-I*, which encodes the biosynthetic machinery to produce streptolysin S (SLS), the toxin responsible for the hallmark β-hemolytic phenotype of the bacterium grown on blood agar. Both SLS and SLO are important for establishment of *S. pyogenes* skin infection and the formation of necrotizing skin lesions in invasive disease^41–43^.

Because iModulons are based on matrix decomposition, some genes may be present in multiple iModulons. The effects of all iModulons add together to reconstruct individual expression levels, which may be due to multiple TFs influencing the target genes. This is the case for the *nga-ifs-slo* operon, which is part of the designated SLO iModulon (**Fig. 2A**), MalR2 iModulon (**Fig. 2B**), CovR-1 iModulon, and CovR-2 iModulon (**Fig. 1F**). Indeed, the *nga-ifs-slo* operon are the only genes overlapping among these 4 iModulons (**Fig. 2C**). These findings indicate there are the multiple regulatory inputs governing expression of the *nga-ifs-slo* operon. In the subsequent sections, we speculate and validate environment cues under which the *nga-ifs-slo operon* is regulated based on iModulon information and corresponding activities.

**Figure 2:**
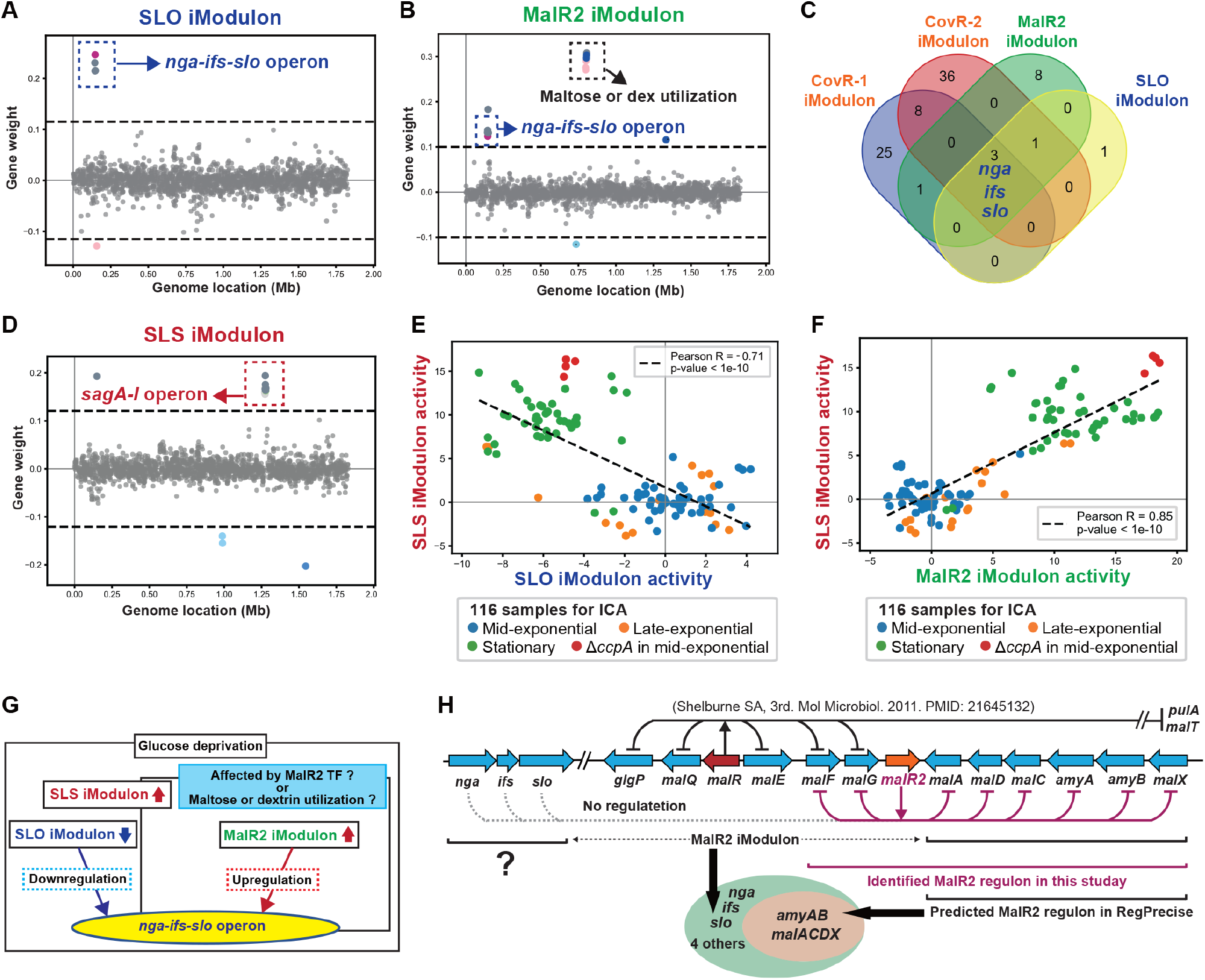
The *nga-ifs-slo operon* is in 4 iModulons, suggesting this operon is under multiple regulations. (A) Gene weights in SLO iModulon. Colored circles indicate clusters of orthologous groups of proteins (COG) categories, which are detailed in Supplementary information 1. The list of genes in each iModulons is shown in Supplementary Data 4. (B) Gene weights in MalR2 iModulon. (C) Overlapping genes among CovR-1 iModulon, CovR-2 iModulon, MalR2 iModulon, and SLO iModulon. (D) Gene weights in SLS iModulon. (E) Comparison between SLS and SLO iModulon activities across 116 samples. (F) Comparison between SLS and MalR2 iModulon activities across 116 samples. (G) Schematic diagram of the hypothesis that *nga-ifs-slo* operon is antagonistically regulated in specific condition. (H) Genomic organization of genes involved in maltose/dextrin transport, metabolism, and metabolic regulation. To identify the actual MalR2 regulon, transcriptomes of wild-type and *malR2* deletion mutant strains were compared at both the mid-exponential and stationary growth phase in THY broth (Supplementary Fig. 3, Supplementary Data 5).

### SLS and SLO iModulons suggests the antagonistic regulation between *sag A-I* and *nga-ifs-slo* operons in the stationary phase

The SLS iModulon contains the *sag A-I* operon encoding SLS (**Fig. 2D**). Among 42 iModulons, the SLS iModulon is the one containing all the *sagA-I* genes. Interestingly, there is a transcriptional tradeoff between the two major toxin-encoding virulence systems: the SLS iModulon activities are negatively correlated with SLO iModulon activities (**Fig. 2E**). SLS and SLO iModulon activities tend to be antagonized in samples obtained from stationary phase growth and a CcpA deletion mutant at mid-exponential phase (**Fig. 2E**). *S. pyogenes* stationary phase cultures in THY show evidence of glucose depletion^44^. CcpA senses carbohydrate availability ^11,45,46^. These observations lead us to hypothesize that the antagonistic regulation between *sag A-I* and *nga-ifs-slo* operons depends on a change in carbon source or TFs sensing the availability of carbon sources (**Supplementary Fig. 2**). SLO iModulon activities are also strongly deactivated in CcpA deletion mutants (**Fig. 2E**), suggesting CcpA deletion downregulates the *nga-ifs-slo* operon. However, other studies have indicated that the *ccpA* deletion in *S. pyogenes* serotype M1 induces the upregulation of the *slo* gene^6^ or does not change the expression level of the *slo* gene^46^. Therefore, the SLO iModulon alone is not enough to explain the regulation of the *nga-ifs-slo* operon.

### MalR2 iModulon harboring *nga-ifs-slo* operon is activated in the stationary phase, contrary to SLO iModulon

The MalR2 iModulon contains not only the *nga-ifs-slo* operon but also an operon for maltose or dextrin utilization (*malACDX, amyAB*) (**Fig. 2B**). Maltose is a disaccharide of D-glucose, whereas dextrin is a polysaccharide of D-glucose. Since maltose and dextrin are the products of the action of salivary amylases on dietary starch, it is possible *S. pyogenes* utilizes these carbohydrates during throat colonization or the development of pharyngitis.

Contrary to the SLO iModulon, MalR2 iModulon activities are positively correlated with SLS iModulons activities and are very high in samples from the stationary phase or CcpA deletion mutants at mid-exponential phase (**Fig. 2F**). These facts suggest that the transcription of the *nga-ifs-slo* operon is both downregulated (suggested by the SLO iModulon) and upregulated (suggested by the MalR2 iModulon) in stationary phase or under glucose-depleted conditions. These effects may counteract one another, resulting in the upregulation of the *slo* gene in the MalR2 iModulon-activated state even under conditions that deactivate the SLO iModulon (**Fig. 2G**).

### MalR2 TF does not contribute to the regulation of the *nga-ifs-slo* operon

The MalR2 iModulon is named due to a statistically significant overlap with the MalR2 regulon predicted by RegPrecise^37^ (**Fig. 2H**). However, the predicted MalR2 regulon in RegPrecise does not contain the *nga-ifs-slo* operon. In the genome of serotype M1 *S. pyogenes*, two genes predicted to encode LacI family TFs, *malR* and *malR2*, are situated near the *malACDX* and *amyAB* operons ^47,48^. The MalR regulon was experimentally defined using MalR-deletion mutant strains^5^, but no such data have been generated for MalR2.

To investigate whether the *nga-ifs-slo* operon is directly regulated by MalR2 TF, we compared the transcriptomes of wild-type (WT) and *malR2* deletion mutant (Δ*malR2*) strains at both mid-exponential and stationary growth phases in THY broth. Deletion of *malR2* induced the upregulation of only the *malACD* operon in mid-exponential phase (**Supplementary Fig. S3A, Supplementary Data 5**), but in stationary phase it upregulated both the *malACDFGX* and *amyAB* operons (**Supplementary Fig. S3B, Supplementary Data 5**). The *nga-ifs-slo* operon was not affected, which indicates that its inclusion with the MalR2 iModulon was due to co-stimulation, but not direct regulation. Therefore, we decided to explore the carbon sources that may stimulate the operon.

### Maltose or dextrin utilization changes bacterial hemolytic activity as compared to the glucose utilization

Despite the lack of direct MalR2 regulation, the *nga-ifs-slo* operon may still have been included in the MalR2 iModulon due to co-stimulation by inducers such as maltose or dextrin. To investigate whether maltose or dextrin utilization changes bacterial hemolytic activity, we used a red blood cell hemolysis assay (**Fig. 3A**). We adjusted the chemically defined medium (CDM) for *S. pyogenes* by following a previous report^49^. CDM without carbohydrate sources could not support the growth of serotype M1 *S. pyogenes* strain 5448, but its viability (bacterial colony forming units CFU) was maintained. Supplementation of the CDM with glucose, maltose, or dextrin allowed *S. pyogenes* 5448 strain growth (**Fig. 3B**). In a hemolysis assay using WT *S. pyogenes* supernatant, the glucose (+) condition produced a high hemolytic titer, while the maltose (+) or the dextrin (+) conditions yielded a low titer (**Fig. 3C**). To clarify whether this hemolysis was caused by SLS or SLO, we used SLS deletion mutant (Δ*sagA*), and SLO deletion mutant (Δ*slo*) strains (**Fig. 3D**). In the glucose supplemented condition, SLO is completely responsible for the hemolytic activity. In contrast, in the maltose or dextrin supplemented conditions, SLS is completely responsible for the hemolytic activity. In a hemolysis assay by using a live culture of the WT *S. pyogenes* strain, the glucose and dextrin conditions produced high hemolytic titer, while the carbon (−) and maltose conditions resulted in low titer (**Fig. 3E**). Here again, we used Δ*sagA* and Δ*slo* strains (**Fig. 3F**). In the glucose supplemented condition, SLO is principally responsible for the hemolytic activity, while SLS contributed only weakly. In other conditions, SLS is mainly responsible for the hemolytic activity. However, SLO-dependent hemolytic activity was confirmed in a dextrin supplemented condition, while it was not present in the carbon (−) and maltose conditions. These results show how supplementation of CDM with different carbon sources changes hemolytic activity and virulence factor expression in *S. pyogenes*. Therefore, we reasoned that the bacterial transcriptome obtained from similar experimental conditions might allow us to validate the hypothesis suggested in **Fig. 2G**.

**Figure 3:**
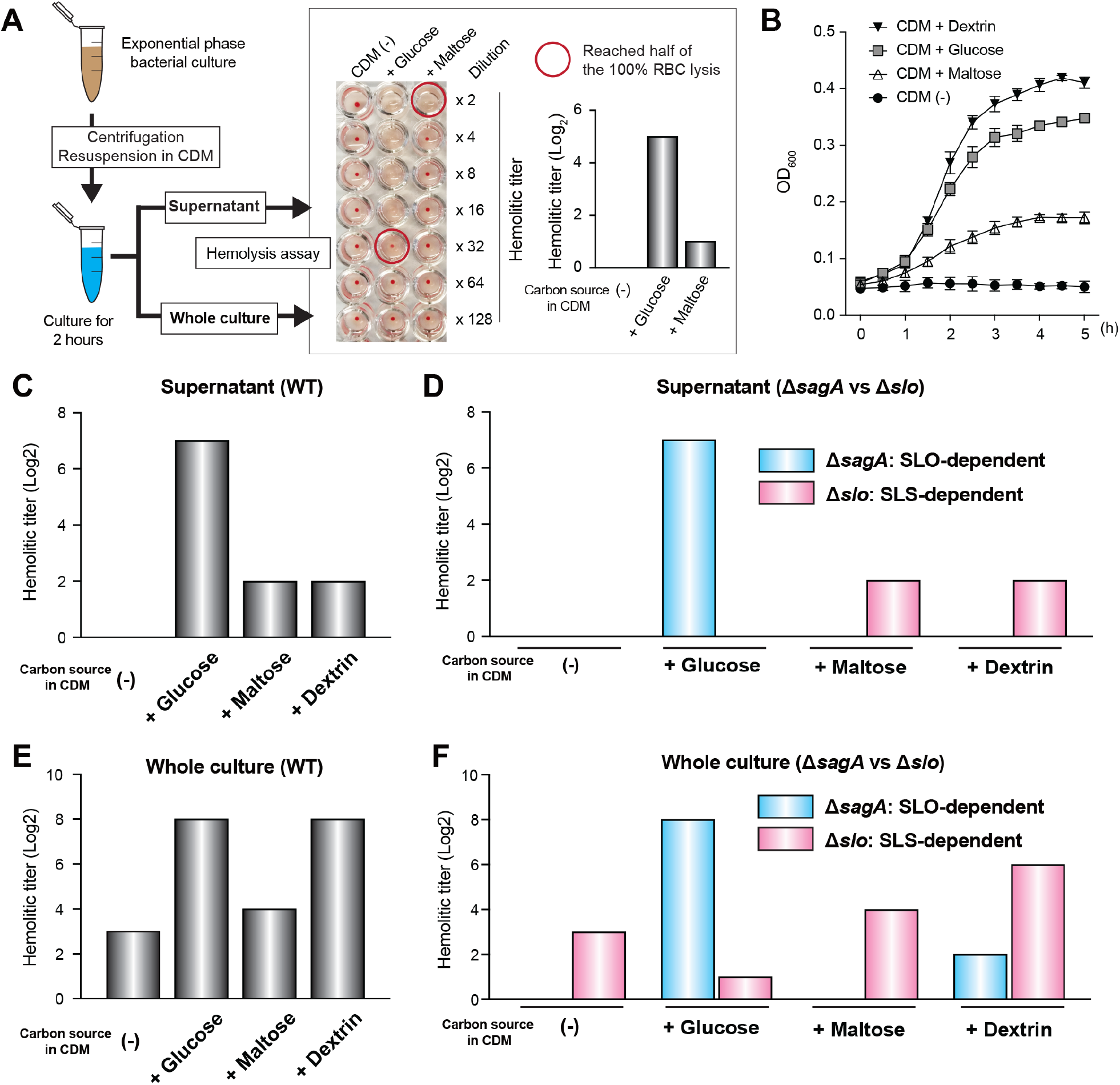
The switch between SLS and SLO activities is induced by the difference of supplemented carbon sources in CDM. (A) Experimental workflow for red blood cell hemolysis assay. (B) Growth curves in CDM supplemented with indicated carbon sources. (C) Hemolytic activity of the supernatant from WT. (D) Hemolytic activity of the supernatant from Δ*sagA* and Δ*slo* strains. (E) Hemolytic activity of the whole culture from WT. (F) Hemolytic activity of the whole culture from Δ*sagA* and Δ*slo* strains.

### Utilization of glucose, maltose, or dextrin significantly activates different iModulons, containing *nga-ifs-slo* operon

To examine the influence of different carbon sources on bacterial iModulon activities, we conducted RNA-seq analysis by using the RNA samples isolated from the *S. pyogenes* M1 strain after 2 h incubation in CDM (detailed conditions and results presented in **Supplementary Data 5**). Samples isolated from *S. pyogenes* cultured in CDM were evaluated together with those cultured in THY and shown earlier. In a hierarchical clustering (**Fig. 4A**) and principal component analysis (PCA, **Supplementary Fig. 4**) of the results, samples from CDM carbon (−) samples are located close to samples from stationary phase growth in THY. Therefore, the CDM carbon (−) condition seems to recapitulate the carbohydrate-depleted condition in THY. Although CDM glucose, maltose, and dextrin conditions supported *S. pyogenes* growth, samples from these conditions are located far from that mid-exponential phase in THY on the hierarchical clustering. Since THY broth is a nutrient-rich media containing many kinds of carbon sources, a CDM is better suited for assessing the influence of the supplementation of carbon sources.

**Figure 4.**
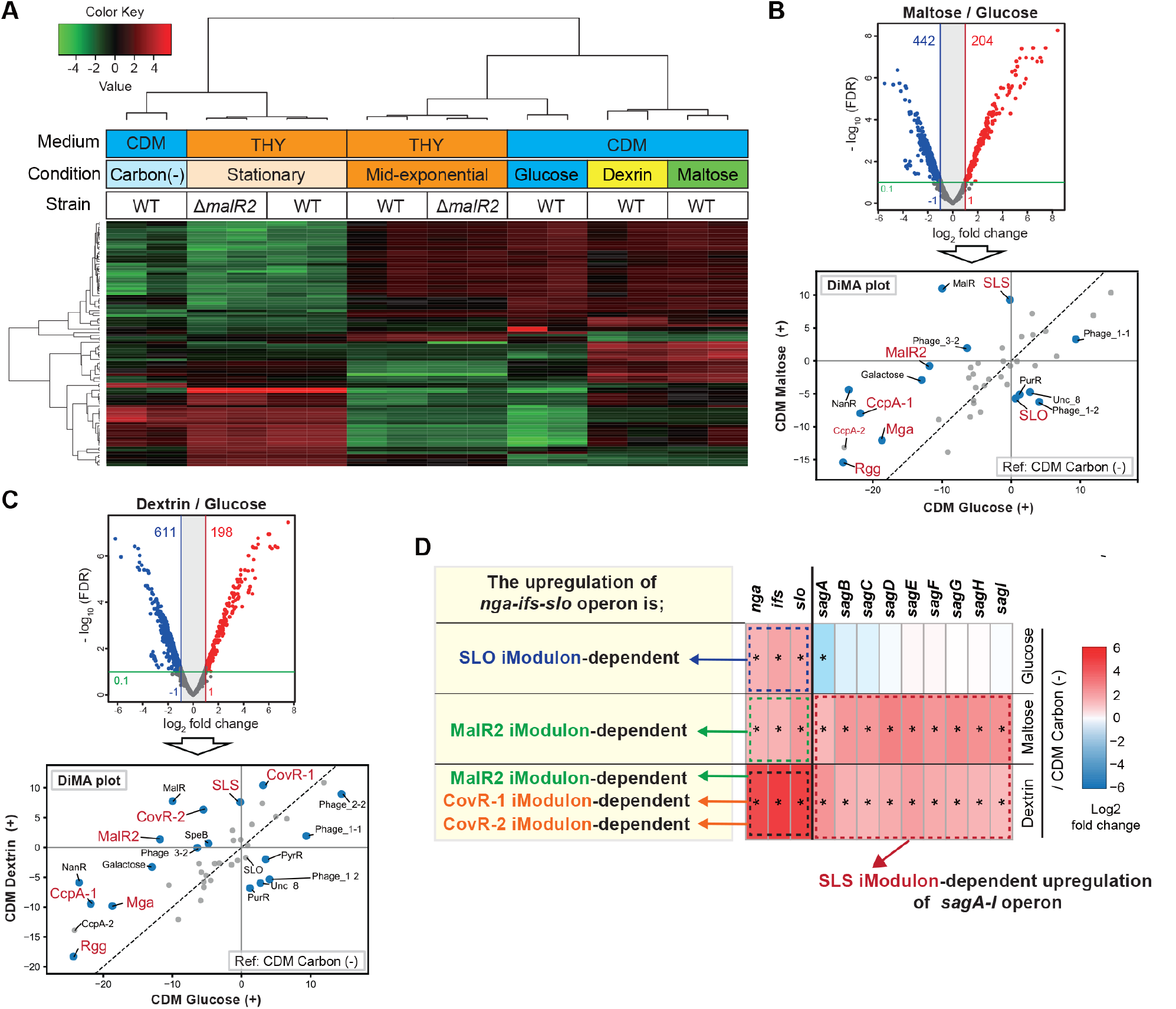
The activated iModulons that contribute to the upregulation of *nga-ifs-slo* operon differed by each carbon source. (A) Heat map of TPM data from 16 RNA-seq data set in this study. Heat map showing clustering of top 100 genes that contribute to the differences in each group. (B, C) Volcano plot and Differential iModulon Activity (DiMA) plot show transcriptome differences under the comparison conditions indicated in each figure. In volcano plots, colored circles indicate significantly upregulated (red) and downregulated (blue) genes (absolute log_2_ fold change, > 1; adjusted P < 0.1). In DiMA plots, blue circles indicate the significantly altered iModulons. (D) Expression levels of *nga-ifs-slo* and *sag A-I* operons in samples from CDM glucose (+), maltose (+), or dextrin (+) condition, as compared to those in samples from CDM carbon (−) condition. Based on the results of the DiMA plot, iModulons that contribute to the upregulation of *nga-ifs-slo* or *sag A-I* operons were determined.

In the CDM + maltose condition, a total of 648 *S. pyogenes* genes were altered in total compared to the CDM + glucose condition (**Fig. 4B, Supplementary Data 5**). Rather than anayzing so many DEGs individualy, we propose that iModulon activities can be used to facilitate interpretation of RNA-seq data. To that end, we calculated iModulon activities of samples in CDM + maltose and CDM + glucose conditions, which were centered to samples in CDM carbon (−) condition. After that, a Differential iModulon Activity plot (DiMA plot) allowed us to identify the significantly altered iModulons. In the CDM + maltose condition, antagonistic activation of the SLS and SLO iModulons was observed (**Fig. 4B**). The iModulons associated with the detection of carbohydrate depletion, including iModulons mapped to TFs of CcpA, MalR2, Mga^9^, and Rgg^10^, were likewise significantly activated in the CDM + maltose condition as compared to the CDM + glucose condition.

*S. pyogenes* grown in CDM + dextrin showed alterations in a total of 809 genes compared to the CDM + glucose condition (**Fig. 4C, Supplementary Data 5**). DiMA indicated that the significantly altered iModulons in the CDM + dextrin condition compared to CDM + glucose were similar to those significantly altered by maltose in **Fig. 4B**. However, SLO iModulon activity was not significantly altered. Notably, CovRS-related iModulons (CovR-1 and CovR-2) were significantly activated in the CDM + dextrin condition compared to CDM + glucose. Although the utilization of glucose, maltose, or dextrin each resulted in the upregulation of the *nga-ifs-slo* operon, the activated iModulons that contributed to operon upregulation differed by each carbon source (**Fig. 4D**).

### The stress environment *S. pyogenes* were exposed in necrotizing fasciitis (NF) seemed both the carbohydrate depletion and stresses affecting the CovRS regulator

The iModulons presented in this study help us to interpret RNA-seq data from other studies. When we perform an analysis in this way, we can make certain observations about how data behaves with respect to our TRN even if the data are of low quality. Here, we assess *S. pyogenes* transcriptome changes in the inflammatory environment of a murine model of necrotizing fasciitis (NF), in which bacterial RNA samples were isolated from infected hind limbs obtained at 24, 48, and 96 h post-infection^50^. At first, we calculated iModulon activities of samples isolated from NF at 24h, 48 h, and 96 h, which were centered to samples isolated from THY culture at mid-exponential phase (THY-ME, control). After that, we identified the significantly altered iModulons at each time point. Then, a set of 18 consistently altered iModulons were visualized (**Fig. 5A**). The iModulons associated with the detection of the carbohydrate depletion, such as CcpA-related, MalR, MalR2, Mga, and Rgg iModulons, were significantly activated in RNA samples isolated from NF *in vivo* compared to those obtained from THY-ME *in vitro*. In addition, CovRS-related iModulons (CovR-1 and CovR-2) are significantly activated in RNA samples isolated from NF compared to THY-ME.

**Figure 5:**
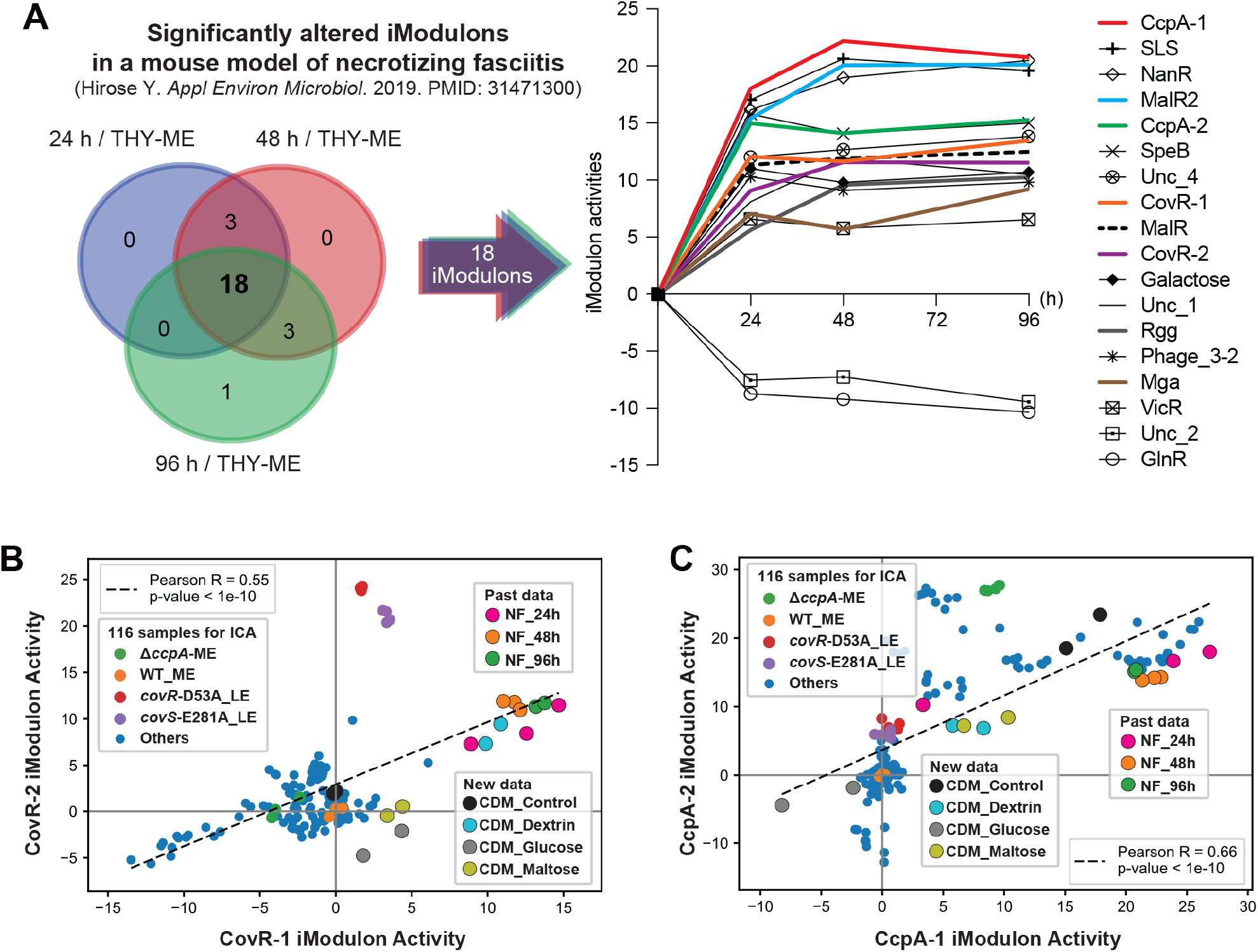
Bacterial iModulon activities in necrotizing fasciitis (NF) suggest that *S. pyogenes* sensed both the carbohydrate depletion and stresses affecting the CovRS regulator at the infected site. (A) Three-way Venn diagram illustrating the consistently altered iModulons during infection relative to samples isolated from THY culture at mid-exponential phase (THY-ME) (24 h versus THY-ME, 48 h versus THY-ME, and 96 h versus THY-ME). (B) Comparison between CovR-1 and CovR-2 iModulon activities across 116 samples, new samples, and past samples. (C) Comparison between CcpA-1 and CcpA-2 iModulon activities across 116 samples, new samples, and past samples.

To compare data from NF with new data obtained in this study, we also re-calculated iModulon activities of samples in CDM carbon (−), glucose, maltose, and dextrin conditions, which were centered to samples from THY-ME of the WT strain. Of interest, the activities of CovRS-related iModulons in RNA samples isolated from NF are located very close to that from CDM dextrin condition (**Fig. 5B**). In addition, the activities of CcpA-related iModulons in the CDM carbon (−) condition are located relatively close to that those in NF (**Fig. 5C**). These results suggest *S. pyogenes* in NF sensed both carbohydrate depletion and stresses affecting the CovRS regulator. Taken with our other results about regulation of the virulence factors, a coherent picture of carbon source depletion, CovRS, SLS, and SLO is presented. The most pronounced changes in the mouse model expression are in the same iModulons that we have been using to understand the regulation of virulence operons that contribute to NF pathology, which suggests this approach can be fruitful for studying in vivo disease models or clinical data.

## Discussion

In this study, 116 existing, high-quality RNA-seq data sets of *S. pyogenes* serotype M1 were decomposed using ICA. This decomposition identified 42 iModulons and their iModulon activities; 26 of the iModulons correspond to specific biological functions or transcriptional regulators. Based on the gathered information on iModulons and their activities across samples, we could formulate hypotheses and identify carbon sources that modulate hemolysis activity in *S. pyogenes*. Calculation of iModulon activities in new RNA-seq data sets also allowed us to identify that *S. pyogenes* activated CovRS-related iModulons in a CDM + dextrin condition (**Fig. 6**). Furthermore, by computing iModulon activities of published bacterial transcriptomes, we estimate the stress to which *S. pyogenes* is exposed at the infection site of NF.

**Figure 6:**
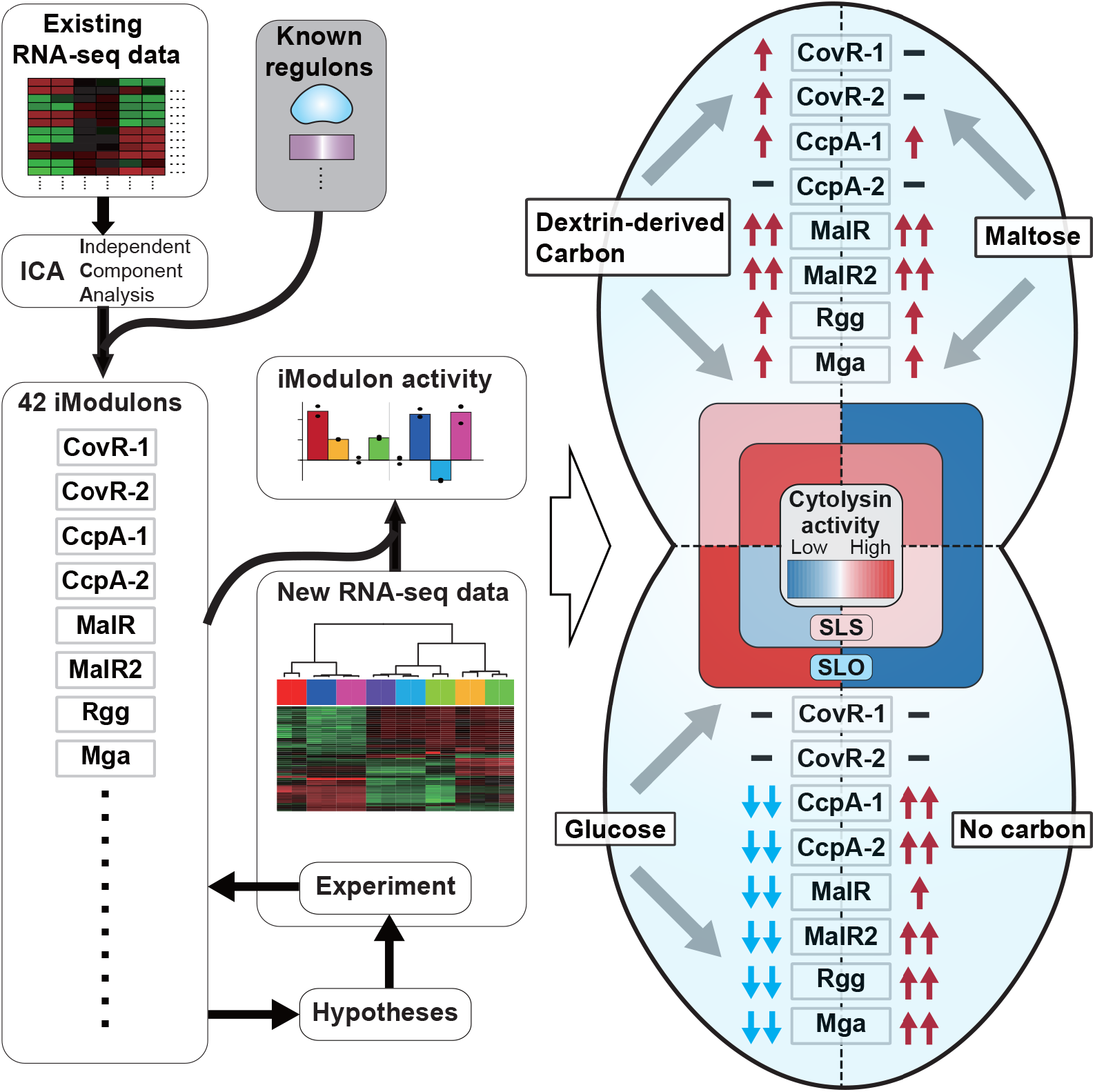
Independently modulated genes in *S. pyogenes* reveals carbon sources that change its hemolytic activity. In this study, we compiled existing high-quality 116 RNA-seq data sets of *S. pyogenes* serotype M1 and conducted ICA-based decomposition. This decomposition identified 42 iModulons and these activities, which allowed us to formulate hypotheses and identify carbon sources that change bacterial hemolysis activity. In particular, dextrin utilization changes bacterial hemolytic activity as compared to the maltose or glucose utilization. Furthermore, visualizing the iModulon activities in the transcriptome help us to interpret RNA-seq data. In particular, we identified that *S. pyogenes* activated CovRS-related iModulons in CDM dextrin (+) condition.

Some iModulons contained sets of genes that had never previously been reported as a regulon. This is likely because iModulons that are defined through an untargeted ICA-based statistical approach represent a set of genes that are co-expressed as a byproduct of multiple regulatory influences. By focusing on *nga-ifs-slo* operon, we could identify here that dextrin utilization alters bacterial hemolytic activity compared to maltose or glucose utilization. The full dataset is available for examination by researchers with interests in the biology and pathogenesis of *S. pyogenes* infections, allowing them to explore the contents of other iModulons in detail and the relations between them. At iModulonDB.org, users can search and browse all iModulons from this data set and view them with interactive dashboards^32^.

Previous reports have identified regulons using single gene-knockout mutants to conclude that the *nga-ifs-slo* operon is regulated by many TFs, including CovRS^6^, Mga^9^, Rgg^10^, and TrxR^3^. In our unbiased global study of available transcriptomes, only the MalR2 iModulon, CovRS-related iModulon and SLO iModulon contained the *nga-ifs-slo* operon. This distinction suggests that *nga-ifs-slo* operon is under multiple regulations at the same time. By comprehensively comparing iModulon activities across all samples, we speculate three environmental cues increase the expression of *nga-ifs-slo* operon: 1) Glucose-rich conditions (through activation of the SLO iModulon); 2) Glucose-depleted conditions (through activation of the MalR2 iModulon); and 3) environmental stress sensed by CovRS (through activation of CovRS-related iModulon). The existence of multiple mechanisms for the upregulation of *nga-ifs-slo* operon may link this operon with bacterial pathogenicity in various experimental conditions^33,34,51,52^.

Despite the structural similarities of glucose, maltose, and dextrin, *S. pyogenes* changed its hemolytic activity and transcriptome according to the carbon source. *S. pyogenes* in CDM + maltose or dextrin conditions activated iModulons associated with CcpA, MalR, and MalR2 TFs compared to those in CDM + glucose conditions. These TFs are repressors of the LacI/GalR family associated with carbon catabolite repression (CCR)^5,48^. Glucose is the primary carbohydrate source of energy for many bacteria, including *S. pyogenes*, which generates ATP from its conversion to pyruvate through glycolysis. However, utilization of maltose or dextrin requires energy input to first convert them to glucose in order to be metabolized. Therefore, it was believed that *S. pyogenes* suppressed genes involved in alternative carbohydrate utilization when glucose was available, leading to CCR^47,53^. In the physiological niche of *S. pyogenes*, the human body, carbon sources are available in different combinations and levels depending on the type of tissue and state of starvation and could be amenable to further studies with these tools.

We identified the the MalR2 regulon, *malACDFGX* and *amyAB* operons. Gene mutations of the *malR2* gene in *S. pyogenes* M1 strain significantly increase fitness in the nonhuman primate model of necrotizing myositis^49^. In another point of note, the *malR2, malACDX*, and *amyAB* genes exist in the M1, M2, M4 and M28 serotypes, but are absent from M3, M5, M6, M12, M18 and M49 serotypes^48^. The MalR2 iModulon may underpin differences in metabolic and virulence properties among serotypes of *S. pyogenes*. Maltose and dextrin are the products of the action of salivary amylases on dietary starch. AmyA, α-amylase, increased murine mortality following mucosal challenge^48^. It is possible that the products of salivary amylase may encourage upregulation of the *nga-ifs-slo* virulence-associated operon. It is presently unclear which particular carbon sources, concentrations, and enzymes are most utilized by the pathogen at infected sites.

CovRS-related iModulons were activated by dextrin utilization. Mutations of the CovRS virulence regulator derepress several different kinds of virulence genes, leading to a hypervirulent phenotype of *S. pyogenes* ^4,54^. Ikebe et al. reported that CovRS mutations in *S. pyogenes* were more common in clinical isolates from severe invasive infection compared to isolates from non-invasive infections^55^. CovRS mediates a general stress response in *S. pyogenes*, whereby specific environmental stimuli such as increased temperature, acidic pH, and high salt concentrations^2^. It had not been previously reported that dextrin utilization releases expression of CovRS-repressed genes; nevertheless, it remains unclear if dextrin utilization affects CovRS directly or indirectly, a subject meriting further study.

In this study, 116 all RNA-seq data sets for ICA come from *S. pyogenes* serotype M1 cultured in THY broth. Therefore, identified iModulons are specific for this most commonly studied media growth condition. Future studies can append other conditions to this data set and observe the changes to the set of iModulons that ensue. Although iModulon activities can often be explained by prior knowledge, they can also present surprising relationships that lead to the generation of hypotheses. We describe new information of iModulons and their activities enabled us to query metabolic and regulatory cross-talk, discover new potential relationships, find coordination between metabolism and virulence, and facilitates the interpretation of the *S. pyogenes* response *in vivo*. These versatilities may allow iModulons identified in this study or in near future to serve as a powerful guidepost to further our understanding of *S. pyogenes* TRN structure and dynamics.

## Materials and Methods

### Data acquisition and preprocessing for independent component analysis (ICA)

We downloaded RNA-seq data of *S. pyogenes* serotype M1 from NCBI SRA. The RNA-seq pipeline for conducting quality control (QC)/Quality assurance (QA) were applied by using a previously reported with minor modifications^56^. Briefly, the sequences were aligned to the *S. pyogenes* strain 5448 genome (Serotype M1, accession number CP008776), using Bowtie2^57^. The aligned sequences were assigned to open reading frames using featureCounts^58^. To reduce the effect of noise, genes with average counts per sample <10 were removed. The final counts matrix with 1,723 genes was used to calculate transcripts per million (TPM). To generate high-quality RNA-seq expression profiles, we removed RNA-seq samples which show the low-quality in FastQC^59^ and no <0.92 of Pearson R correlation between biological replicates. Finally, quality control passed 116 RNA-seq data were used for independent component analysis ^8,33,39,40,60,61^(Supplementary Data 1).

### ICA

Independent component analysis decomposes a transcriptomic matrix (X) into independent components (M) and their condition-specific activities (A) (Supplementary Data 2). The procedure for computing robust components with ICA has been described in detail previously^27^. Log2(TPM + 1) values were centered to strain-specific reference conditions and used as ICA decomposition. Next, Scikit-learn (v0.20.3) implementation of the FastICA algorithm was used to calculate independent components with 100 iterations, convergence tolerance of 10^−7^, log(cosh(x)) as contrast function and parallel search algorithm^27,62^. To determine the ideal number of components, we used our previously developed OptICA method^63^. The resulting M matrix containing source components from the 100 iterations were clustered with Scikit-learn implementation of the DBSCAN algorithm with ε of 0.1, and minimum cluster seed size of 50 samples (50% of the number of random restarts). If necessary, the component in each cluster was inverted such that the gene with the maximum absolute weighting the component was positive. Centroids for each cluster were used to define the final weightings for M and corresponding A matrix. The whole process was repeated 100 times to ensure that the final calculated components were robust. Finally, components with activity levels that deviated more than five times between samples in the same conditions were also filtered out.

### Determining independently modulated sets of genes (iModulons)

ICA enriches components that maximize the nongaussianity of the data distribution. While most genes have weightings near 0 and fall under Gaussian distribution in each component, there exists a set of genes whose weightings in that component deviate from this significantly. To enrich these genes, we used Scikit-learn’s implementation of the D’Agostino K2 test, which measures the skew and kurtosis of the sample distribution (D’Agostino 1990). We first sorted the genes by the absolute value of their weightings and performed the K2 test after removing the gene with the highest weighting. This was done iteratively, removing one gene at a time, until the K2 statistic falls below a cutoff. We calculated this cutoff based on sensitivity analysis on agreement between enriched iModulon genes and known regulons. Regulons predicted by RegPrecise^37^(*S. pyogenes* M1 GAS), shown in previous reports (PMIDs are listed in Supplementary_Data_3), and prophage genes searched by PHASTER^38^ are used for information of known regulons (Supplementary Data 3). For a range of cutoff, we ran the iterative D’Agostino K2 test on all components and checked for statistically significant overlap of iModulons with the known regulons using Fisher’s exact test (Supplementary Data 4). For iModulons with significant overlap, we also calculated precision and recall. The cutoff of 150, which led to the highest harmonic average between precision and recall (F1-score), was chosen as the final cutoff. We also manually identified iModulons that consisted of genes with shared functions (e.g., cytolysins, transporter) or those that corresponded to other genomic features (Supplementary information 1, Supplementary Data 4).

### Bacterial strains and culture conditions

*S. pyogenes* M1T1 strain 5448 (accession no. CP008776) was isolated from a patient with toxic shock syndrome and necrotizing fasciitis and considered to be a genetically representative globally disseminated clone associated with the invasive infections^64^. *S. pyogenes* were grown at 37°C in a screw-cap centrifuge tube (BD Biosciences, San Jose, CA, USA) filled with Todd-Hewitt broth supplemented with 0.2% yeast extract (THY) (Hardy Diagnostics, Santa Maria, CA, USA) in an ambient atmosphere and standing cultures. To obtain cultures for experiments, overnight cultures of *S. pyogenes* were back diluted 1:50 into fresh THY broth and grown at 37°C, with growth monitored by measuring optical density at 600 nm (OD_600_ = 0.35 for mid-exponential samples, or OD_600_ = 0.75 for early-stationary samples). CFUs were determined by plating diluted samples on THY agar.

*Escherichia coli* MC1061 strain was used as a host for derivatives of plasmids pHY304^65^. *E. coli* strains were cultured in Luria-Bertani medium (Hardy Diagnostics) at 37°C with agitation. For selection and maintenance of strains, antibiotics were added to the medium at the following concentrations: erythromycin, 500 μg/mL for *E. coli* and 2 μg/mL for *S. pyogenes*; chloramphenicol, 2 μg/mL for *S. pyogenes*.

### Construction of *malR2* mutant strain

An in-frame *malR2* transcription factor deletion mutant (Δ*malR2*) with a background of strain 5448 (WT) were constructed using the pHY304 temperature-sensitive shuttle vector, as previously reported^66^. Briefly, a pHY304-*malR2*KO plasmid harboring the DNA fragment, in which upstream of the *malR2* gene, a chloramphenicol acetyltransferase gene (*cat*), and downstream regions of the *malR2* gene were linked by overlapping PCR, was electroporated into WT strain and grown in the presence of erythromycin. The plasmid was then integrated into the chromosome via first allelic replacement at 37°C, after which it was cultured at 30°C without antibiotics to induce the second allelic replacement. The deletion of *malR2* was confirmed by site-specific PCR using purified genomic DNA. Primers are listed in Supplementary Table 1.

### Red blood cell hemolysis assay

Human blood was collected by using BD Vacutainer EDTA (Cat. 366643m Becton, Dickinson, Mountain View, CA, USA) tubes. Red blood cells (RBC) were isolated by centrifuging and washing with PBS. Bacterial cultures in THY broth at OD600 = 0.3-0.4 were centrifuged by using eppen tubes and resuspended in equal amount of condition defined medium (CDM)^49^. CDM was supplemented with 4.5 g/L of D-glucose (Sigma-Aldrich, St Louis. MO, USA; Cat. G7021), D-maltose (Sigma-Aldrich; Cat. M5885), or dextrin (Sigma-Aldrich; Cat. 31410). At 2 hours post-incubation at 37 C°, 150 μL of wholes culture or supernatants were collected for serial dilutions in PBS. All wells were added 50 μL of 2% RBC in PBS and incubated 4 hours. Supernatants were collected from assay wells after centrifugation at 3000 × g for 15 min, and hemolysis determined by absorbance with a SpectraMax M3 plate reader at 541 nm using SoftMax Pro software. Each titer was recorded as the point that the hemolysis reached half of the 100% RBC lysis (H_2_O) control.

### RNA Extraction and Library Preparation

The details of new 16 RNA samples analyzed in this study are shown in Supplementary Data 5. All samples were collected in biological duplicates, originating from different overnight cultures. 3 mL samples were added to tubes containing 6 mL RNAprotect Bacteria Reagent (Qiagen, Hilden, Germany) and vortexed. After 5min incubation at room temperature, they were centrifuged to remove the supernatant. RNA was extracted from the pelleted cells using a Quick RNA Fungal/Bacterial Microprep kit (Zymo Research, Irvine, CA, USA). The cells were mechanically lysed with Mini-Beadbeater-96 (Biospec Products, Bartlesville, OK, USA) for 2 min and DNA was removed with DNase I (New England Biolabs, Beverly, MA, USA). during the RNA purification. RNA quality was checked with an Agilent Bioanalyzer (Agilent Technologies, Santa Clara, CA, USA) instrument. Ribosomal RNA (rRNA) was removed using RiboRid^67^ with designed probes for *S. pyogenes* M1T1 strain 5448 (Supplementary Data 6). The library was prepared with KAPA Hyper Prep kit (KAPA Biosystems, Wilmington, MA, USA) following the manufacturer’s protocol.

### RNA-seq and data analysis

Libraries were sequenced using Illumina NextSeq systems, with 40-bp paired-end reads (sequenced at the University of California Davis DNA sequencing facility). Raw reads were trimmed using the fastp^68^, mapped to the *S. pyogenes* strain 5448 genome, and used to calculate the transcripts per kilobase million (TPM) with the commercially available Geneious Prime 2019.2 software package (Biomatters, Auckland, New Zealand). Differential expression and global analyses of RNA-seq expression data were performed using iDEP (ge-lab.org). EdgeR log transformation was used for clustering and PCA (iDEP). Hierarchical clustering was visualized using the average linkage method with correlation distance (iDEP). Functional annotations of *S. pyogenes* strain 5448 genome are obtained by using EggNOG (eggnog5.embl.de) or PATRIC (www.patricbrc.org).

## Data and Code Availability

The data accession numbers can be found in Supplementary Data 1. The normalized log TPM (X), and the calculated M and A matrix of the model can be found in Supplementary Data 2. The code for QC/QA, ICA, and assessment of the regulator enrichment can be found on Github (https://github.com/avsastry/modulome-workflow). Python package for analyzing and visualizing iModulons (PyModulon) can be also found on Github, (https://github.com/SBRG/pymodulon). All new 16 RNA-seq data obtained in this study have been deposited into DDBJ sequence read archive (DRA) under accession number DRA014564. Interactive online dashboards for all iModulons and all data are available at https://imodulondb.org under the data set name “*S. pyogenes*”.

## Acknowledgments

We acknowledge the University of California Davis DNA sequencing facility for their support with RNA sequencing. This study was supported in part by AMED (JP21fk0108044, JP21fm0208007), Japanese Society for the Promotion of Science (JSPS) KAKENHI (grant numbers 20K18474, 20KK0210, and 22K09924), JSPS Overseas Research Fellowships. The funders had no role in study design, data collection or analysis, decision to publish, or preparation of the manuscript.

