## Supplementary information 1 for "Elucidation of independently modulated genes in *Streptococcus pyogenes* reveals carbon sources that control its expression of hemolytic toxins"

### FabT iModulon

**Significantly overlapped TRN: FabT > LacD.1_M14**

**Biological function: Fatty acid biosynthesis**

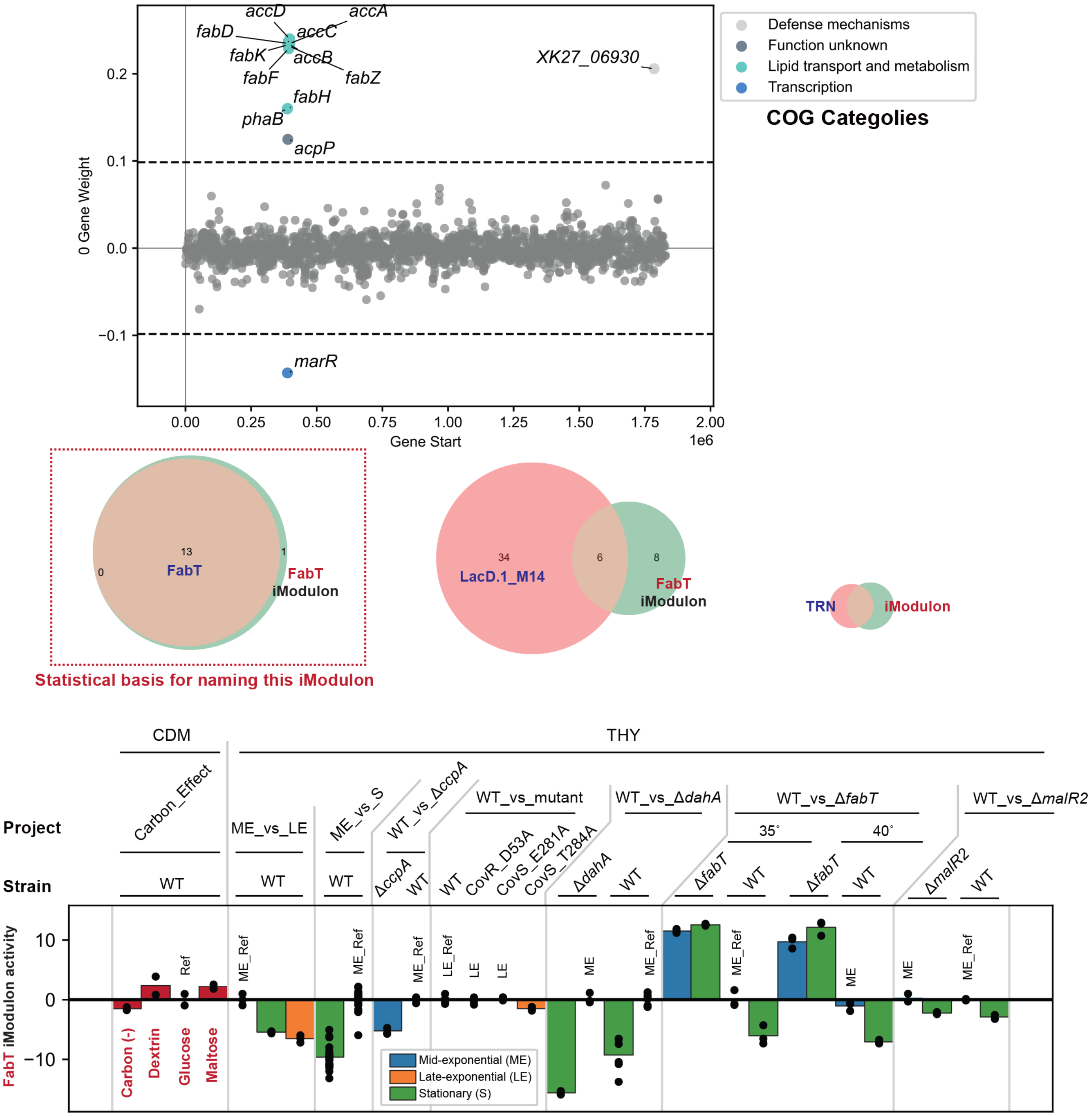

### CovR-1 iModulon

**Significantly overlapped TRN:**

**CovR_1_ME_M1 > Mga_low_Glucose_M1 > CovR_CcpA_M1 >Rgg_exponetional_M49 > CodY_M49**

**Biological function: Phathogenesis and the peptide transport**

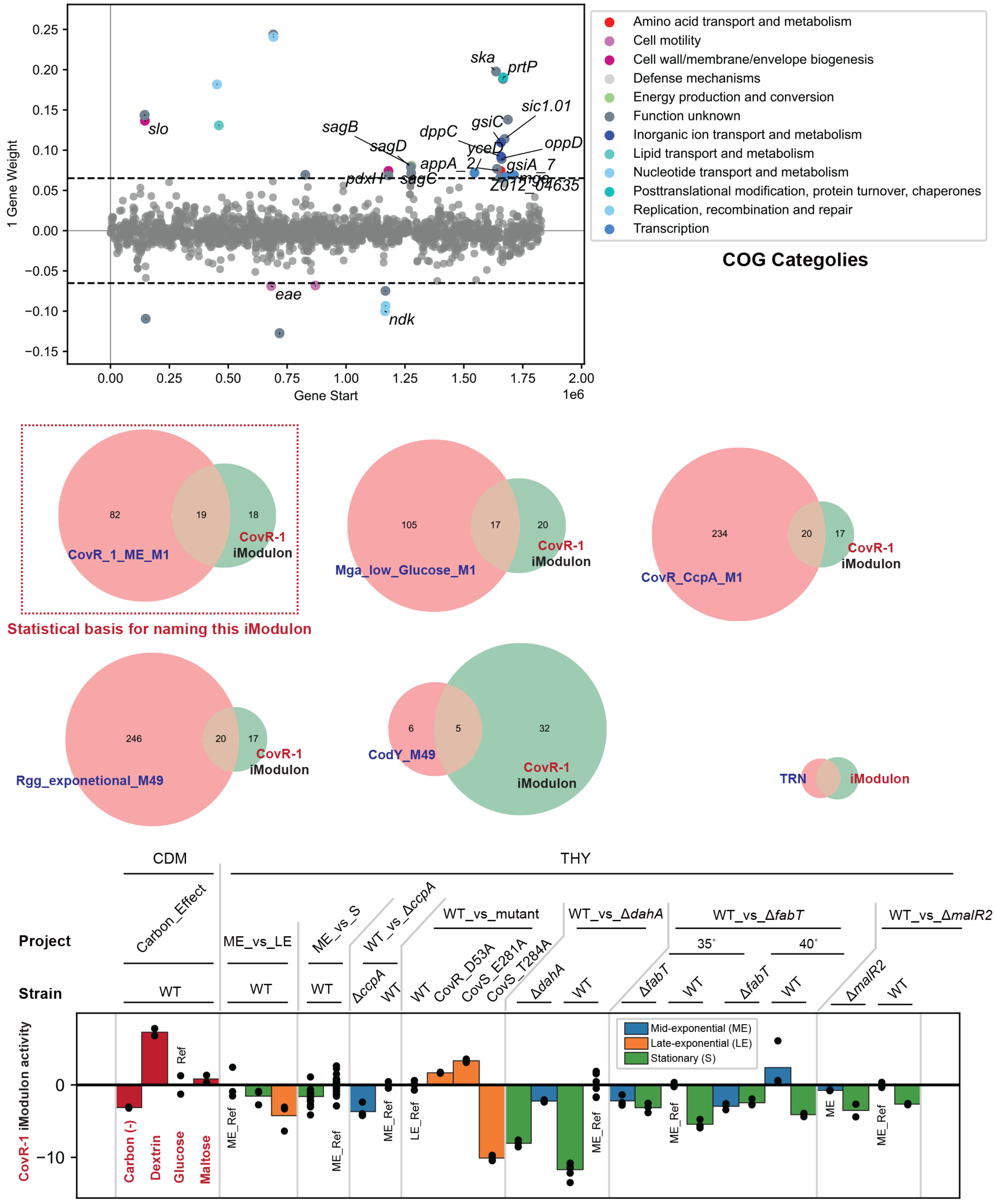

### Phage_1-1 iModulon

**Significantly overlapped TRN: Phage_1 > SPy1285**

**Biological function: Prophages**

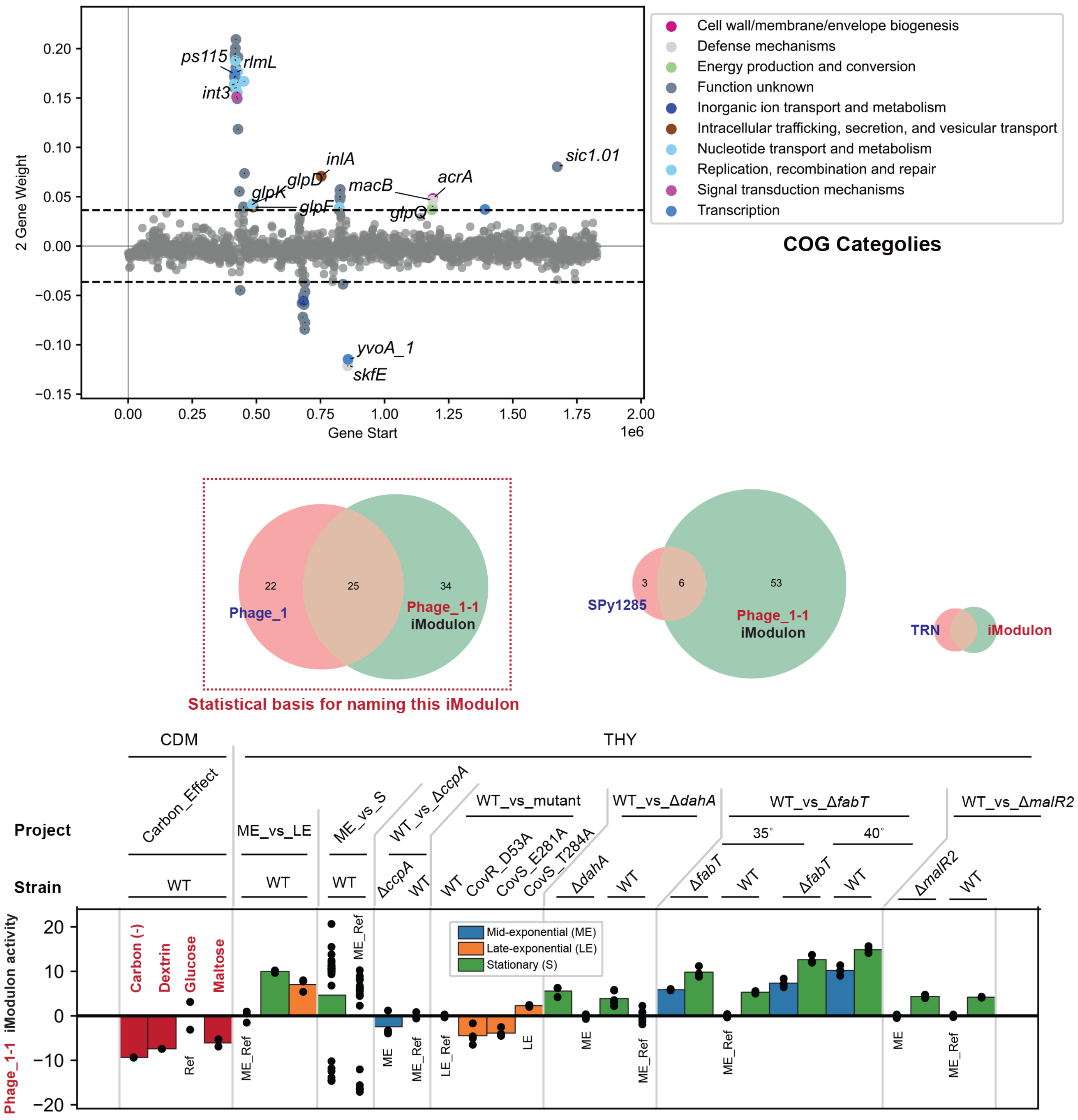

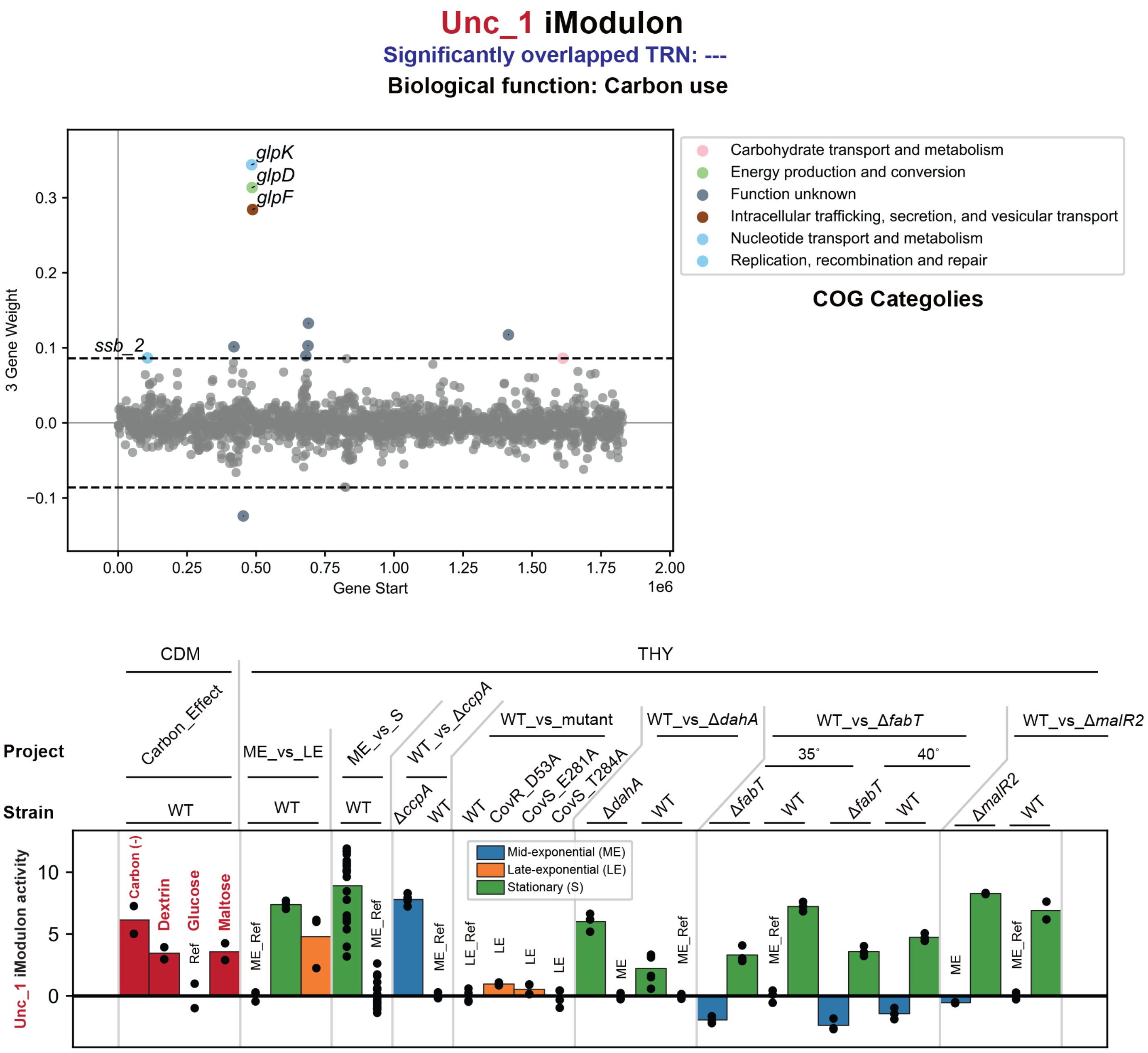
Unc_1 iModulon
Significantly overlapped TRN: ---
Biological function: Carbon use
3 Gene Weight
Gene Start
1e6
glpK
glpD
glpF
ssb_2
Carbohydrate transport and metabolism
Energy production and conversion
Function unknown
Intracellular trafficking, secretion, and vesicular transport
Nucleotide transport and metabolism
Replication, recombination and repair
COG Categolies
CDM
THY
Project
Strain
Carbon_Effect
ME_vs_LE
ME_vs_S
WT_vs_ΔccpA
WT_vs_mutant
WT_vs_ΔdahA
WT_vs_ΔfabT
WT_vs_ΔmalR2
35°
40°
WT
ΔccpA
CovR_D53A
CovS_E281A
CovS_T284A
ΔdahA
ΔfabT
ΔmalR2
Unc_1 iModulon activity
Carbon (-)
Dextrin
Glucose
Maltose
Ref
ME_Ref
LE_Ref
ME
LE
Mid-exponential (ME)
Late-exponential (LE)
Stationary (S)

### GlnR iModulon

**Significantly overlapped TRN: GlnR**

**Biological function: Amino acid use**

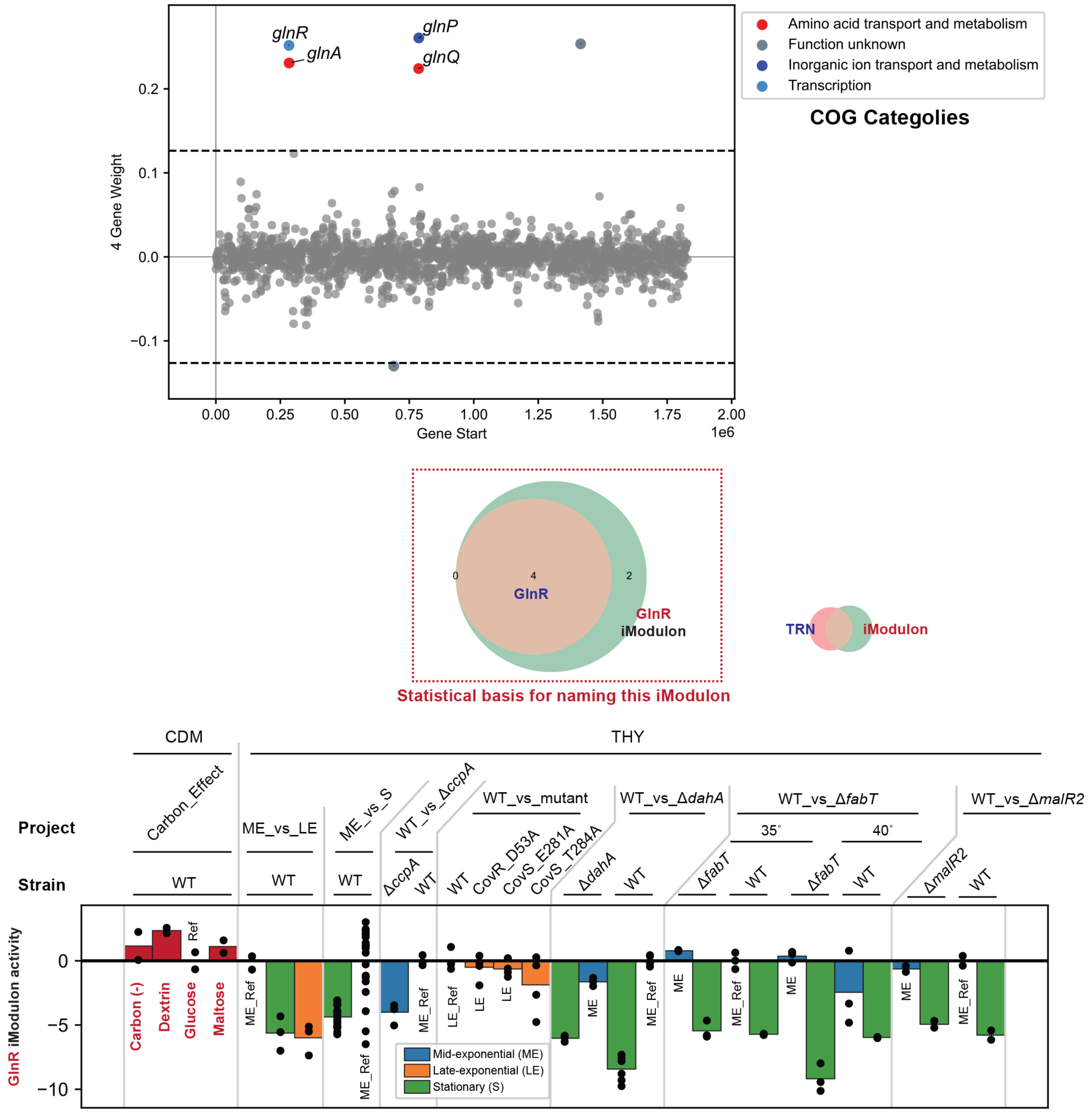

### CcpA-1 iModulon

**Significantly overlapped TRN: CcpA_core_M1 > CovR_2_LE_M1**

**Biological function: Amino acid use**

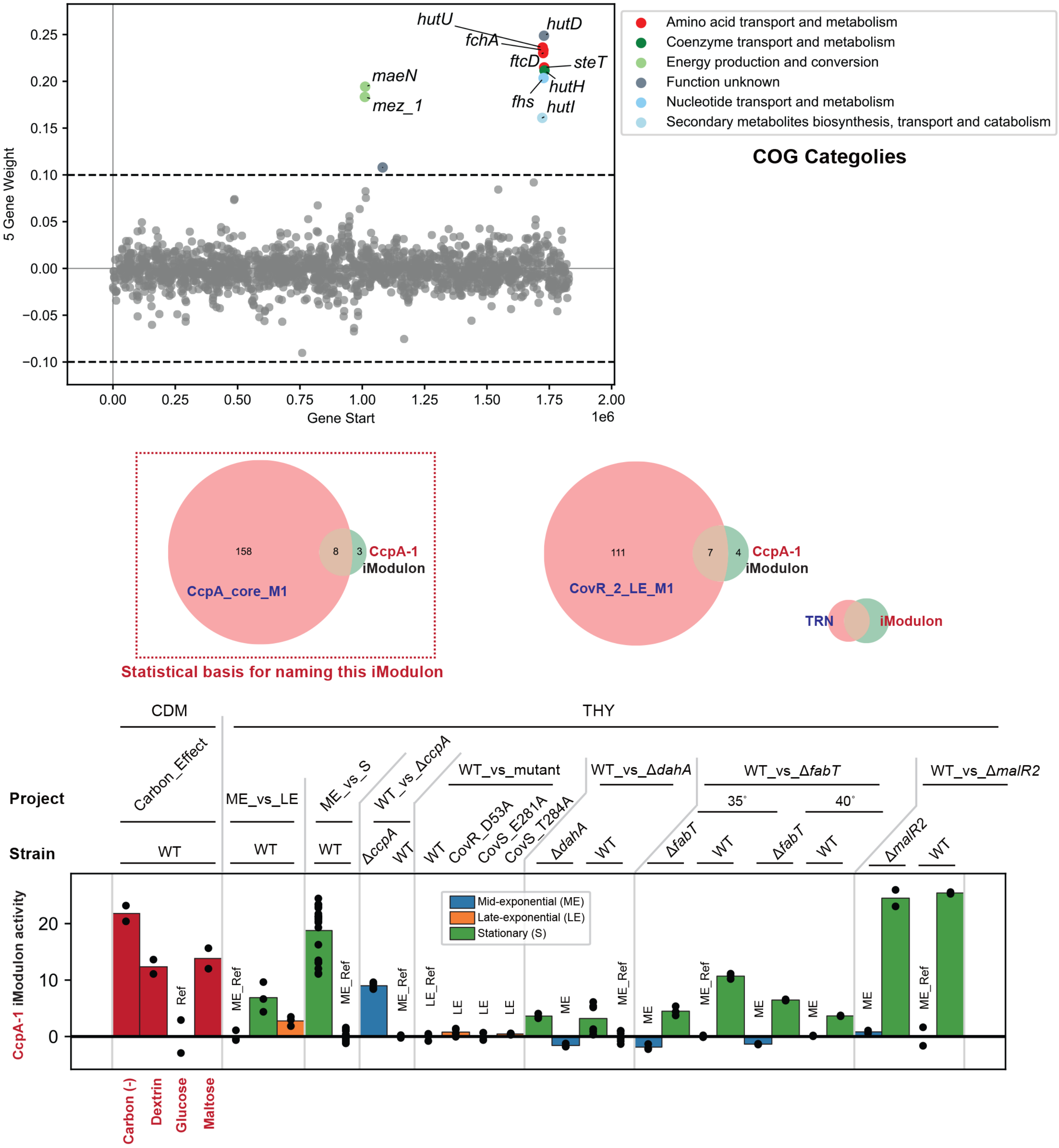

### Rgg iModulon

**Significantly overlapped TRN: Rgg_exponetional_M49 > CcpA_core_M1 > CovR_CcpA_M1**

**Biological function: Multi-purpose metabolism**

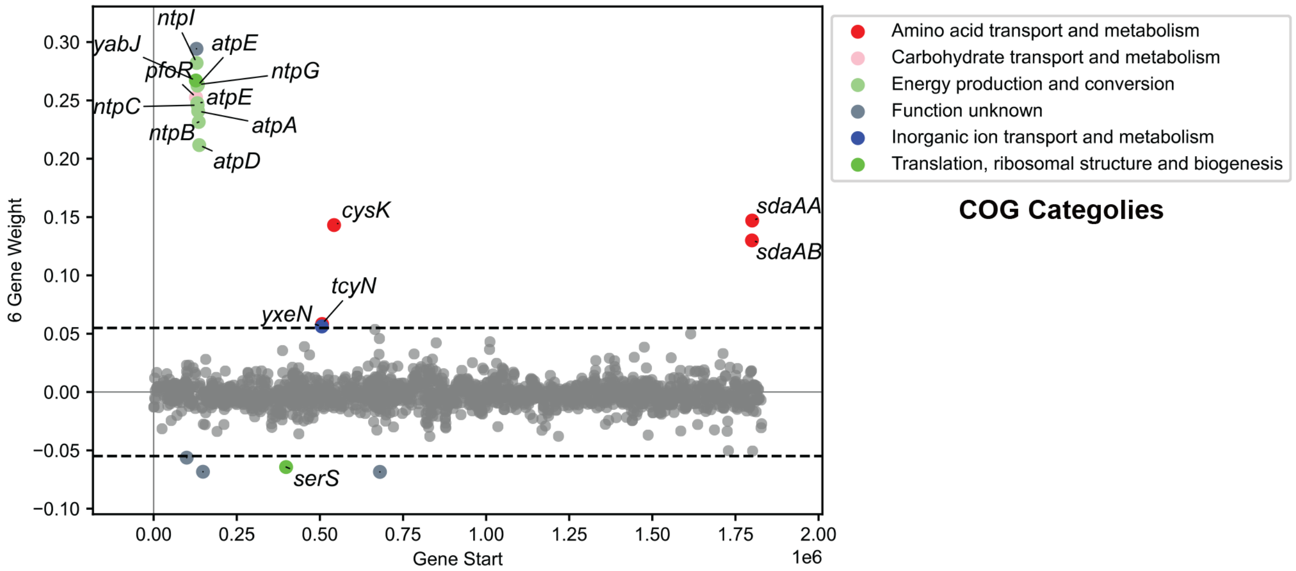


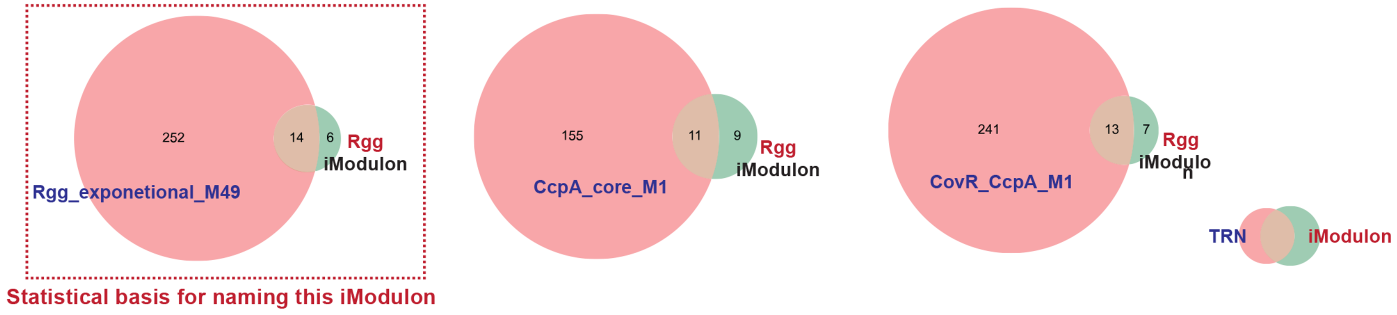


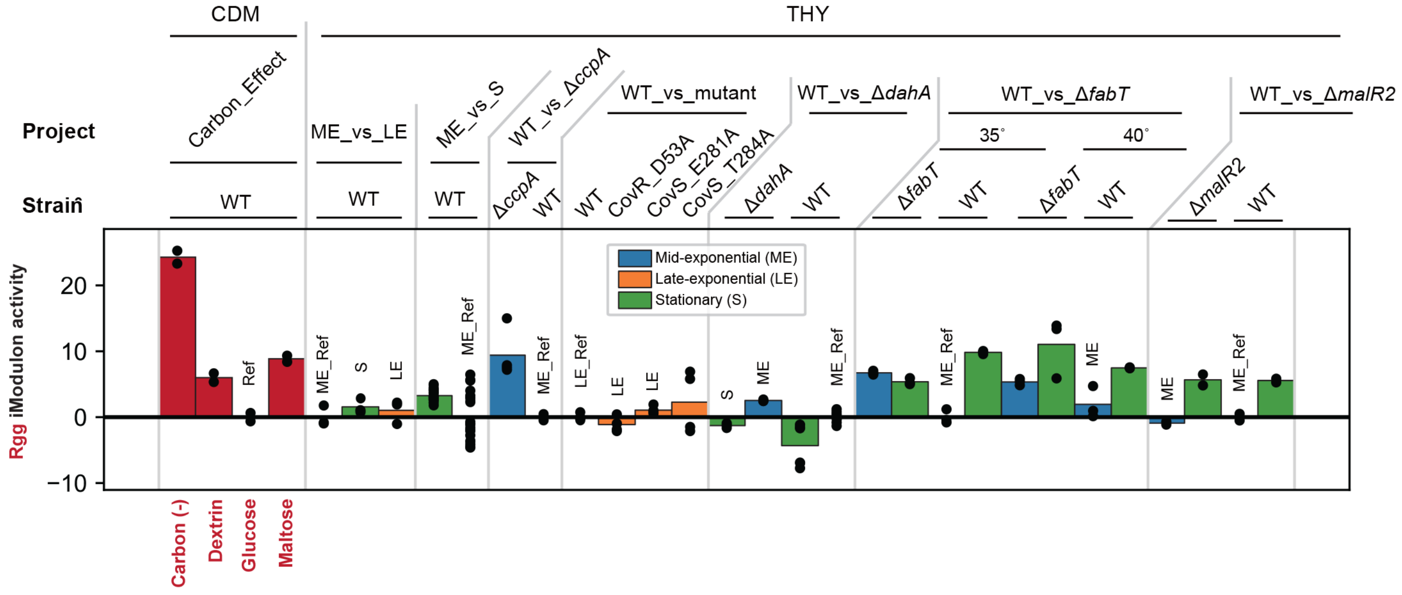

### Phage_2-1 iModulon

**Significantly overlapped TRN: Phage_2**

**Biological function: Prophages**

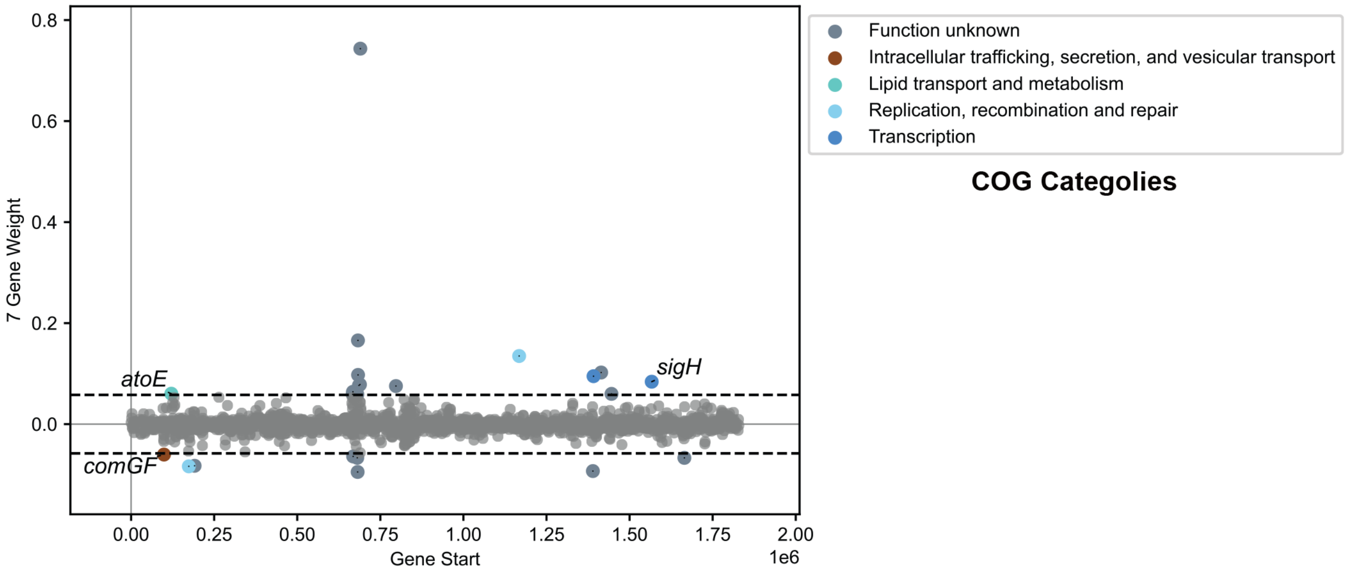


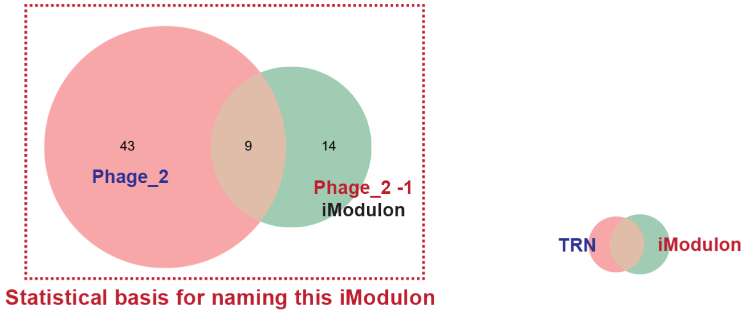


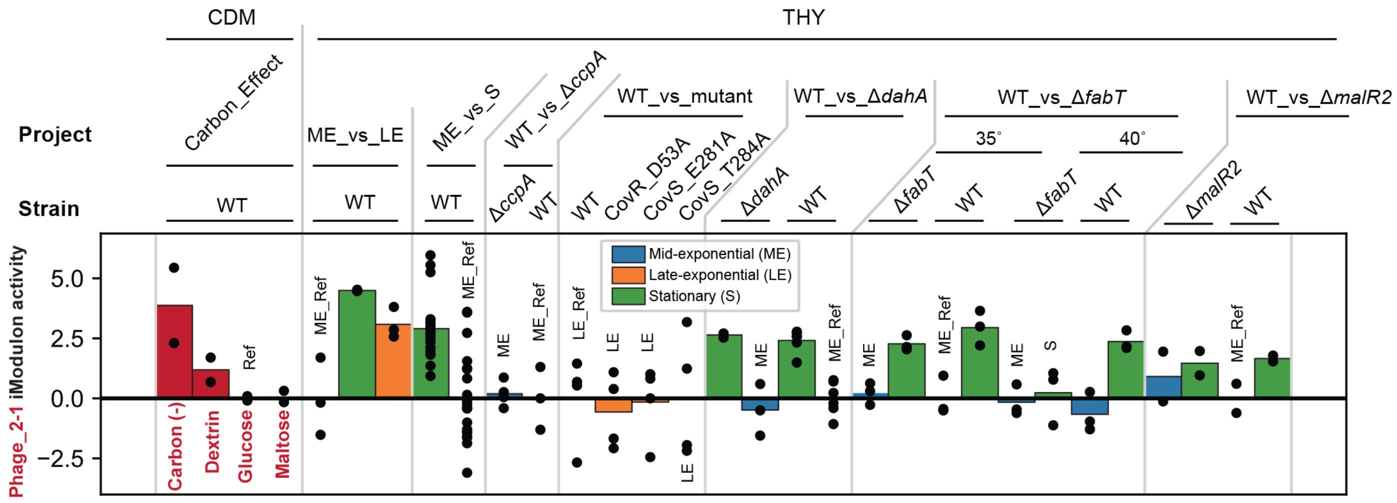

### PyrR iModulon

**Significantly overlapped TRN: PyrR > MtsR_M49 > Rgg_exponetional_M49 > CovR_1_S_M1**

**Biological function: Nucleotide use**

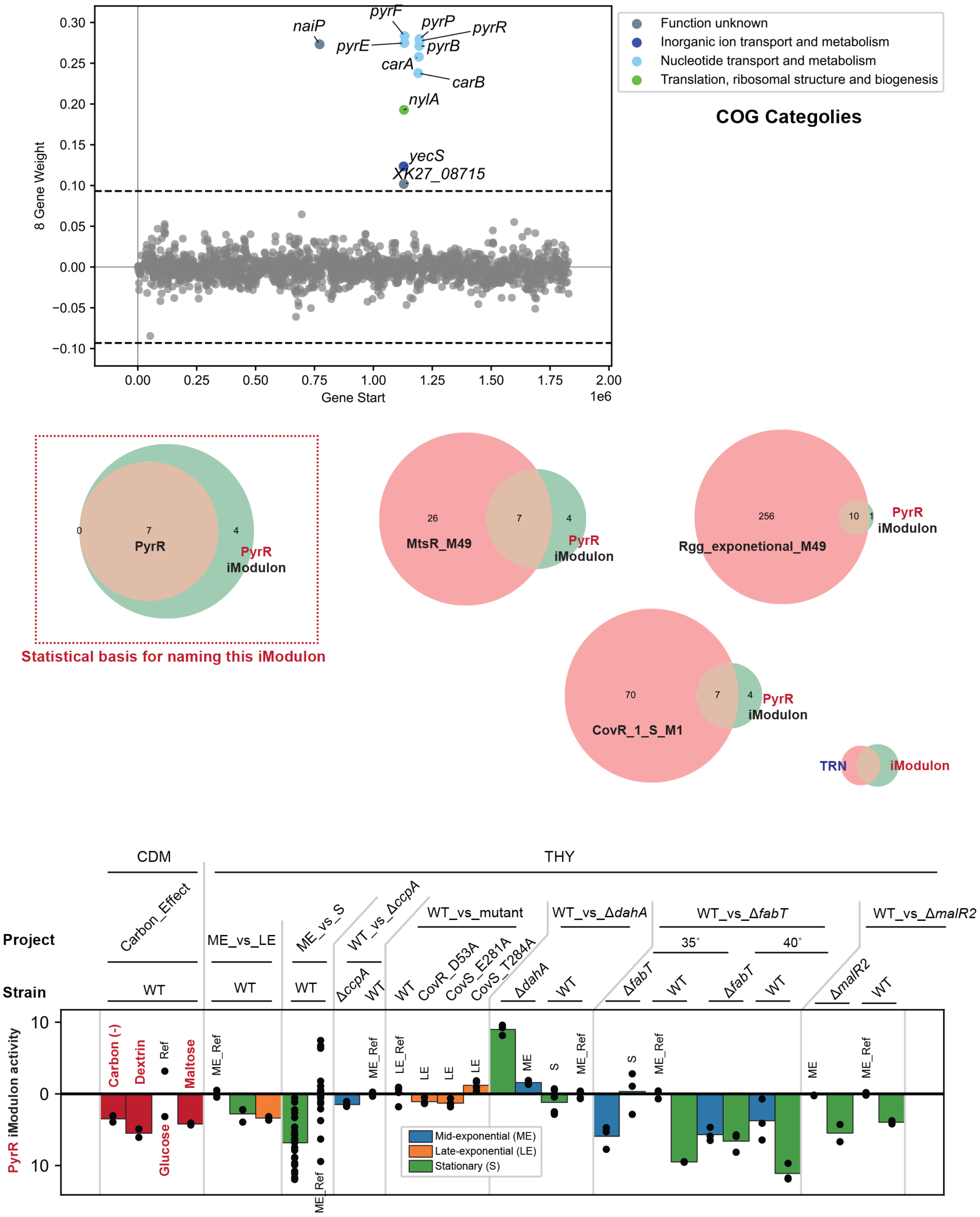

### MalR iModulon

**Significantly overlapped TRN: MalR_M1 > MalR > CcpA > FruR**

**Biological function: Carbon use**

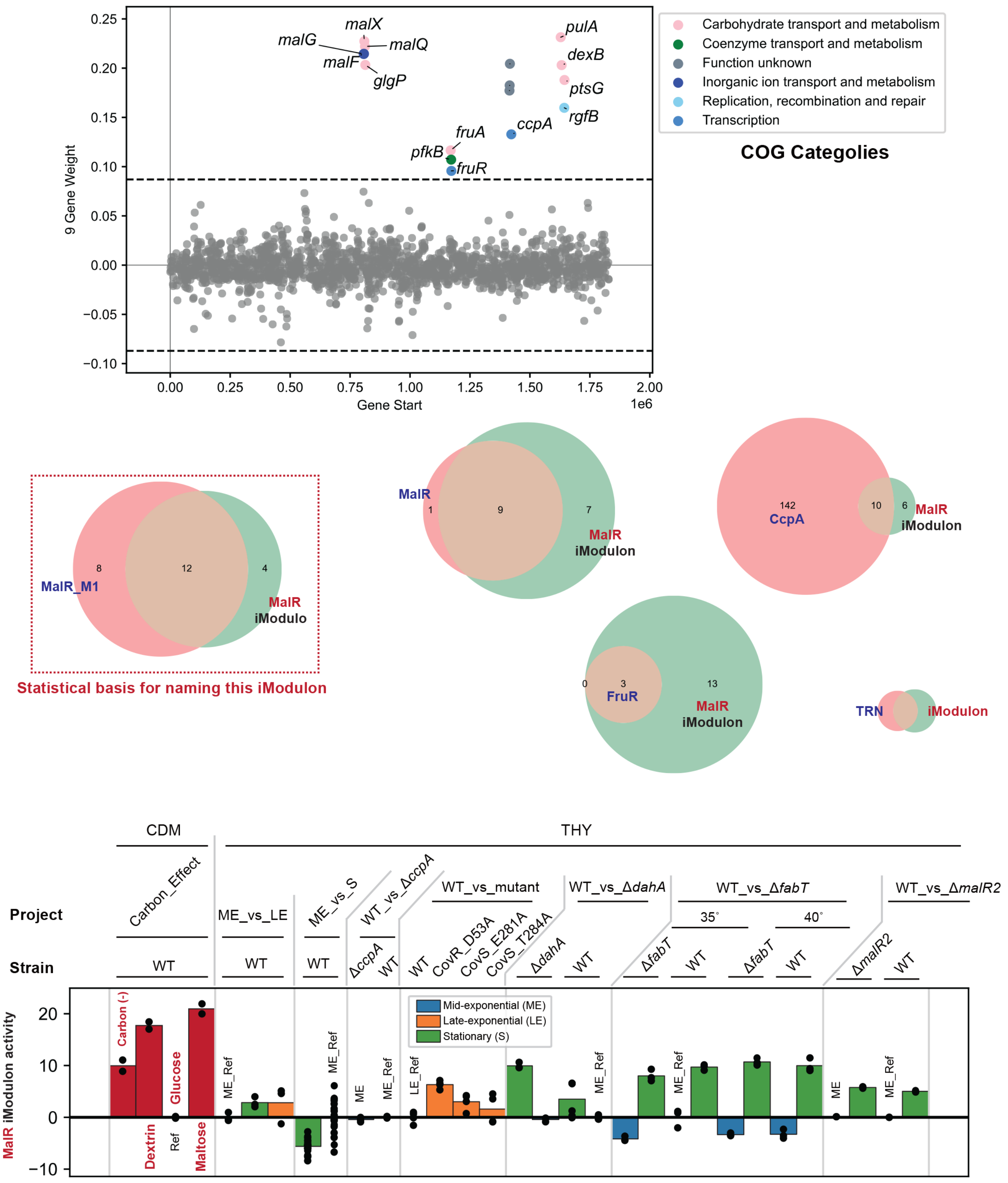

#### Unc_2 iModulon

**Significantly overlapped TRN: ---**

**Biological function: ---**

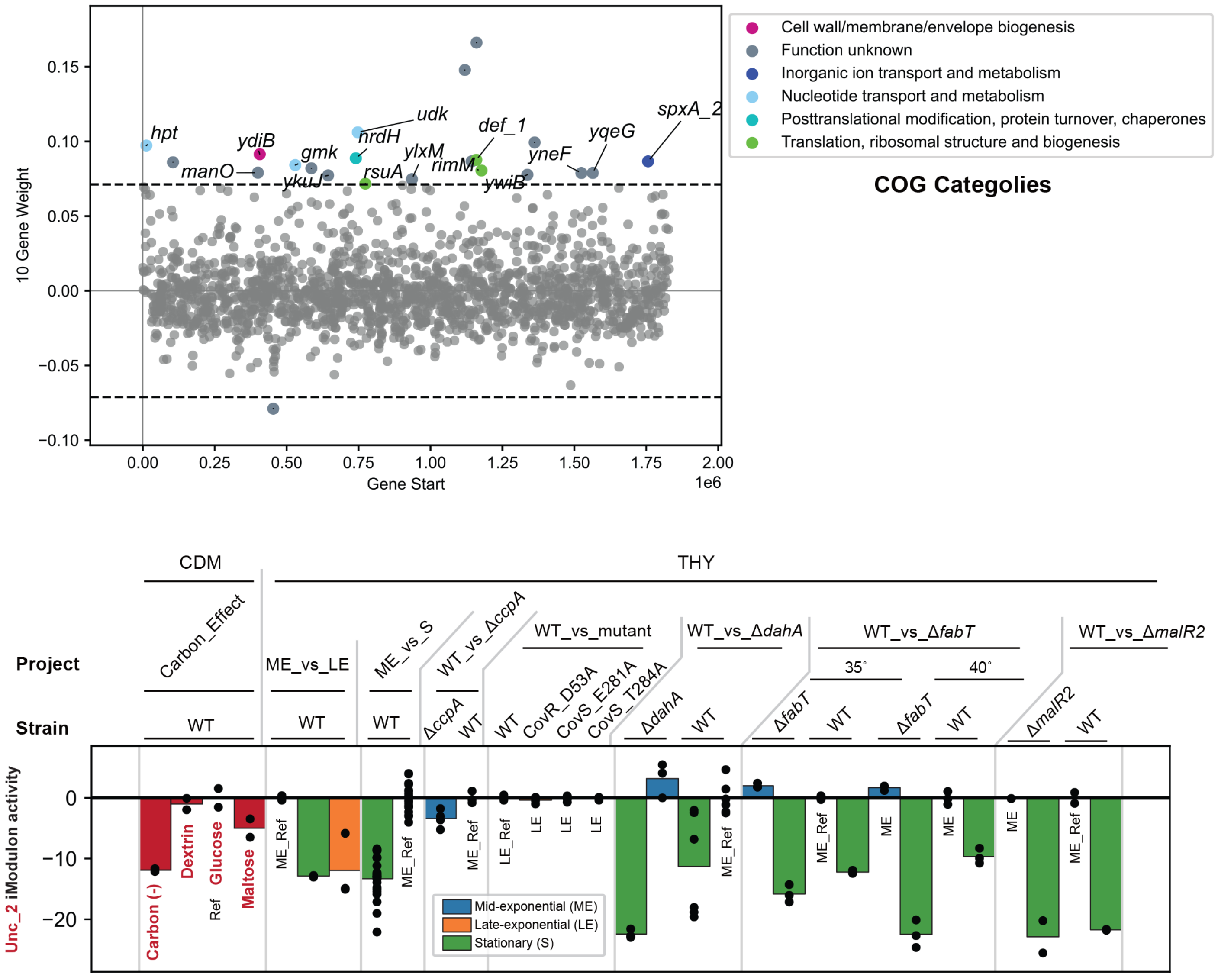

#### Unc_3 iModulon

**Significantly overlapped TRN: ---**

**Biological function: Translation**

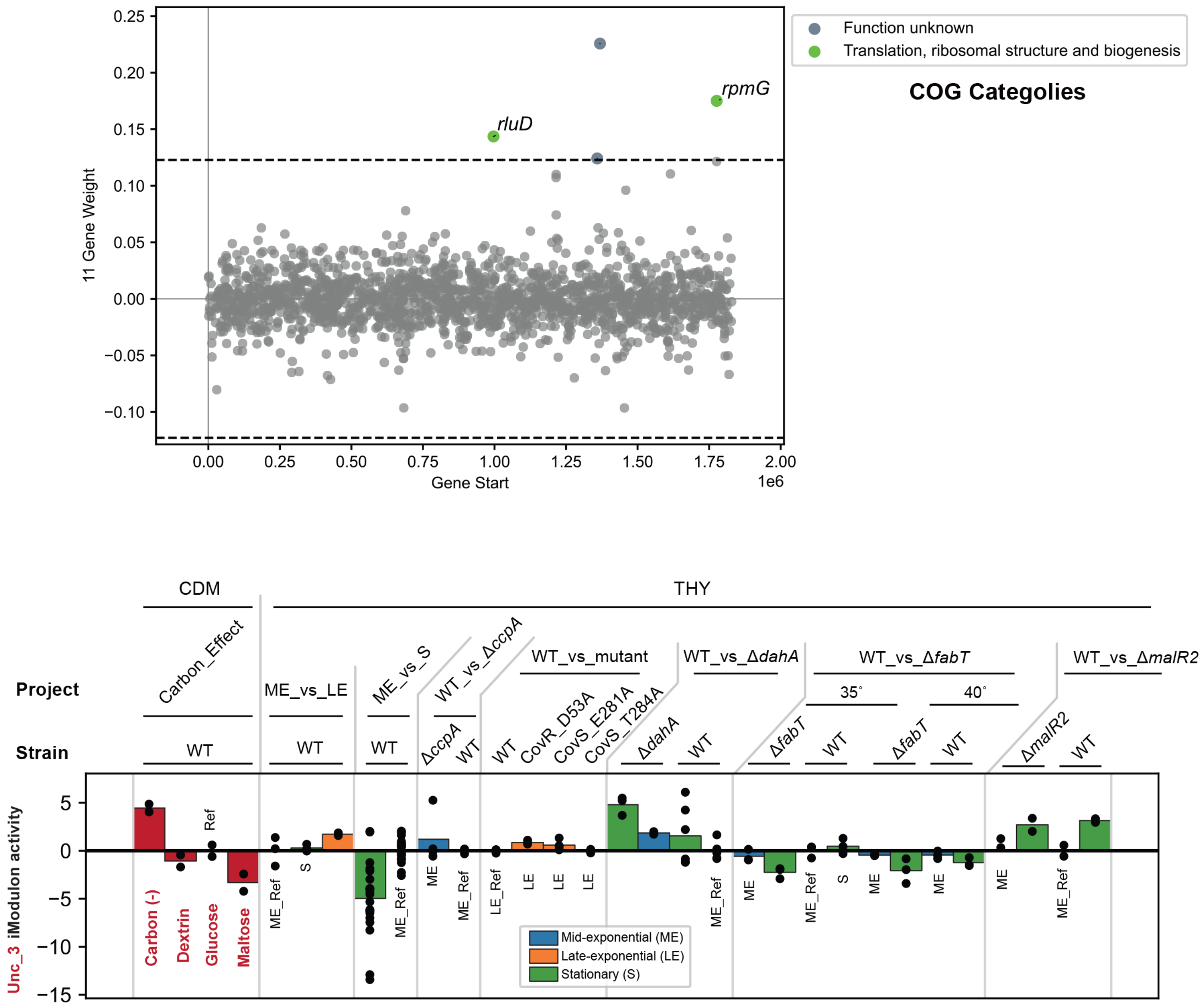

### VicR iModulon

**Significantly overlapped TRN: VicR**

**Biological function: Carbon use**

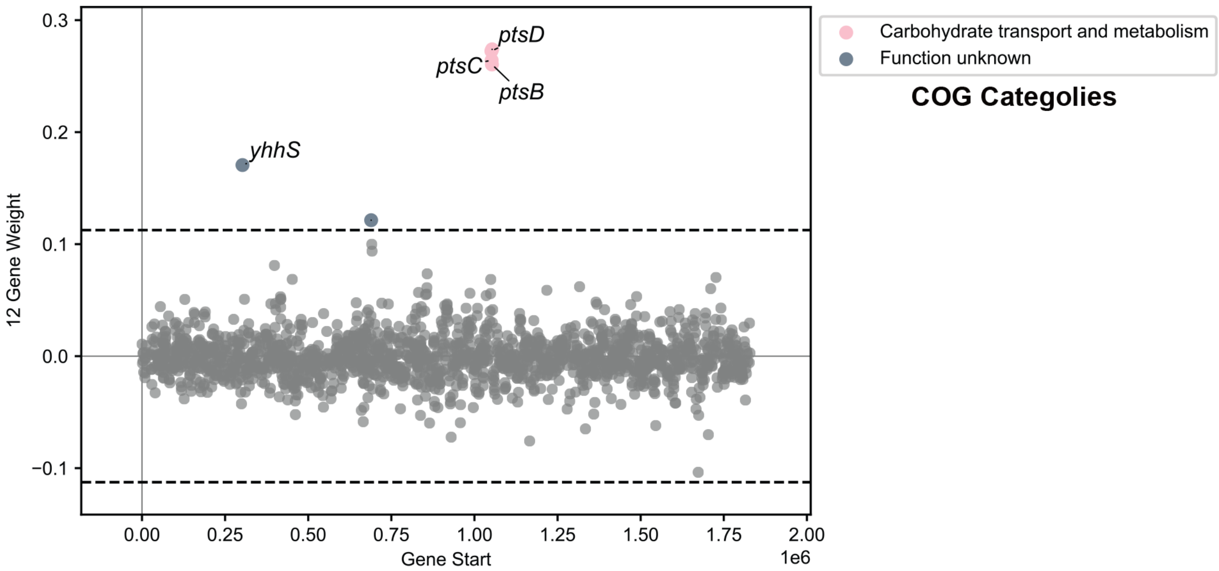


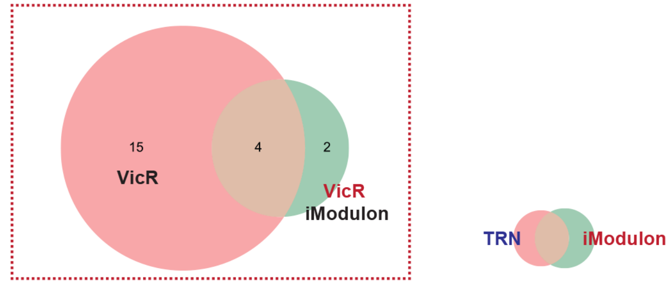


Statistical basis for naming this iModulon

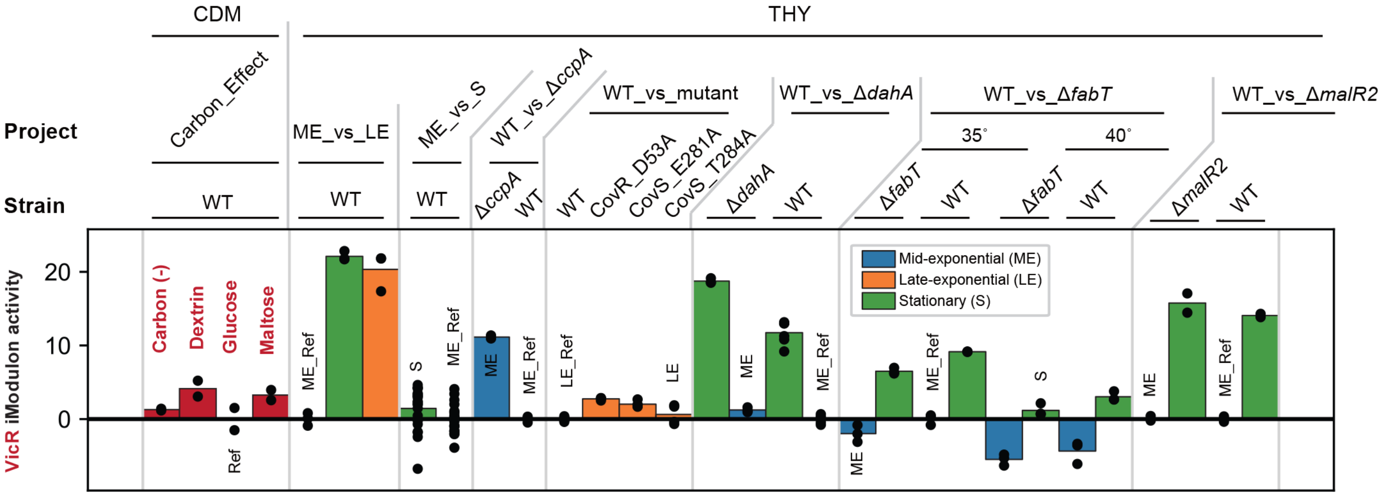

### Phage_1-2 iModulon

**Significantly overlapped TRN: Phage_1 > SPy1285**

**Biological function: Prophages**

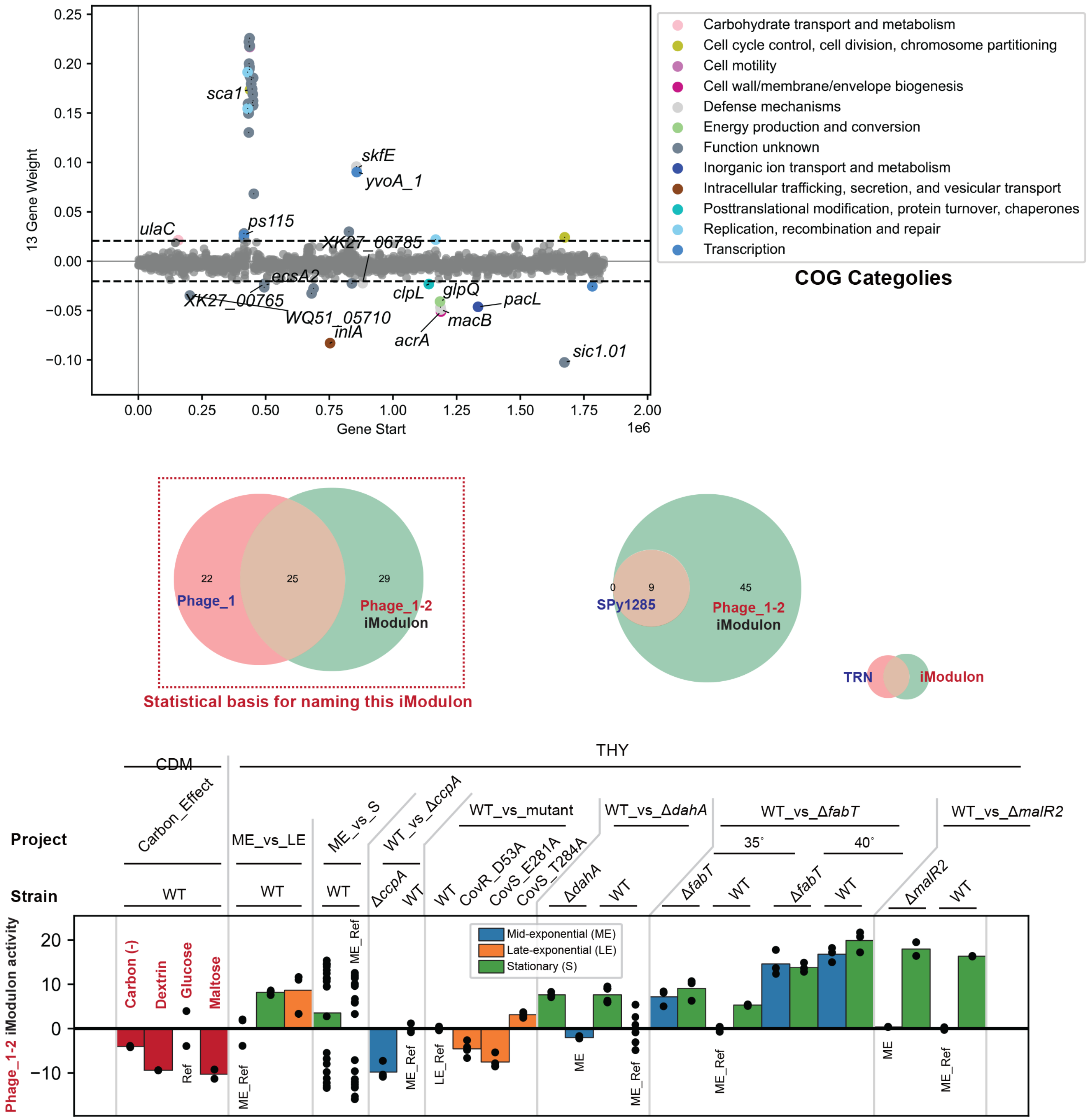

#### Unc_4 iModulon

**Significantly overlapped TRN: ---**

**Biological function: Multi-purpose metabolism**

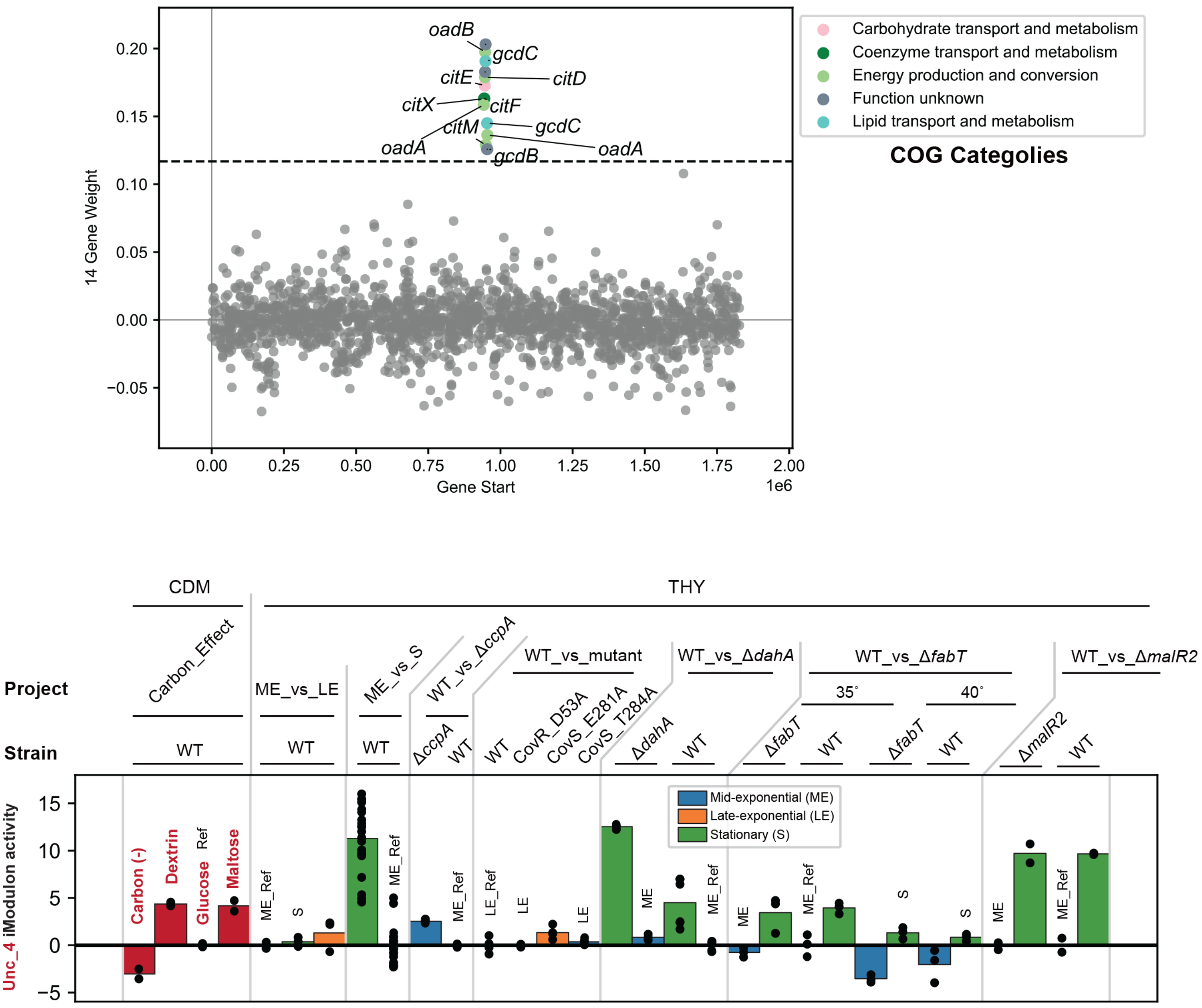

### NanR iModulon

**Significantly overlapped TRN:**

**NanR > CcpA > CcpA_core_M1> CovR_CcpA_M1 >CovR_1_S_M1 > Rgg_exponetional_M49**

**Biological function: Carbon use**

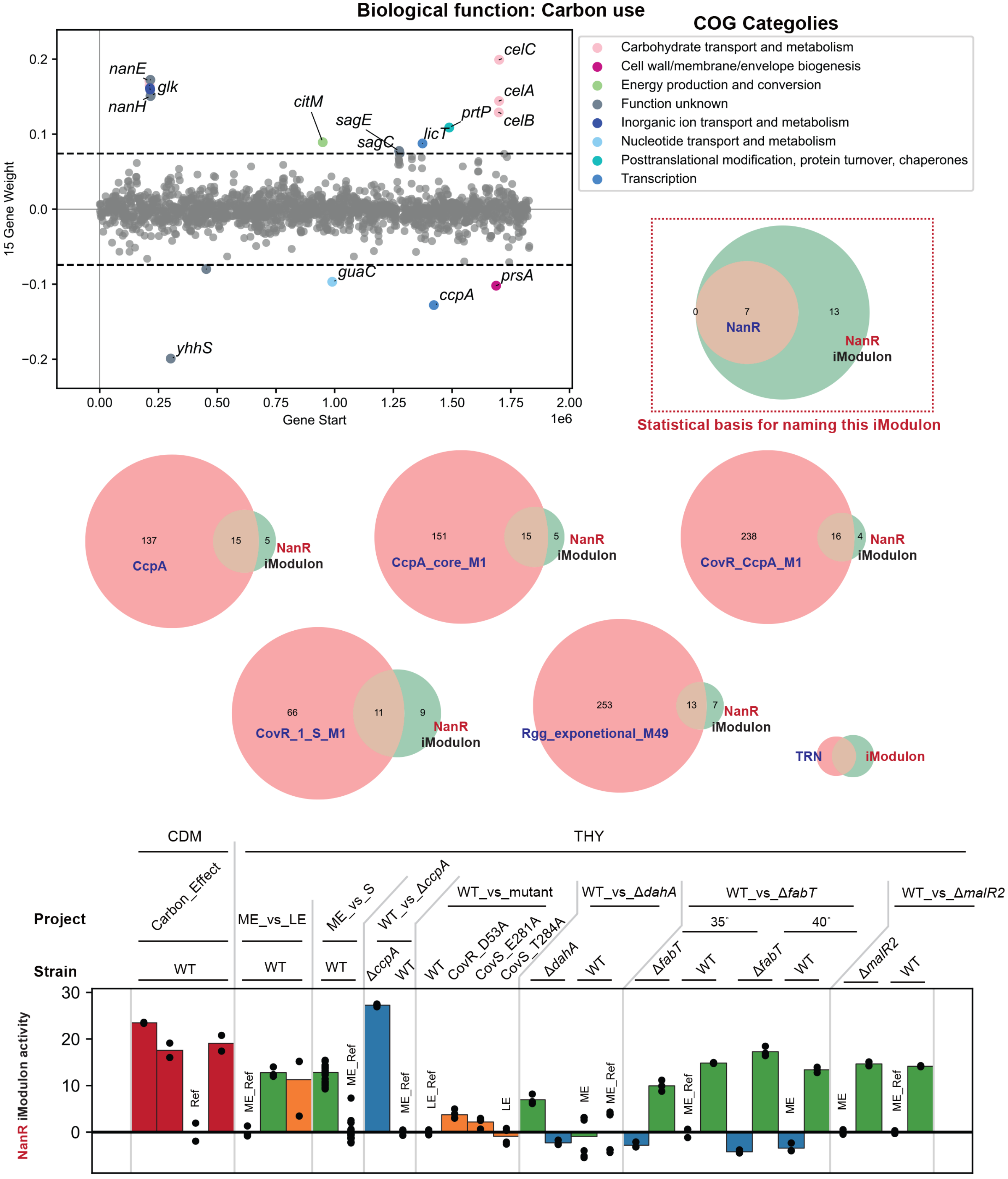

#### NrdR iModulon

##### Significantly overlapped TRN: NrdR

##### Biological function: Carbon use

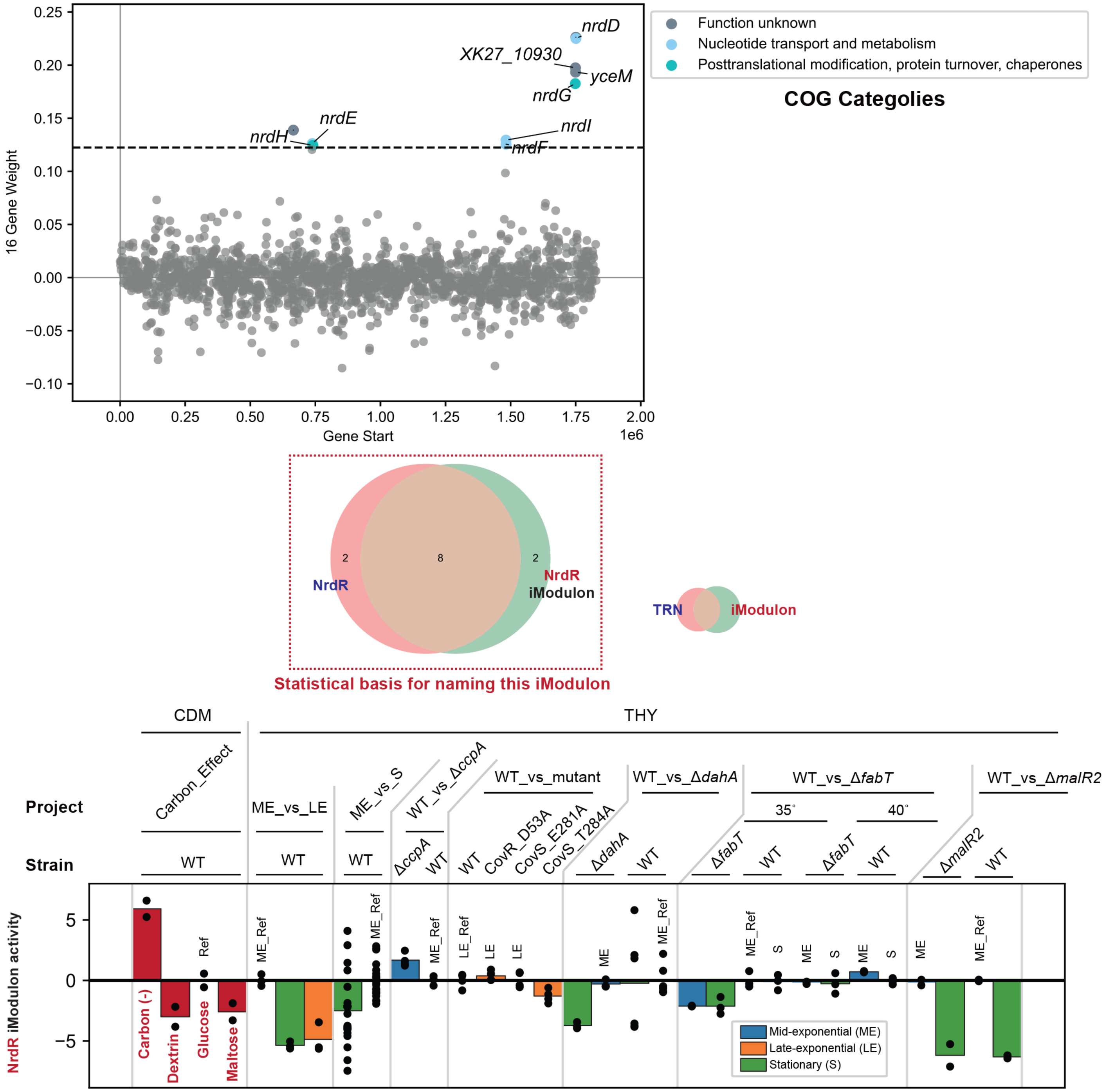

### Unc_5 iModulon

**Significantly overlapped TRN: ---**

**Biological function: ---**

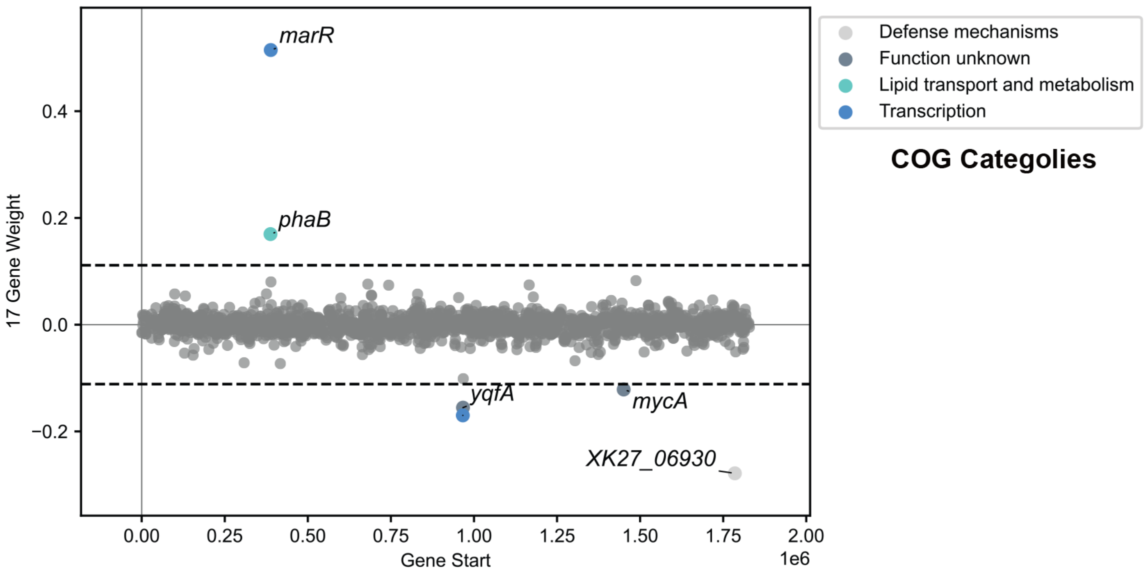


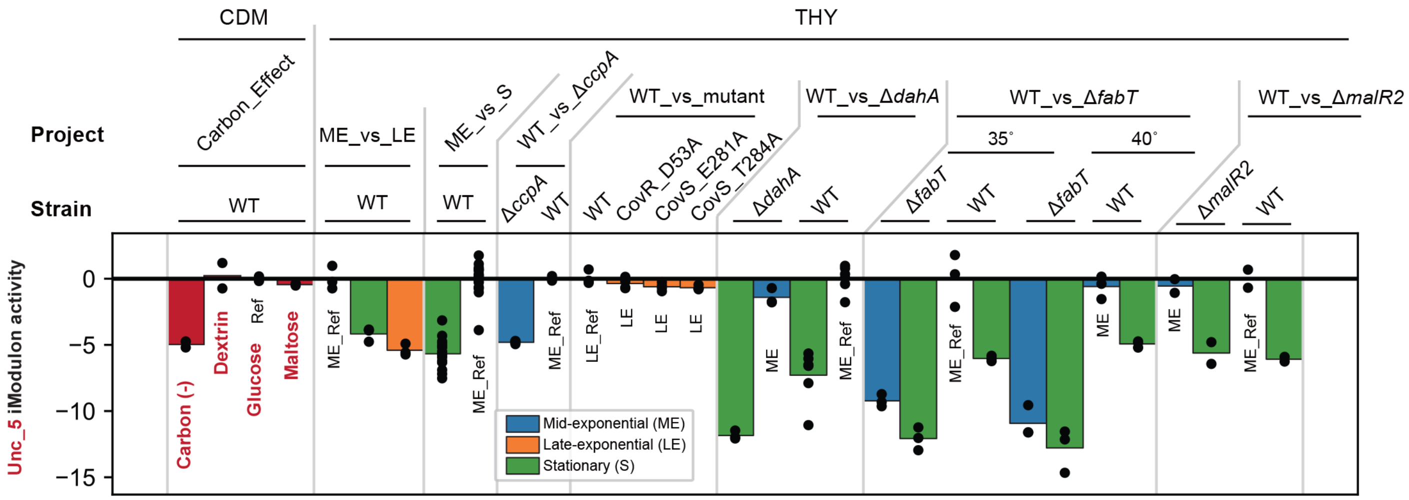

### PurR iModulon

**Significantly overlapped TRN: PurR**

**Biological function: Nucleotides**

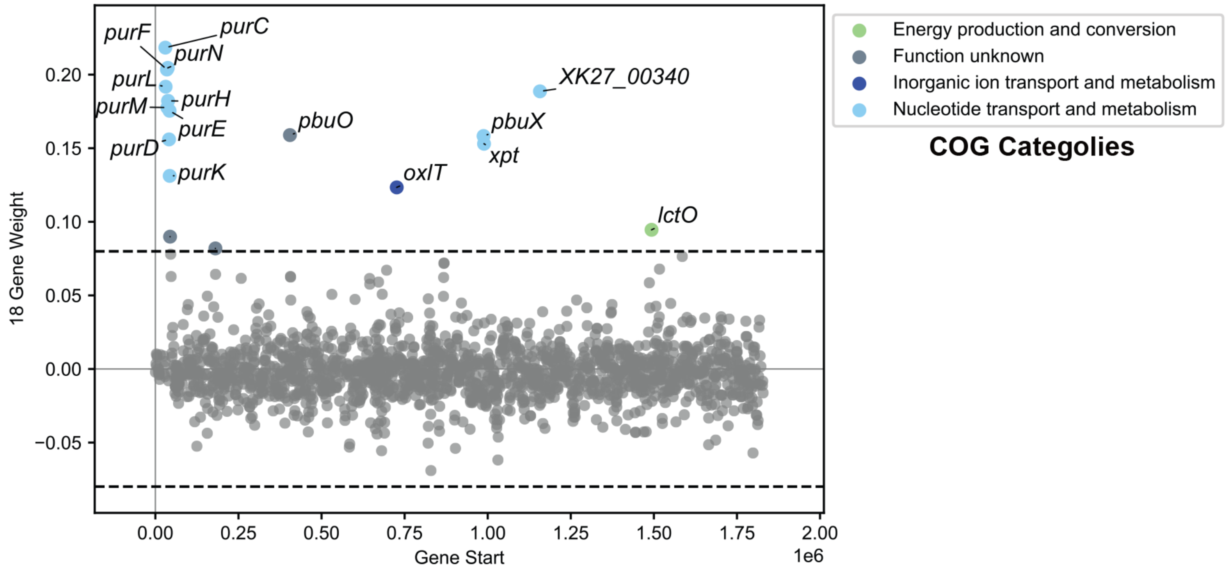


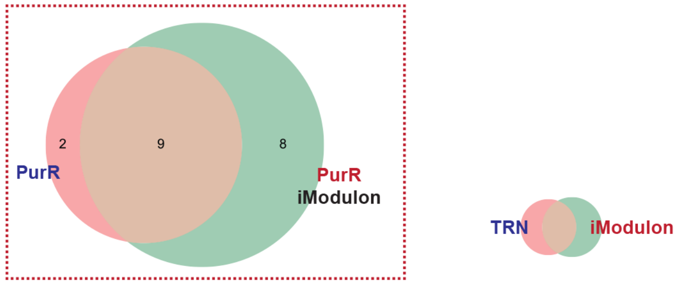


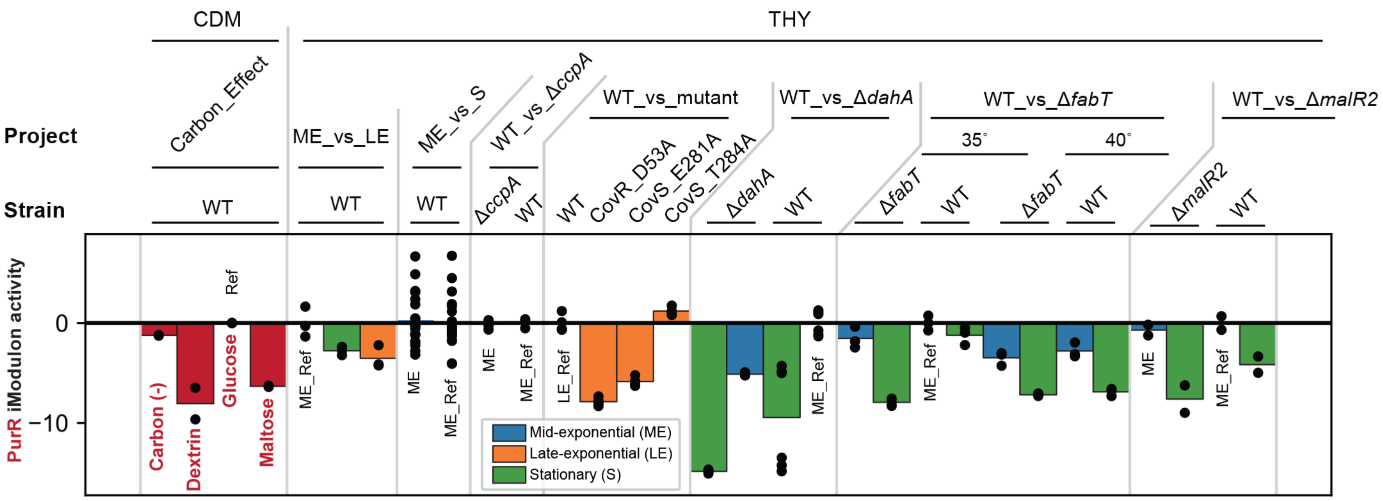

**Unc_6 iModulon**

**Significantly overlapped TRN: ---**

**Biological function: General stress**

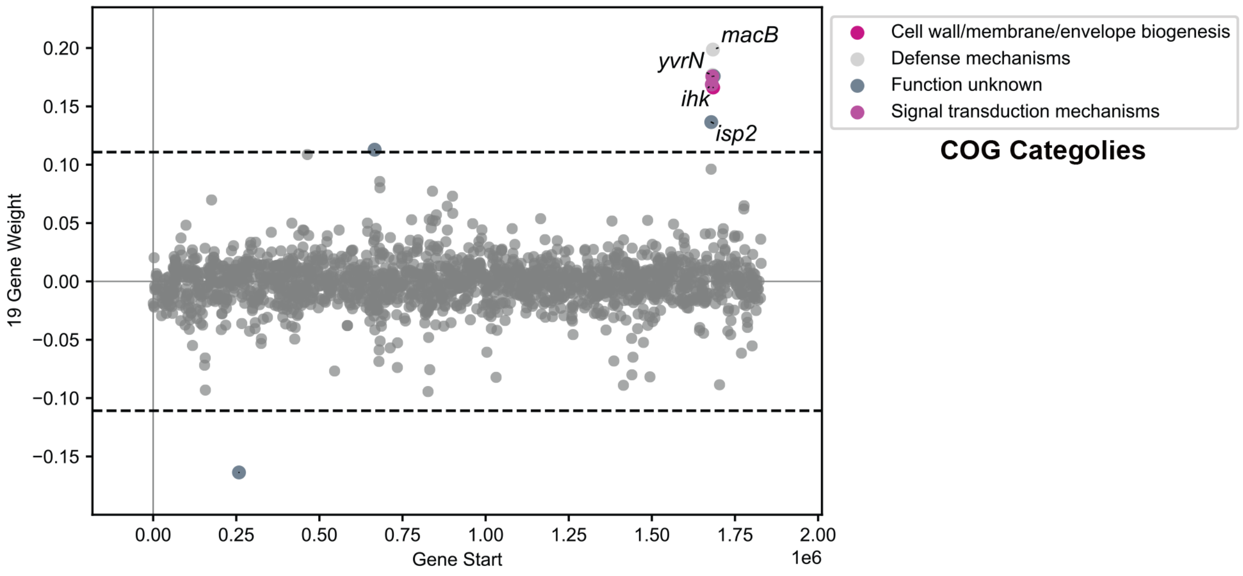


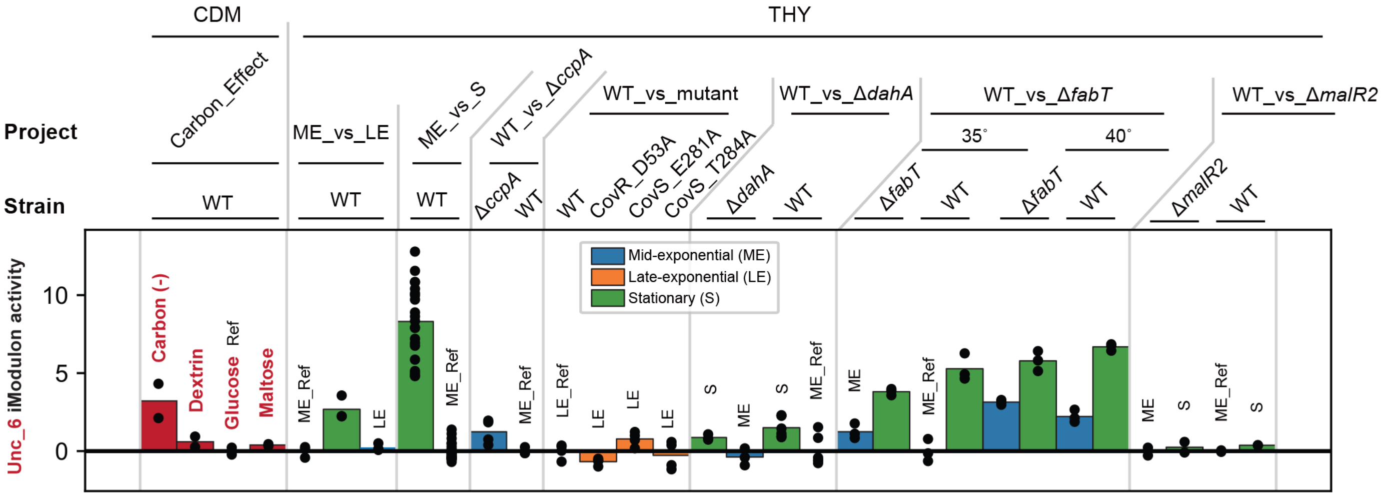

#### Unc_7 iModulon

**Significantly overlapped TRN: ---**

**Biological function: Translation**

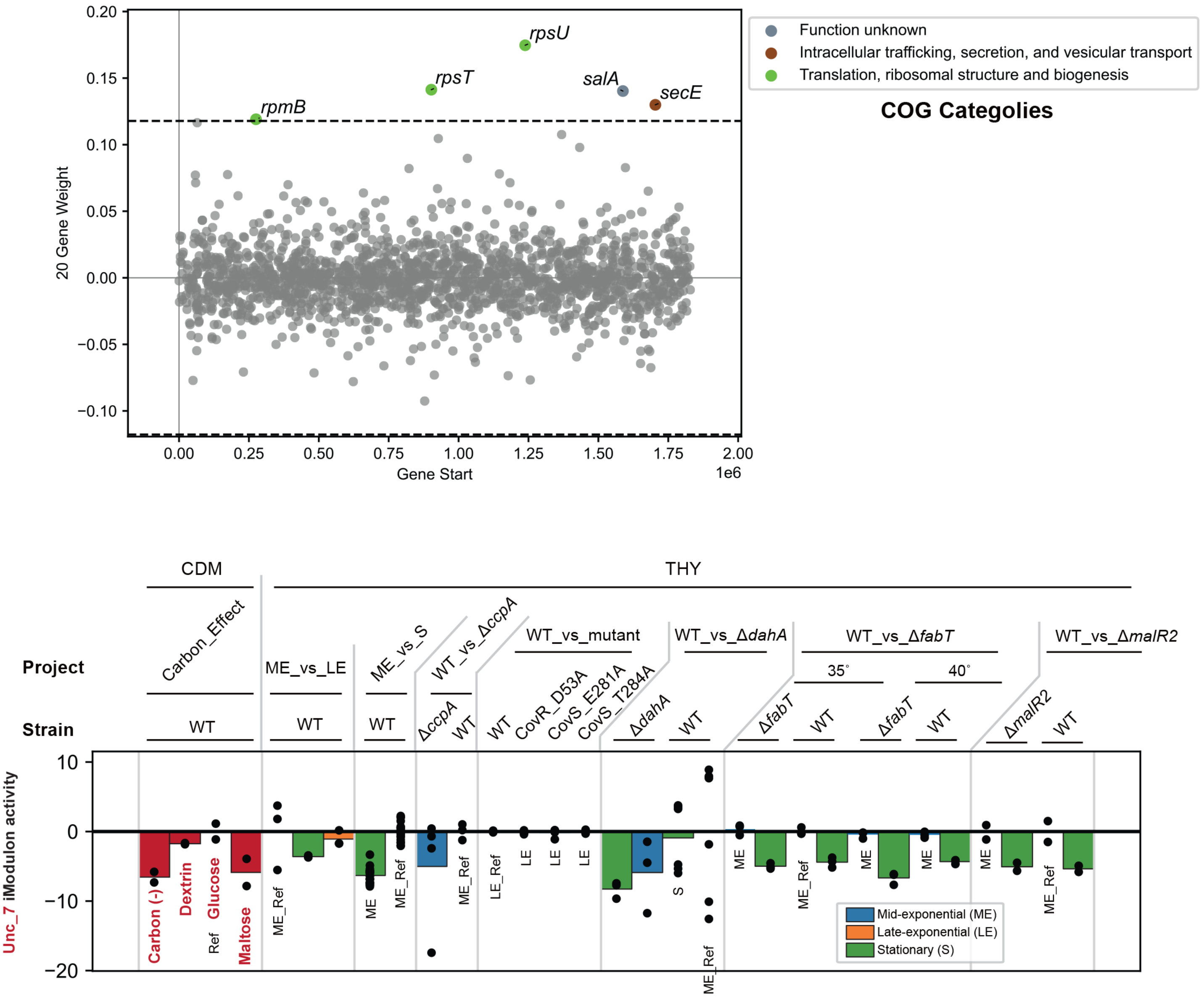

### CcpA-2 iModulon

**Significantly overlapped TRN: CcpA_core_M1 > CovR_CcpA_M1 > LacD.1_M14**

**Biological function:**

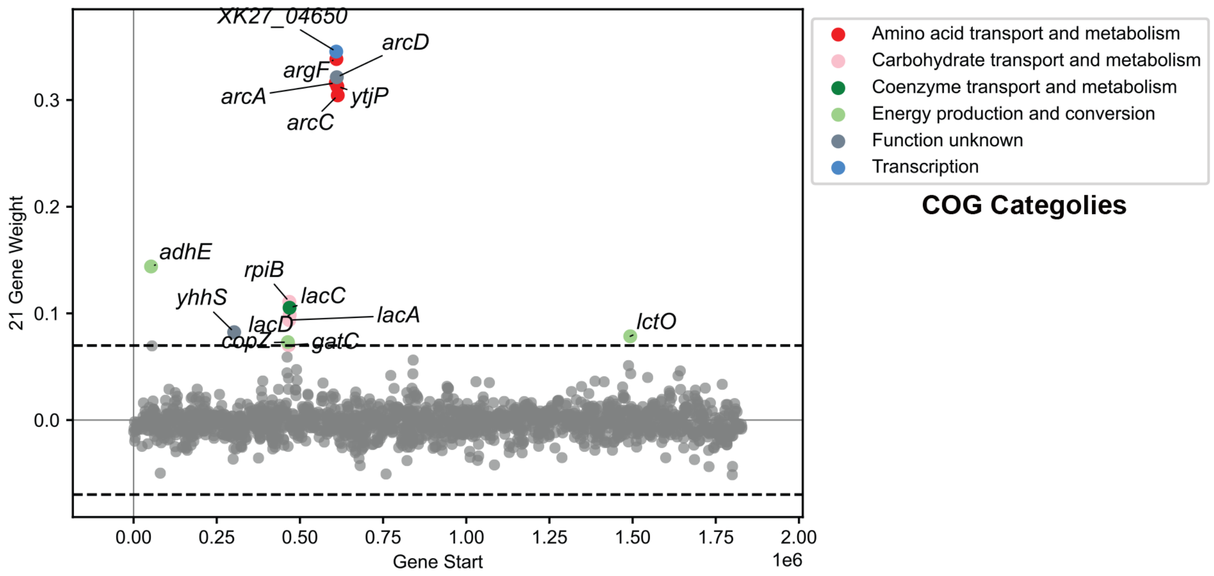


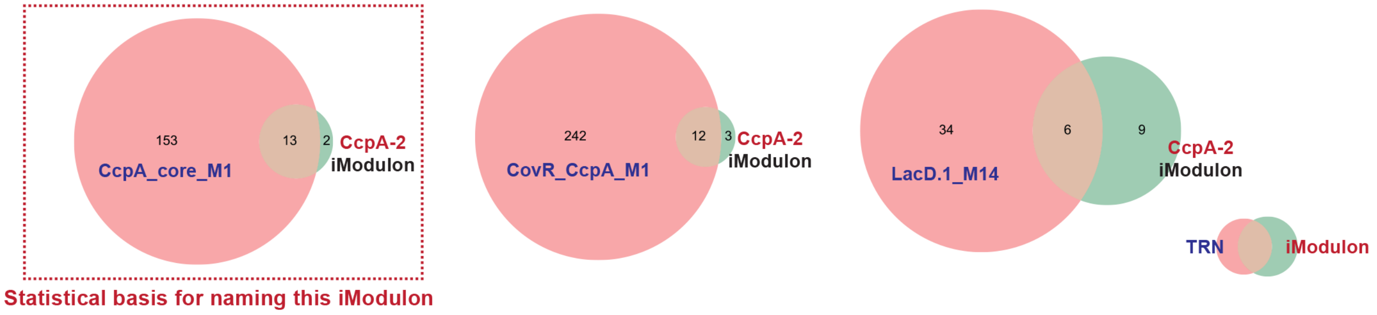


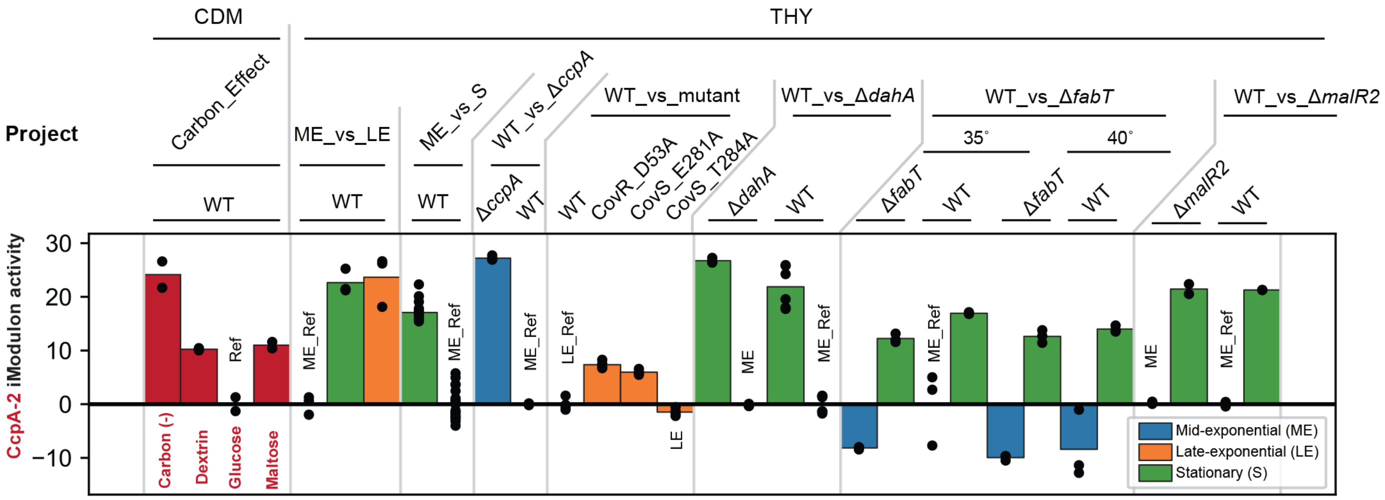

### Phage_2-2 iModulon

#### Significantly overlapped TRN: Phage_2

#### Biological function: Prophages

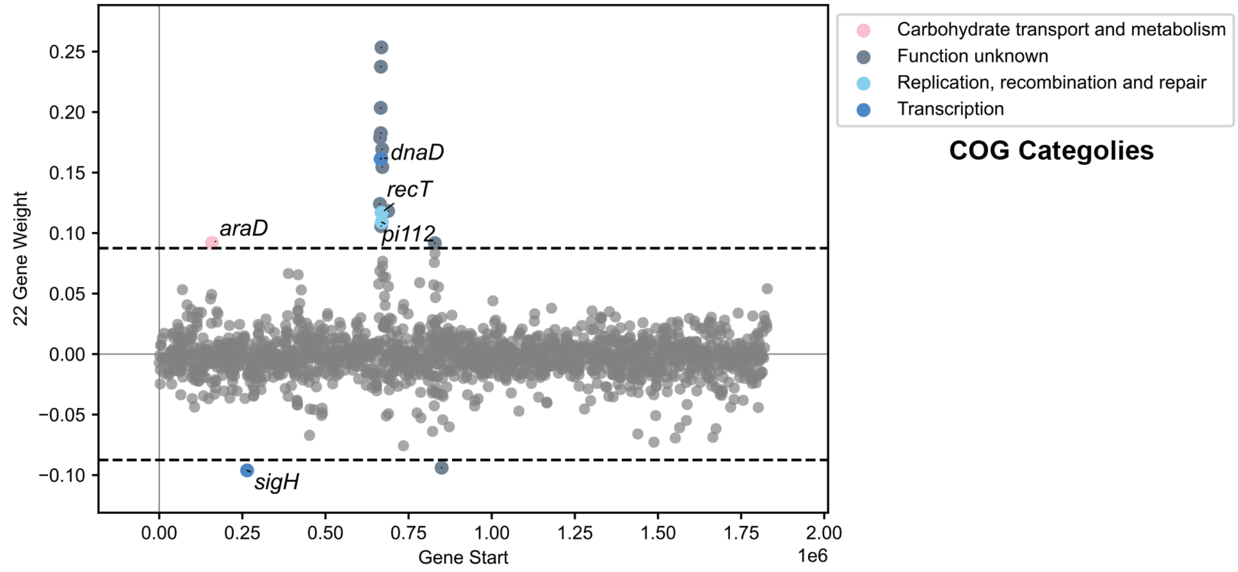


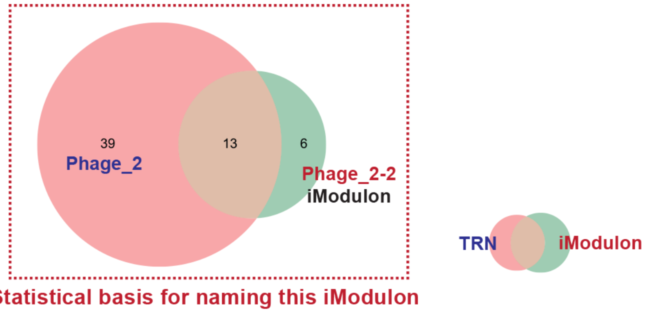


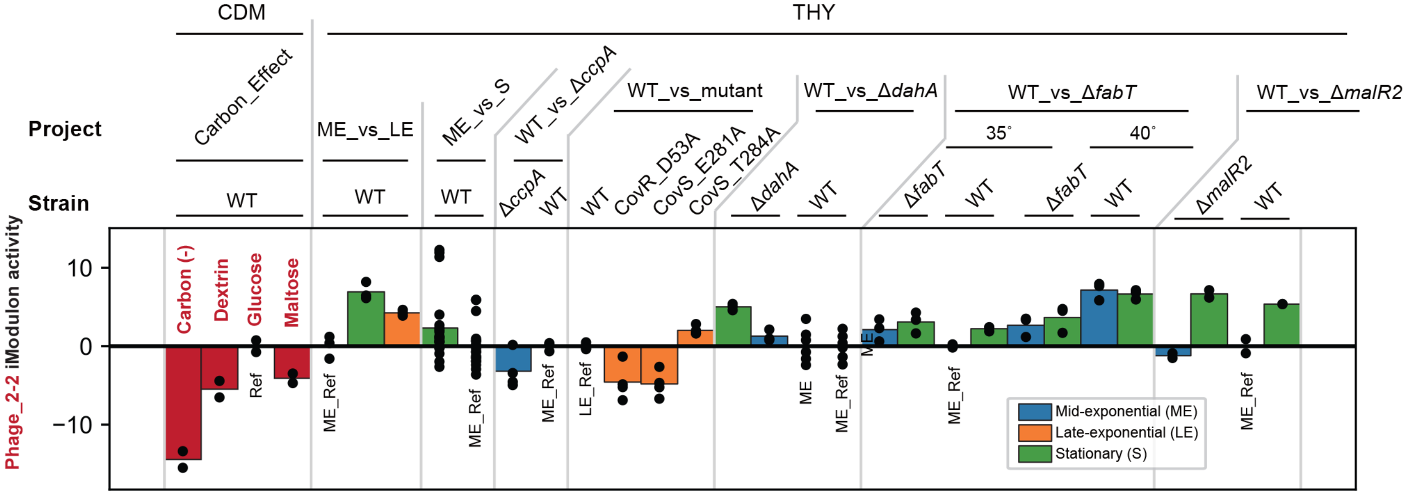

### SLS iModulon

**Significantly overlapped TRN:**
**CovR_1_S_M1 > CovR_1_ME_M1 > CovR_CcpA_M1 > CcpA_core_M1 > Rgg_exponetional_M49 > CovR_2_LE_M1 > CcpA > Mga_low_Glucose_M1 > CovR_2_S_M1 > Rgg_postexponential_M49**

**Biological function: Toxin production**

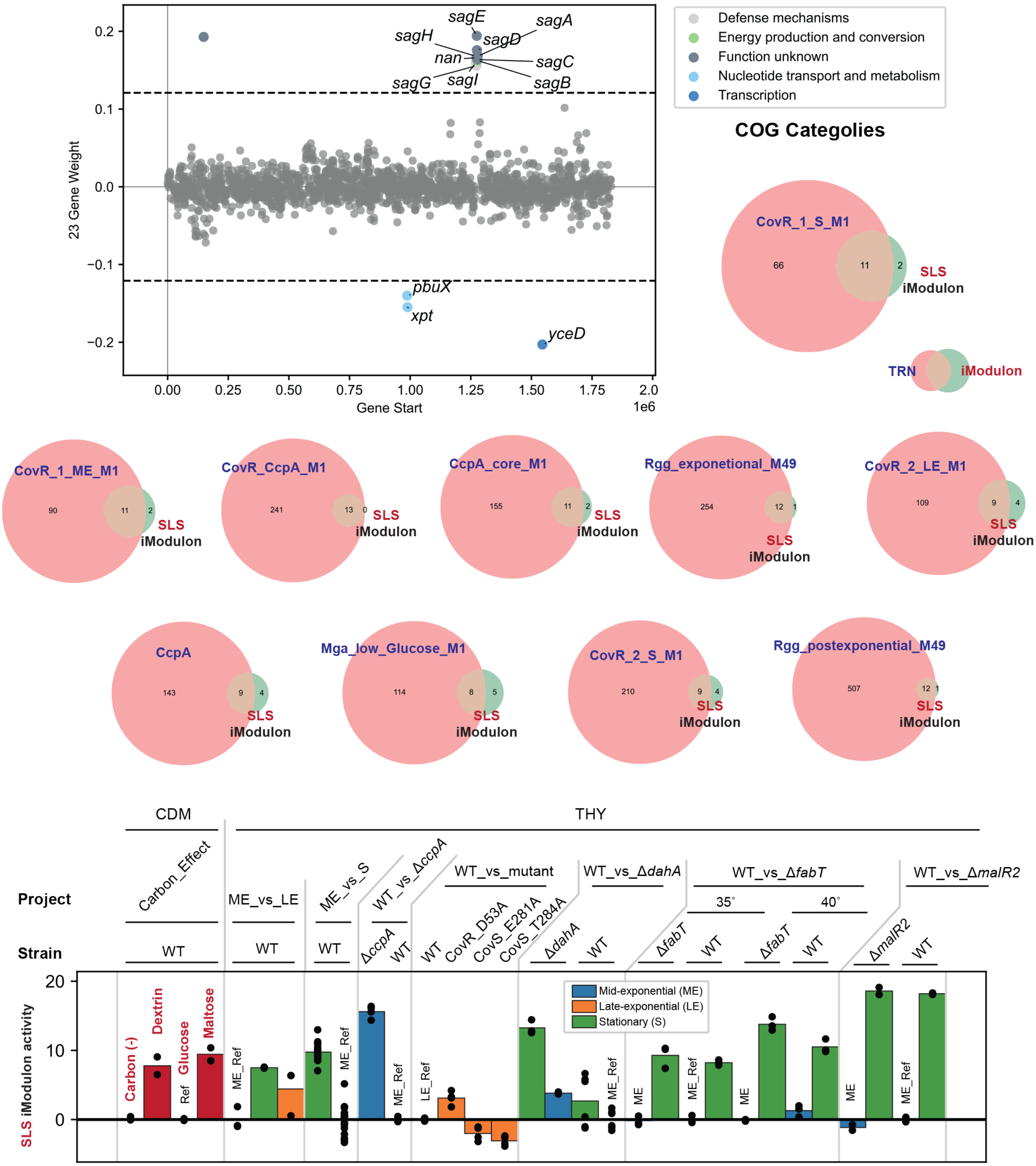

#### Unc_8 iModulon

**Significantly overlapped TRN: ---**

**Biological function: Translation**

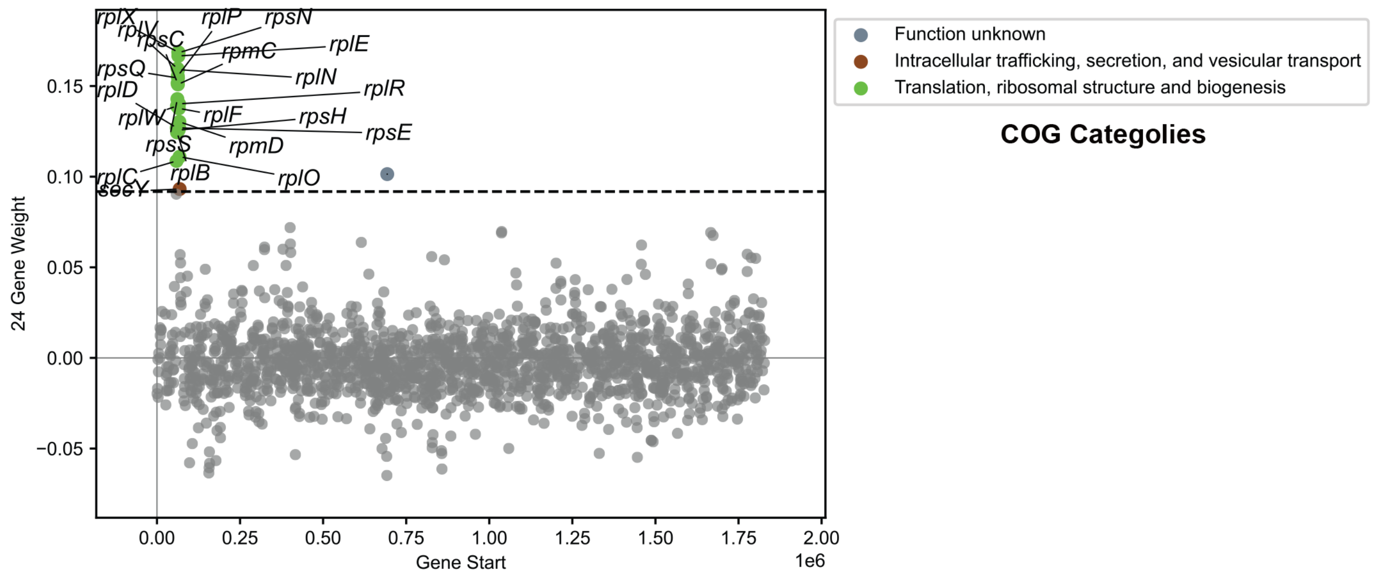


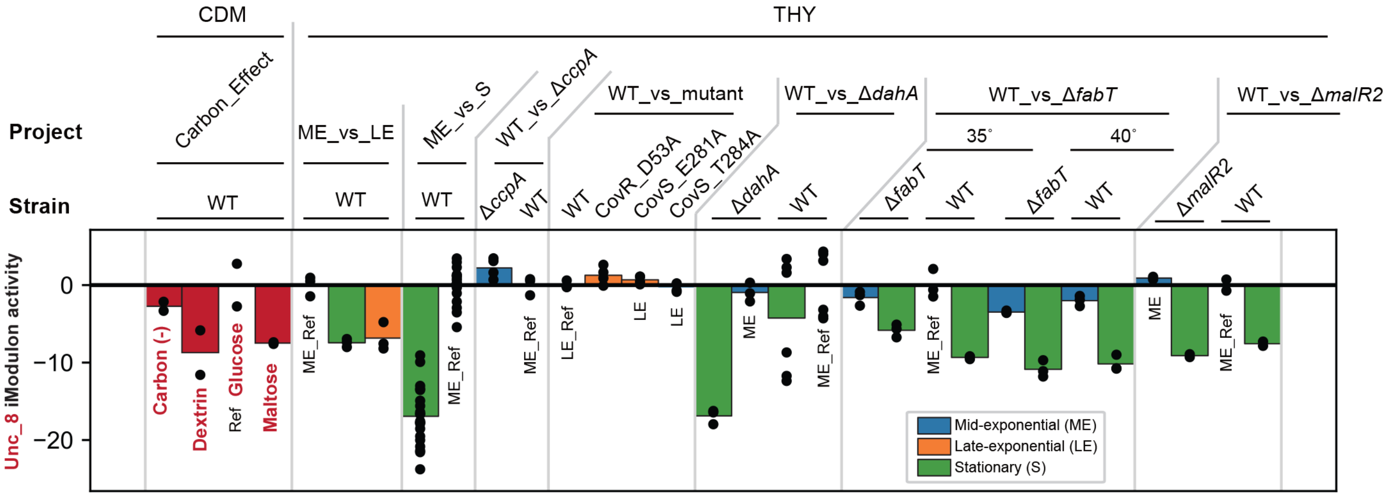

#### Unc_9 iModulon

**Significantly overlapped TRN:**

**Biological function:**

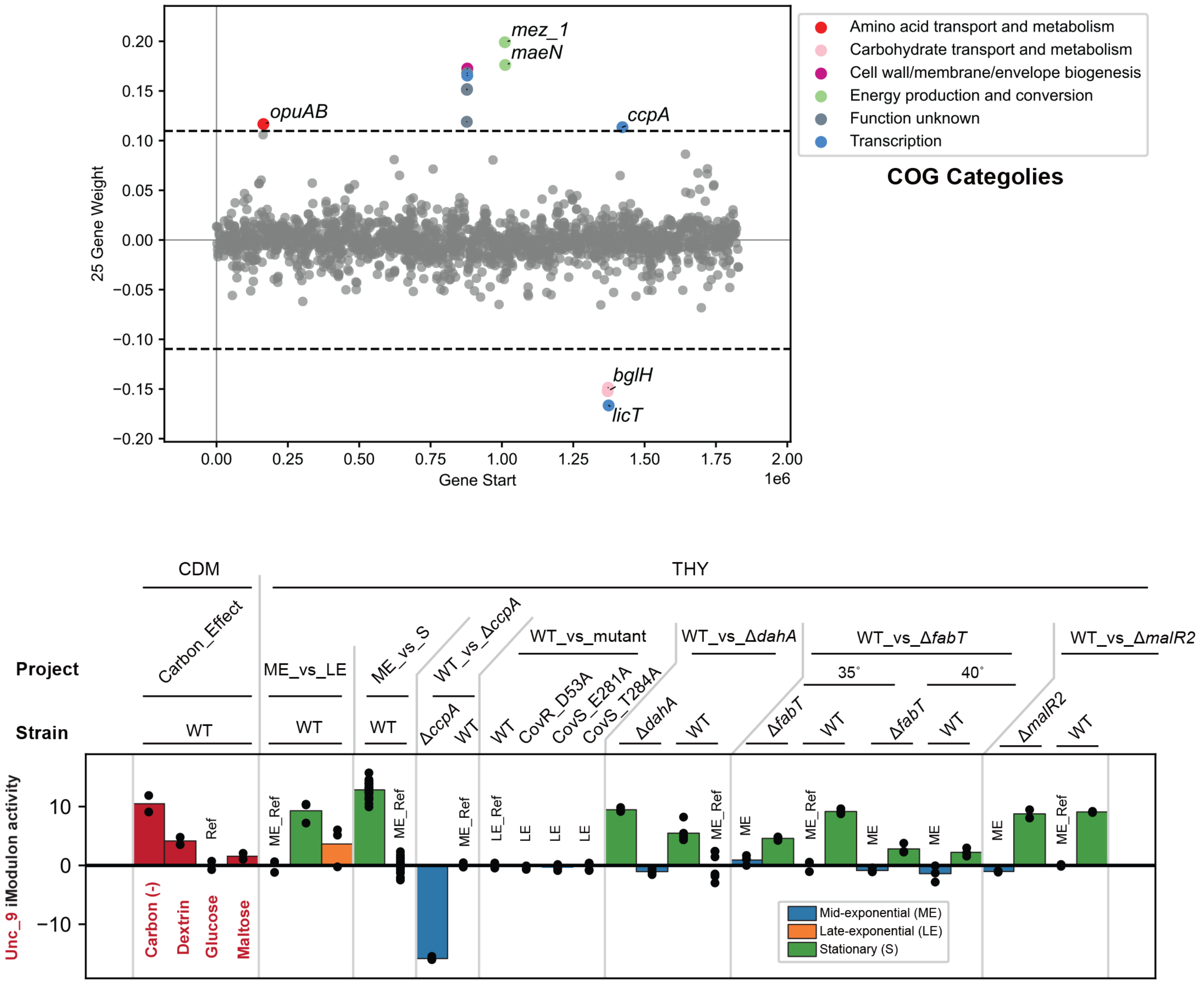

### Mga iModulon

**Significantly overlapped TRN: Mga_high_Glucose_M1**

**Biological function: Carbon use**

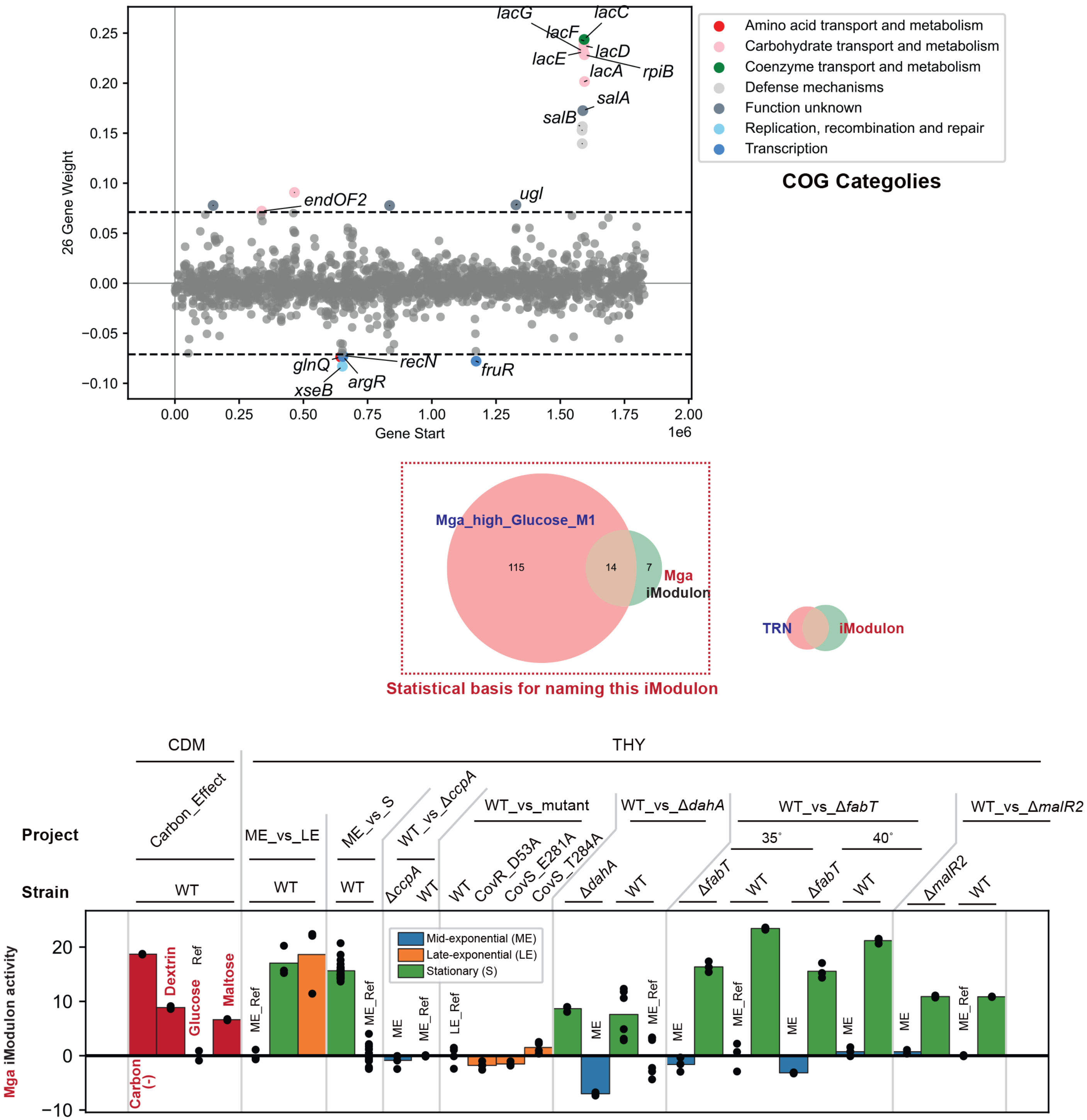

**Unc_10 iModulon**

**Significantly overlapped TRN: ---**

**Biological function: ---**

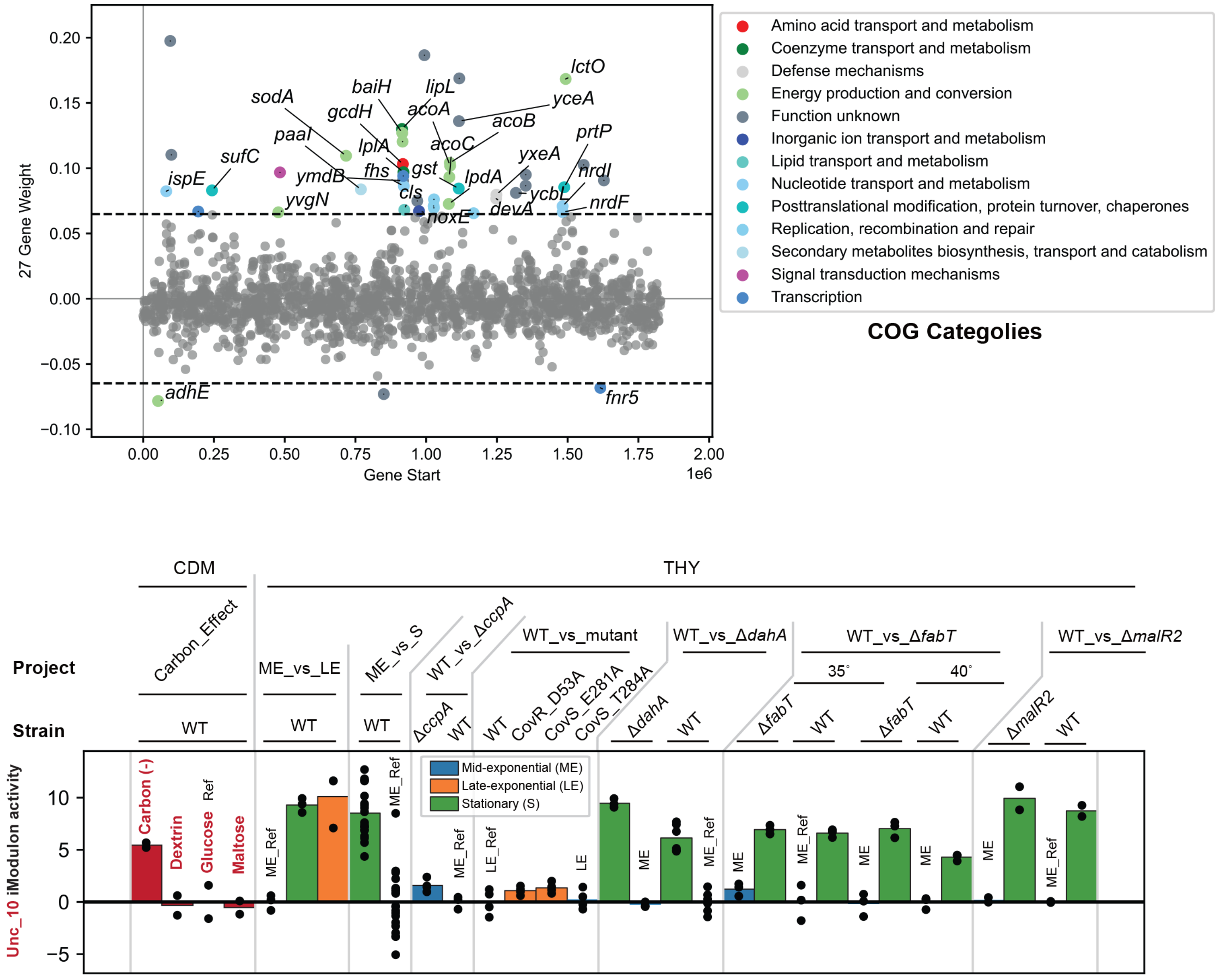

### CovR-2 iModulon

**Significantly overlapped TRN: CovR_1_ME_M1 > CovR_CcpA_M1 > CovR_1_S_M1 > Rgg_exponetional_M49**

**Biological function: Replication**

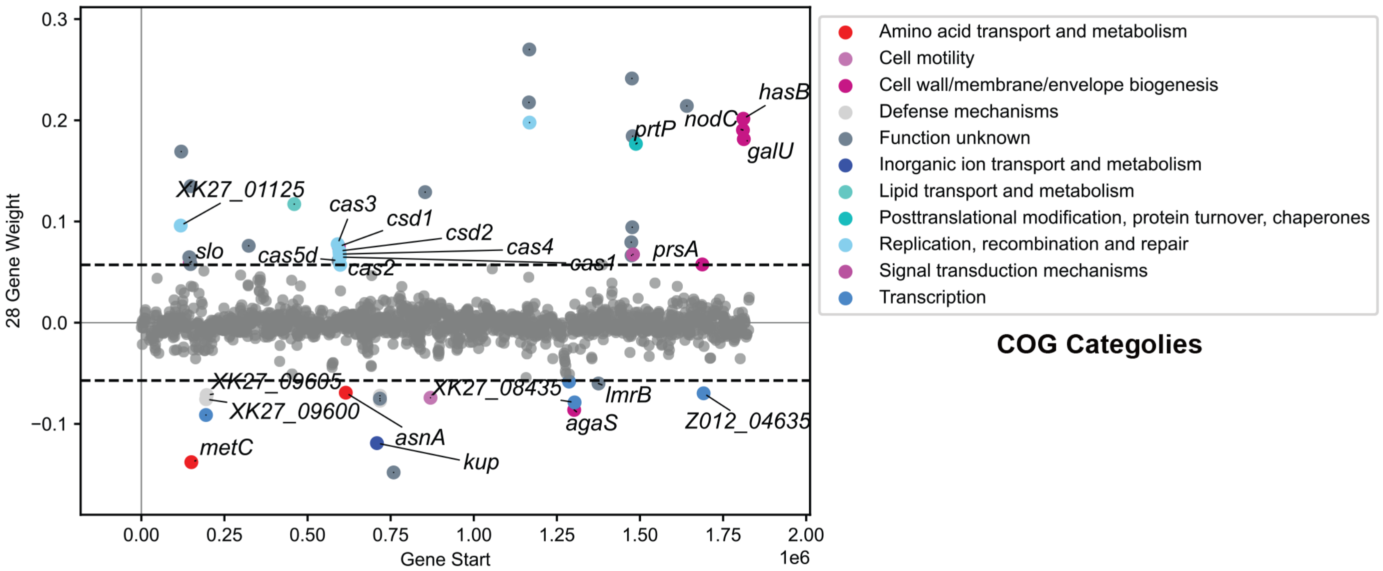


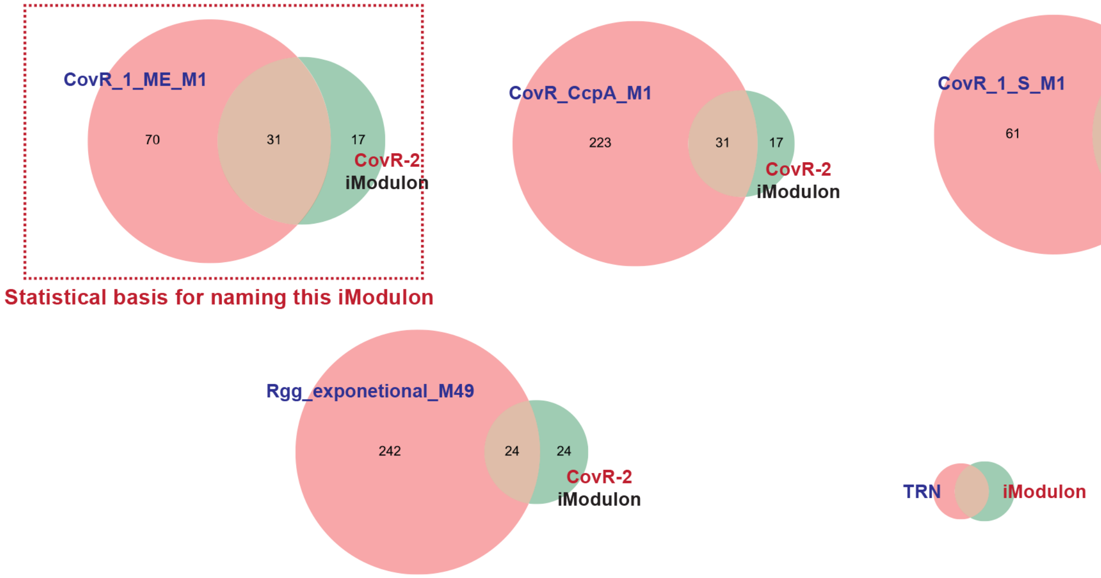


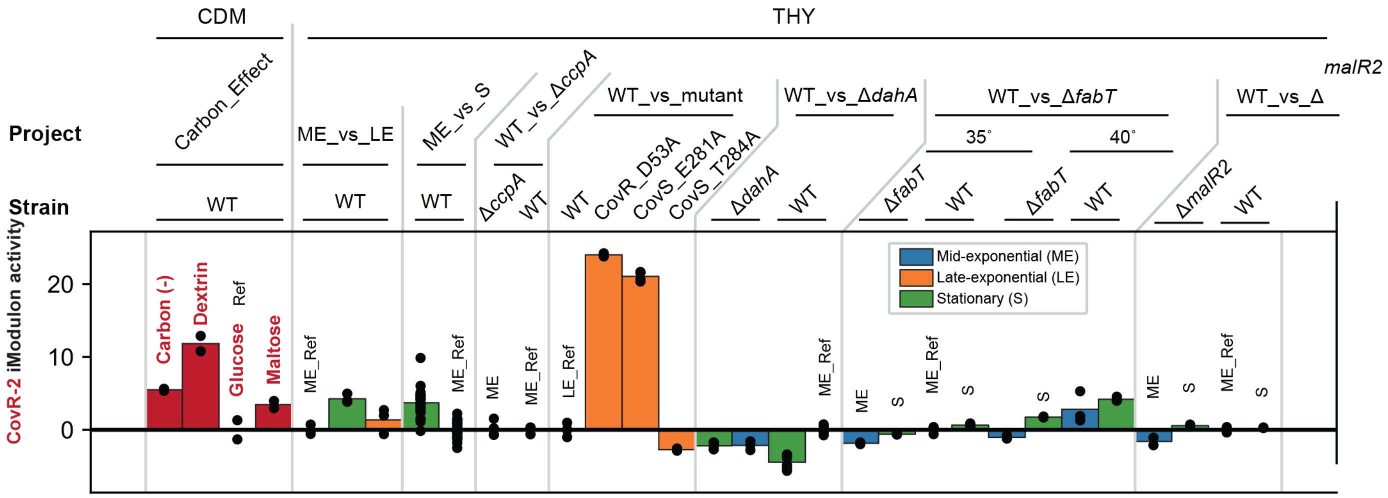

### MalR2 iModulon

#### Significantly overlapped TRN: MalR2

#### Biological function: Carbon use

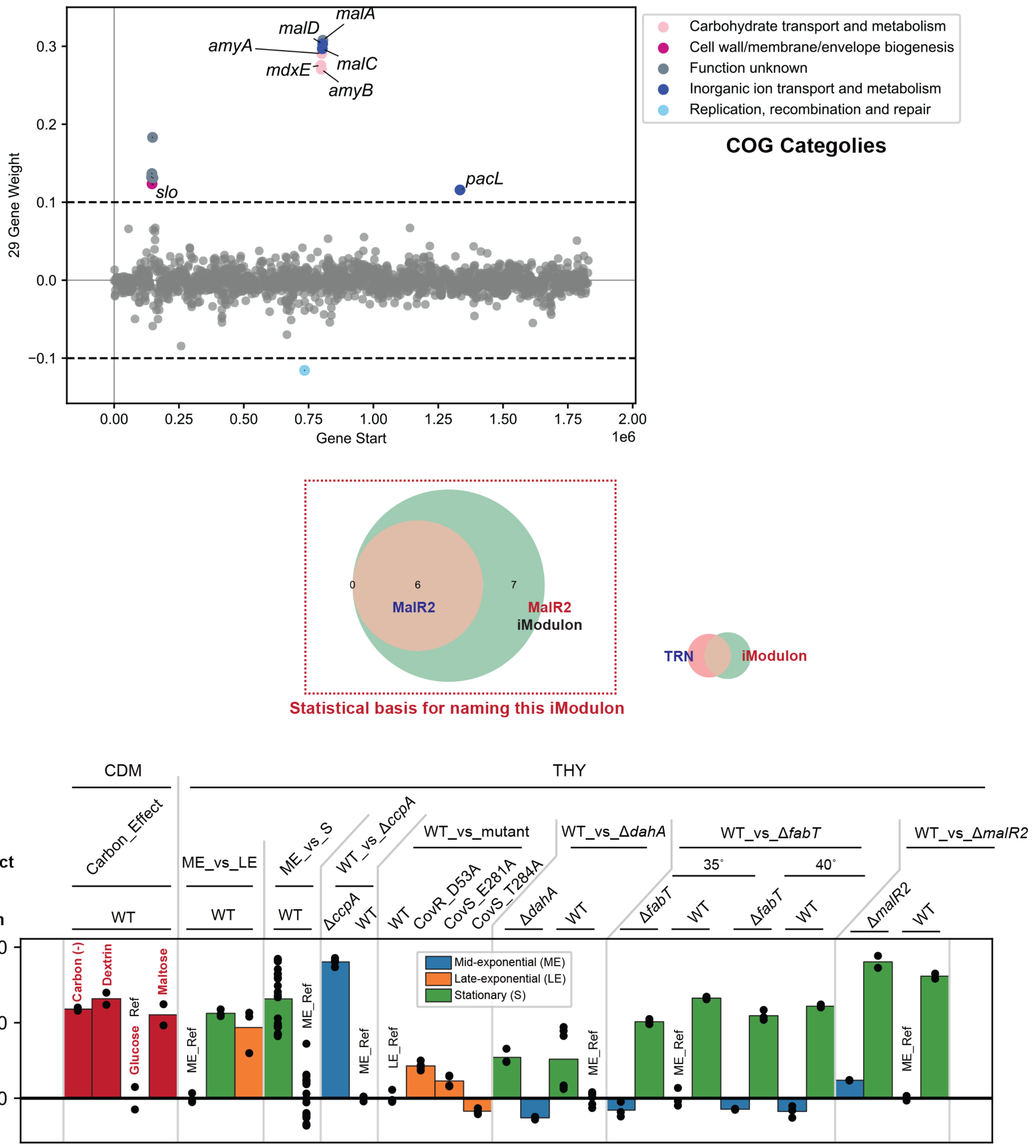

### Phage_3-1 iModulon

**Significantly overlapped TRN: Phage_3 > CovR_CcpA_M1**

**Biological function: Prophages**

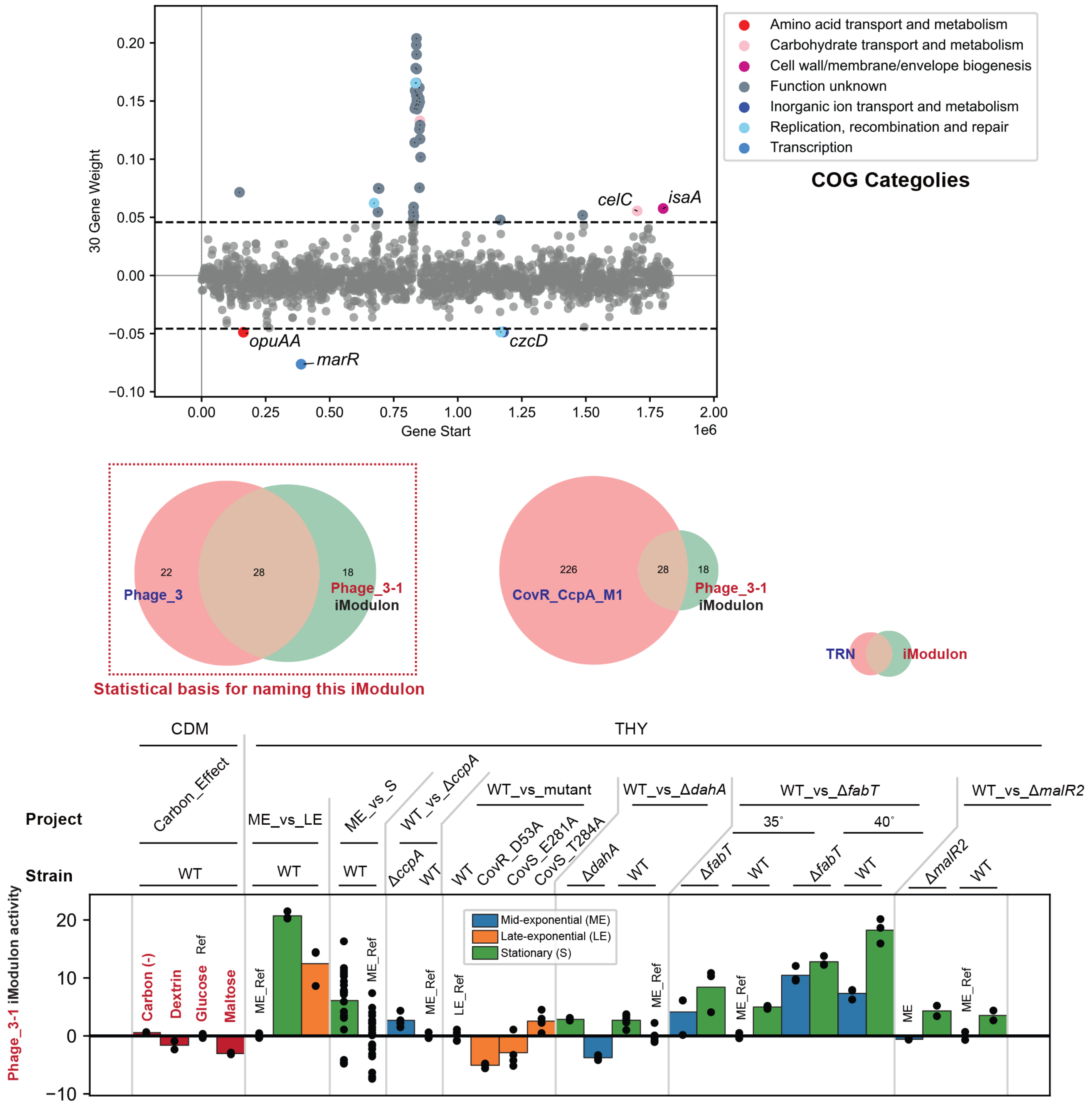

### Phage_2-3 iModulon

**Significantly overlapped TRN: Phage_2 > Phage_3**

**Biological function: Prophages**

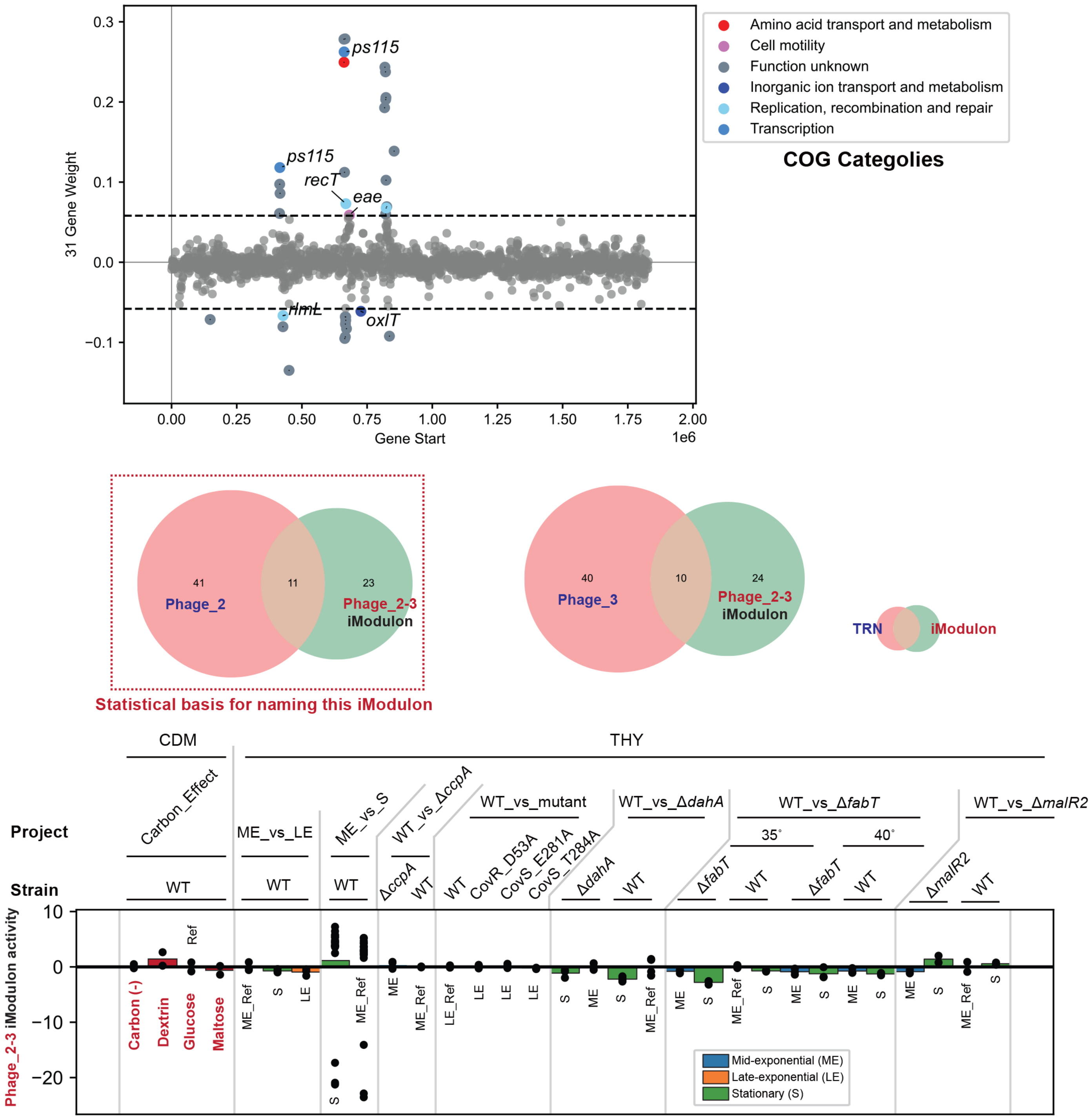

### Unc_11 iModulon

**Significantly overlapped TRN: ---**

**Biological function: Translation**

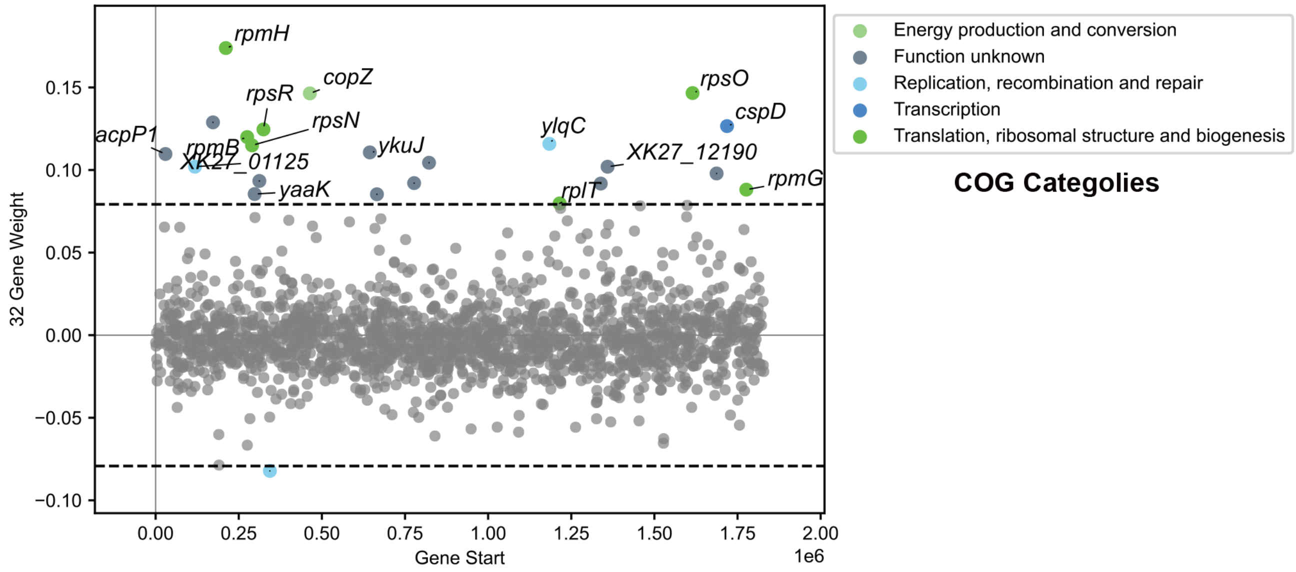


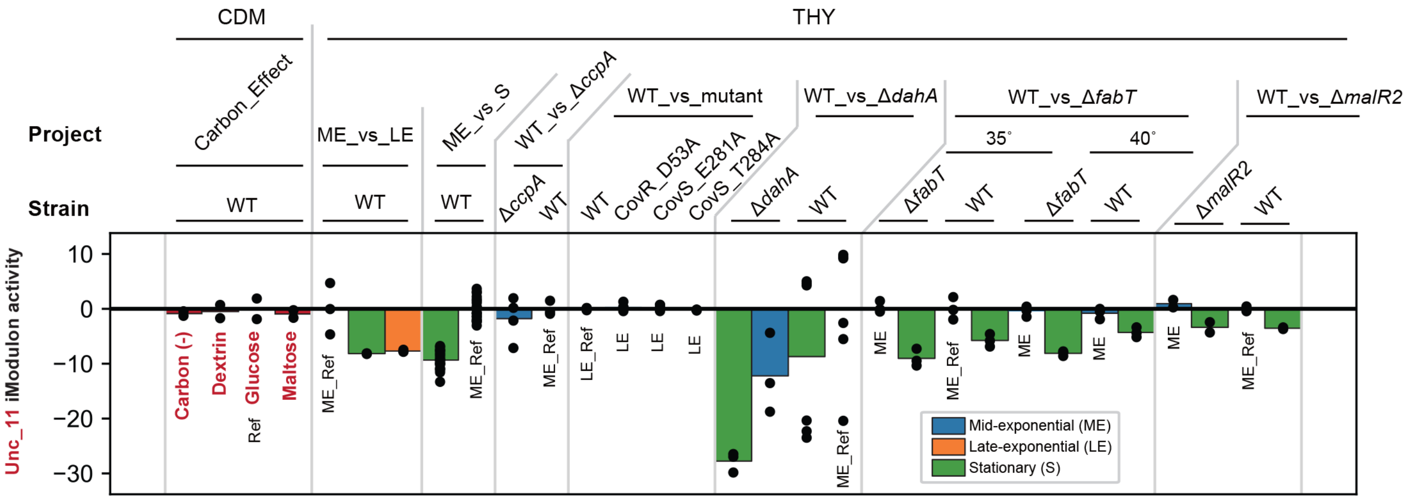

#### Galactose iModulon

**Significantly overlapped TRN: ---**

**Biological function: Carbon use**

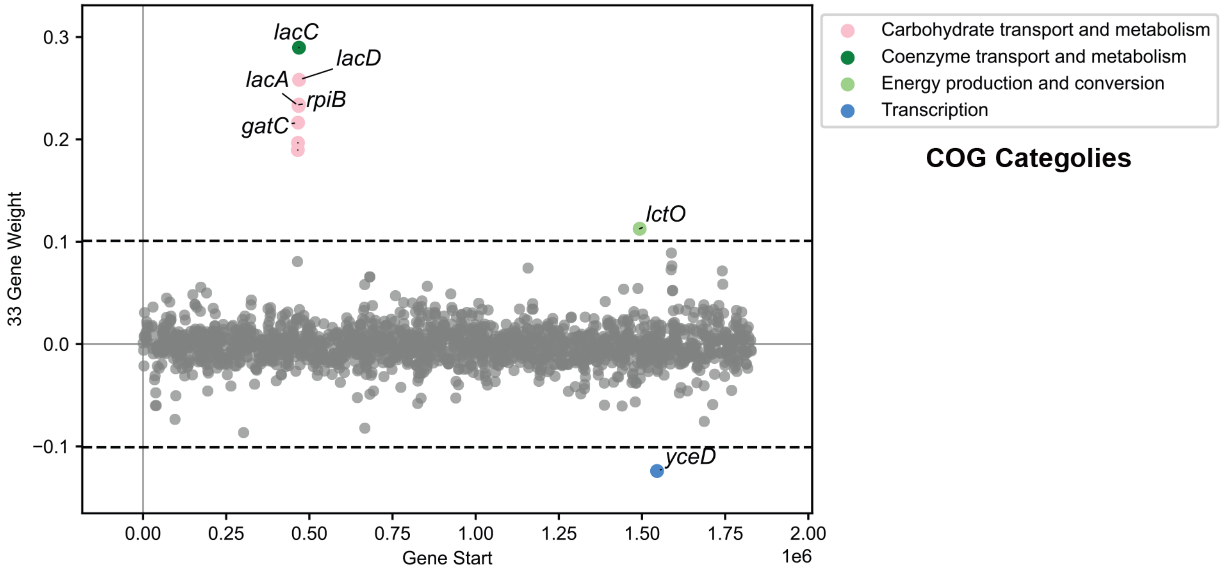


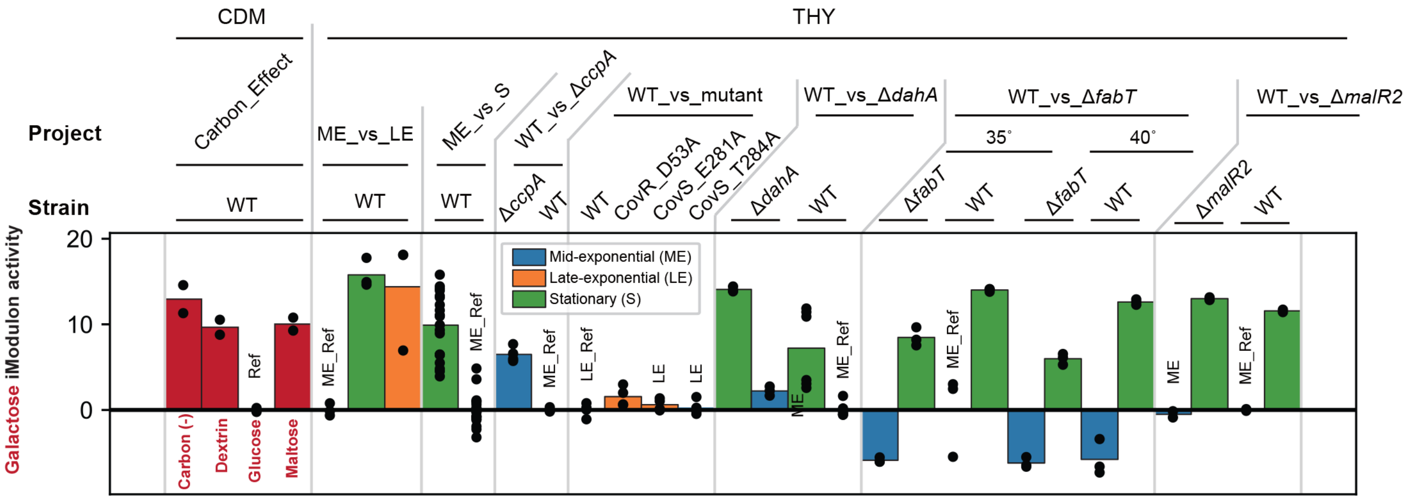

### Transport-1 iModulon

**Significantly overlapped TRN: ---**

**Biological function: Multi-purpose metabolism**

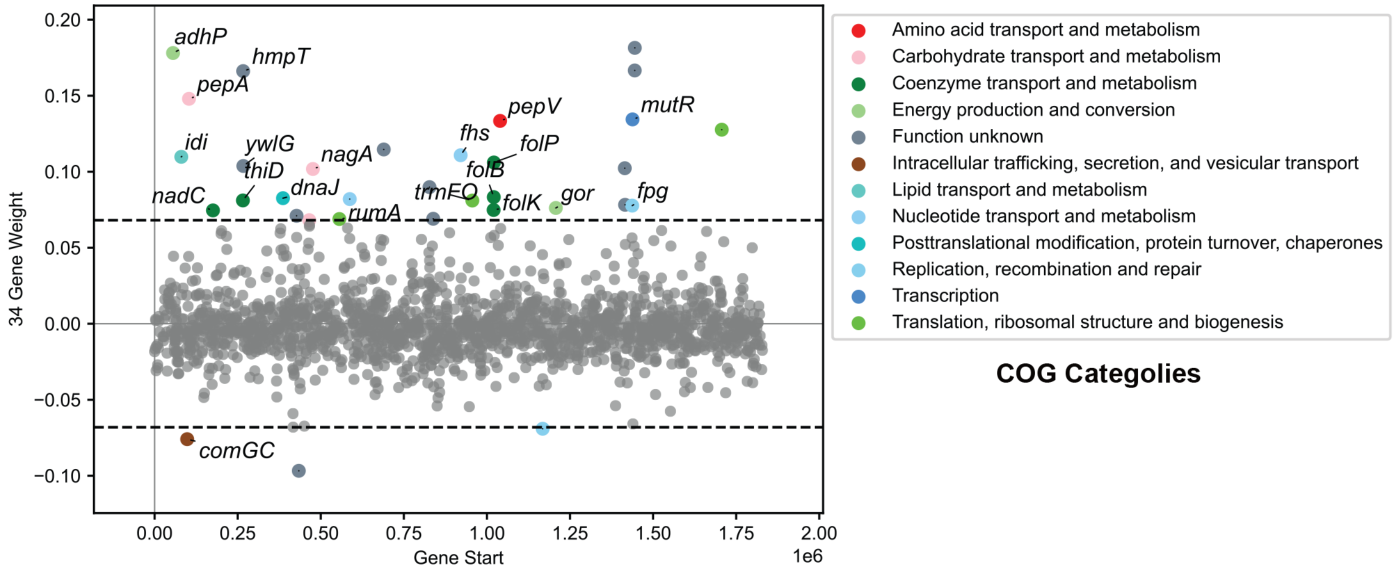


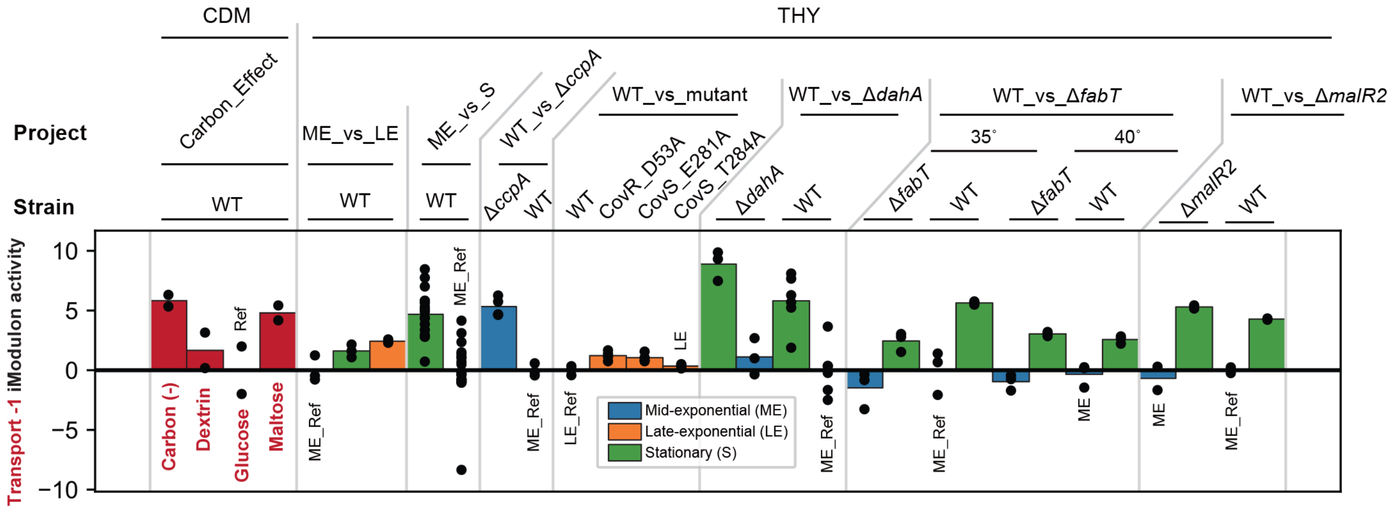

#### YwzG iModulon

**Significantly overlapped TRN: YwzG**

**Biological function: Transport**

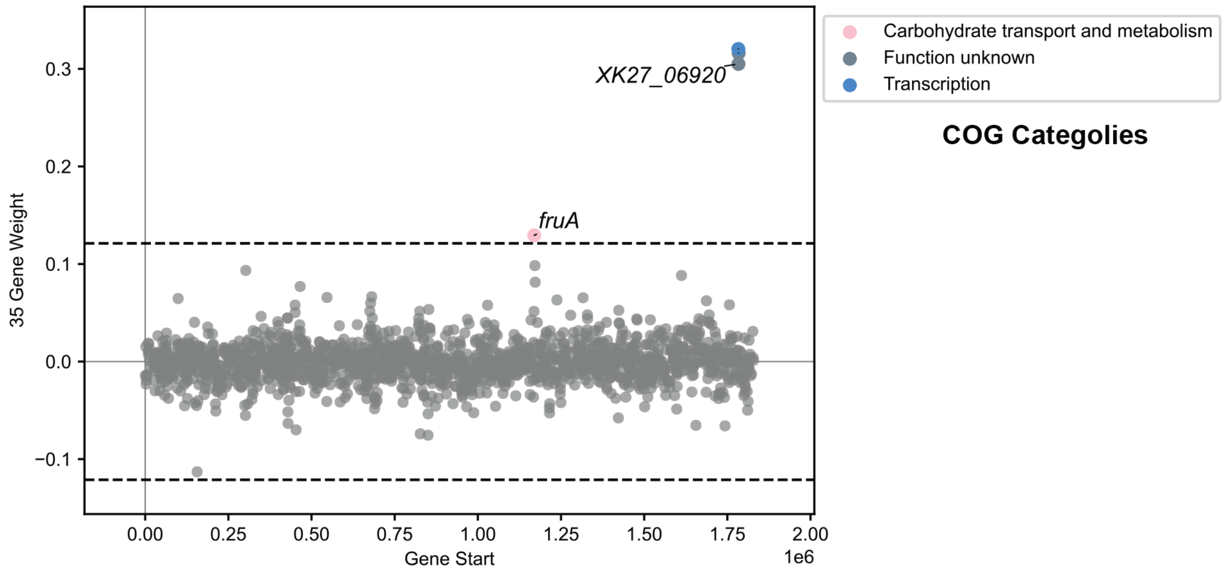


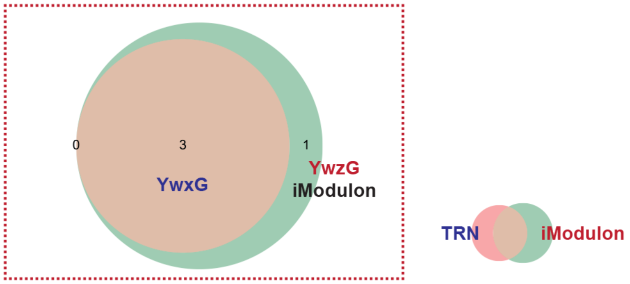


Statistical basis for naming this iModulon

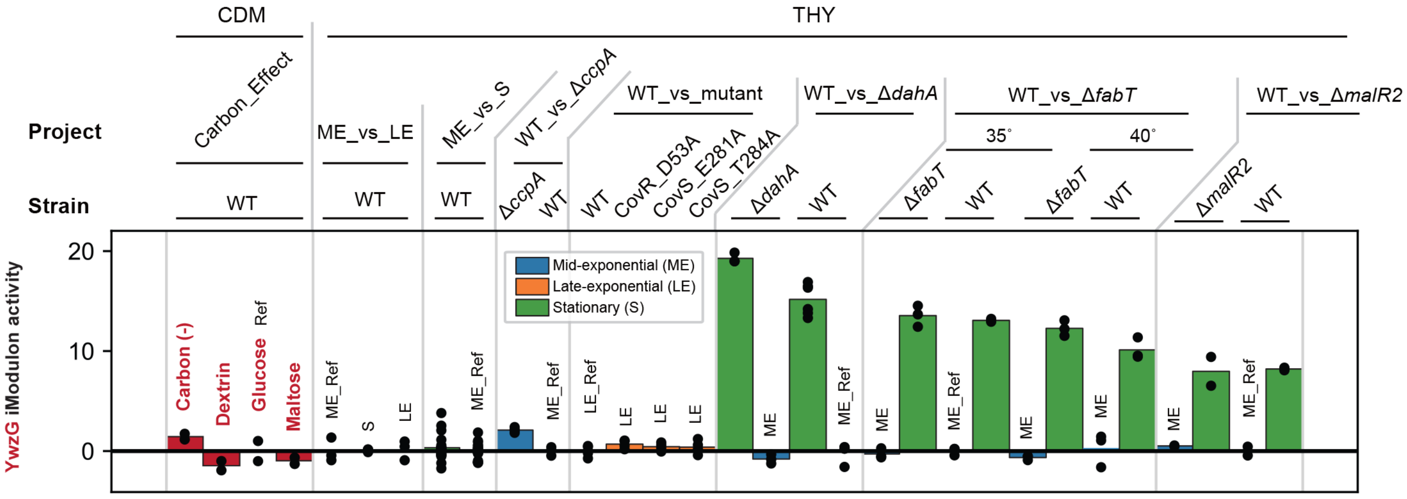

### HrcA iModulon

#### Significantly overlapped TRN: HrcA > CtsR > Nra_transition_M49

#### Biological function: Translation

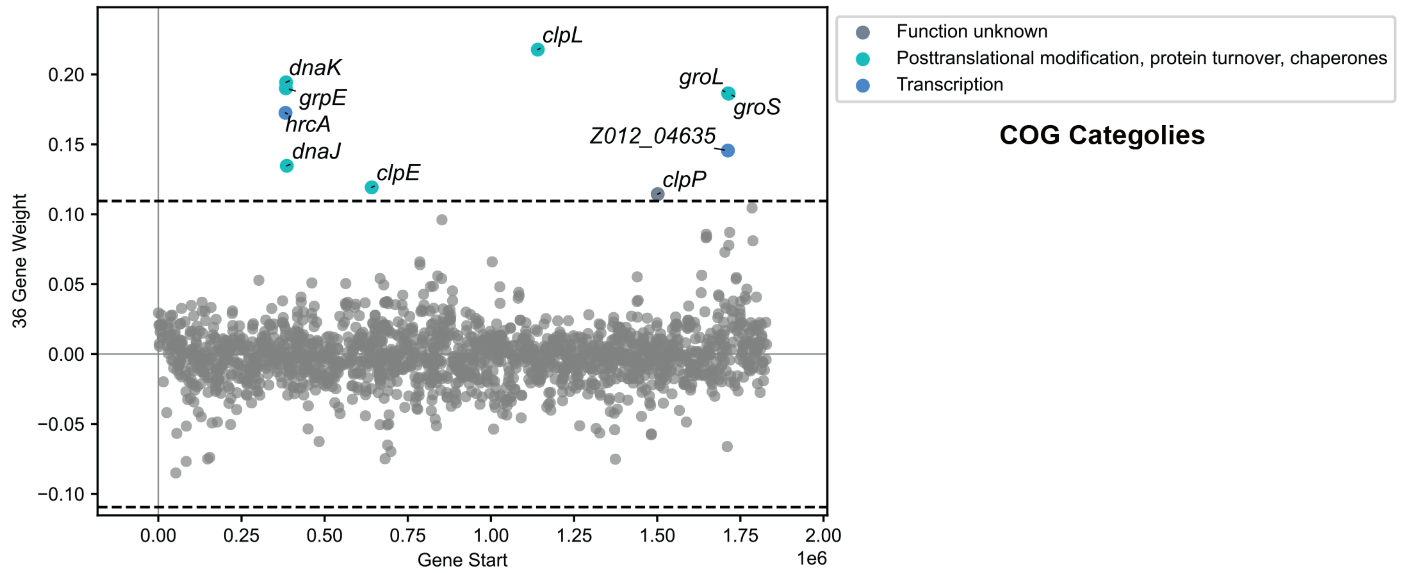


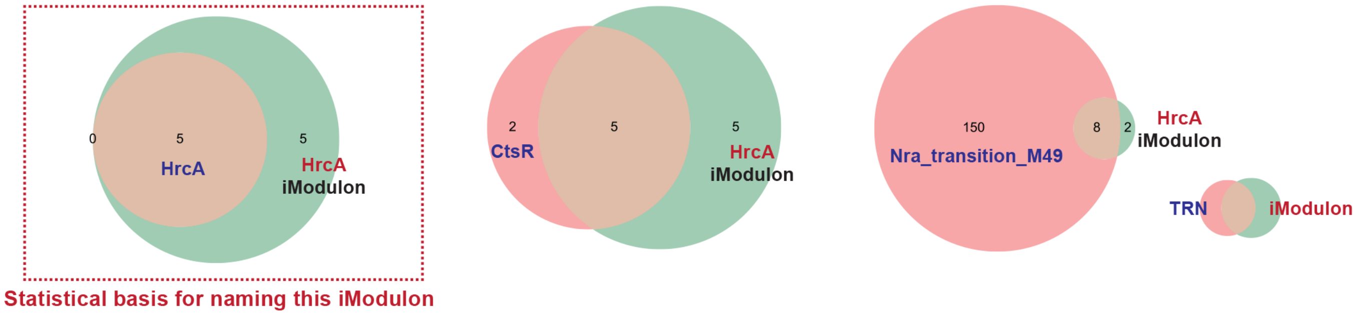


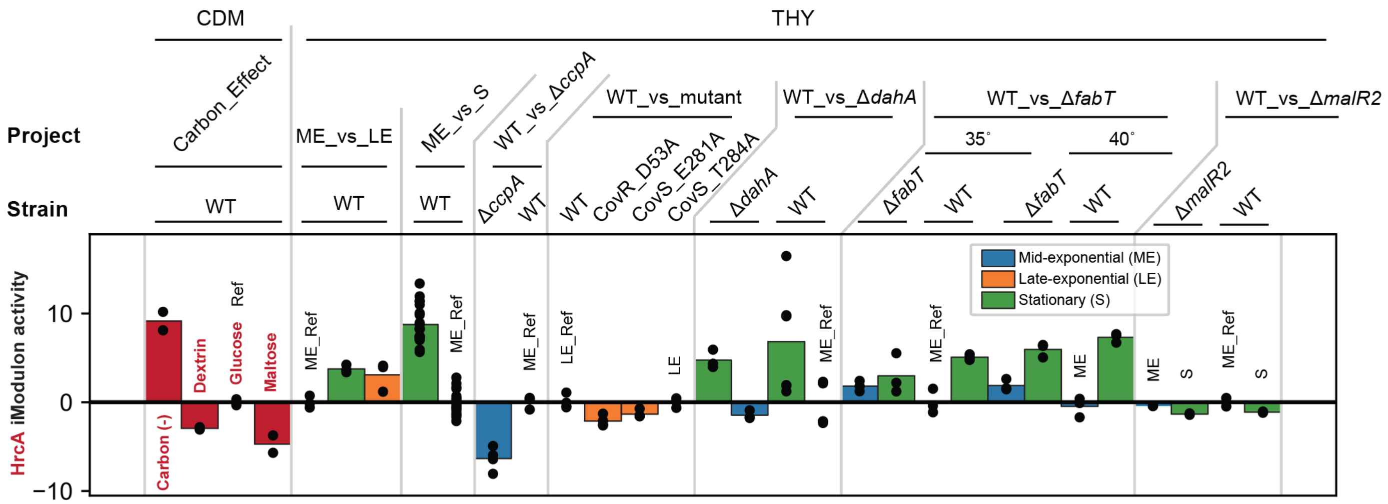

### Phage_3-2 iModulon

**Significantly overlapped TRN: Phage_3 > Phage_1**

**Biological function: Prophages**

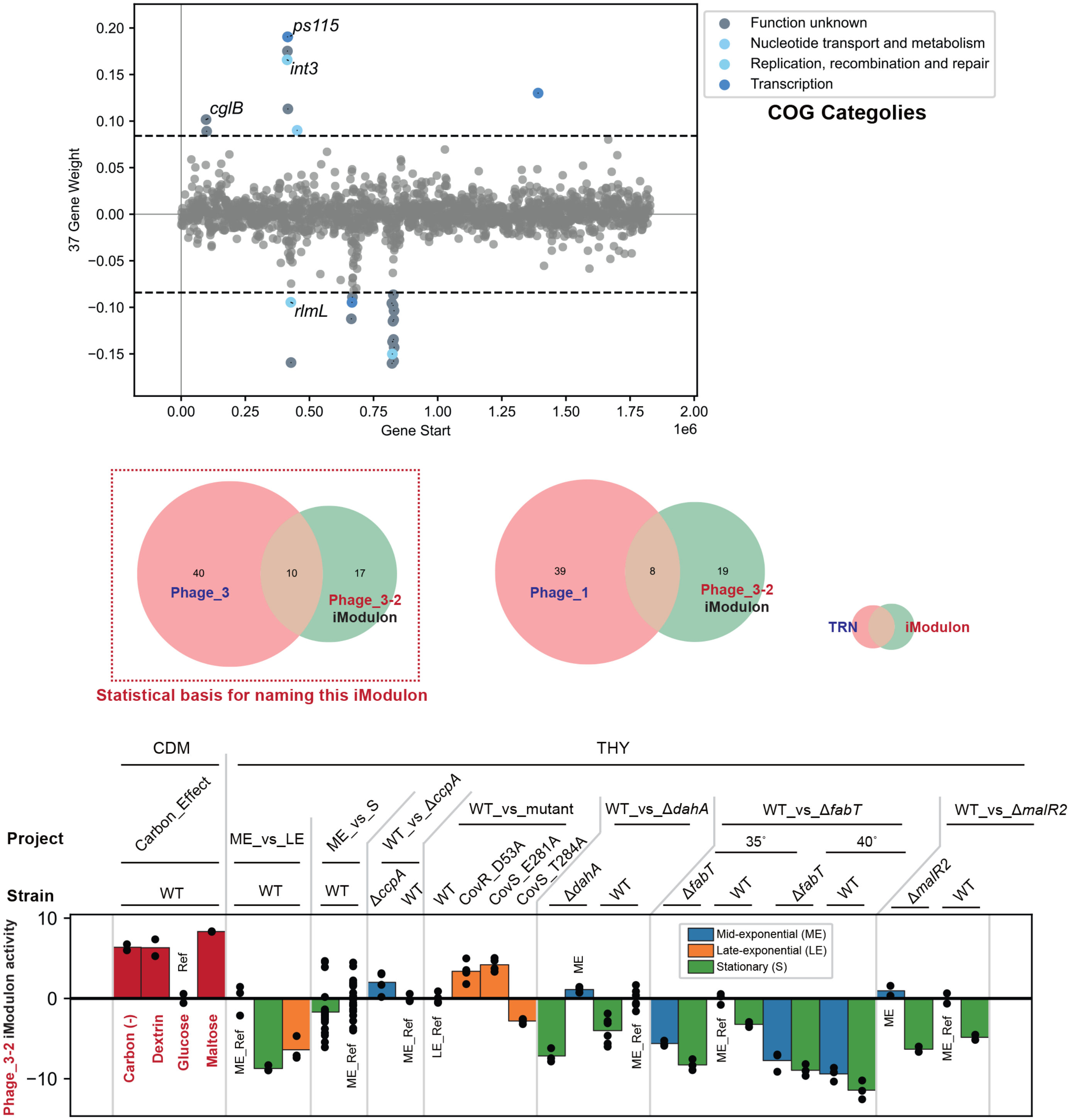

### SpeB iModulon

**Significantly overlapped TRN: ---**

**Biological function: Toxin production**

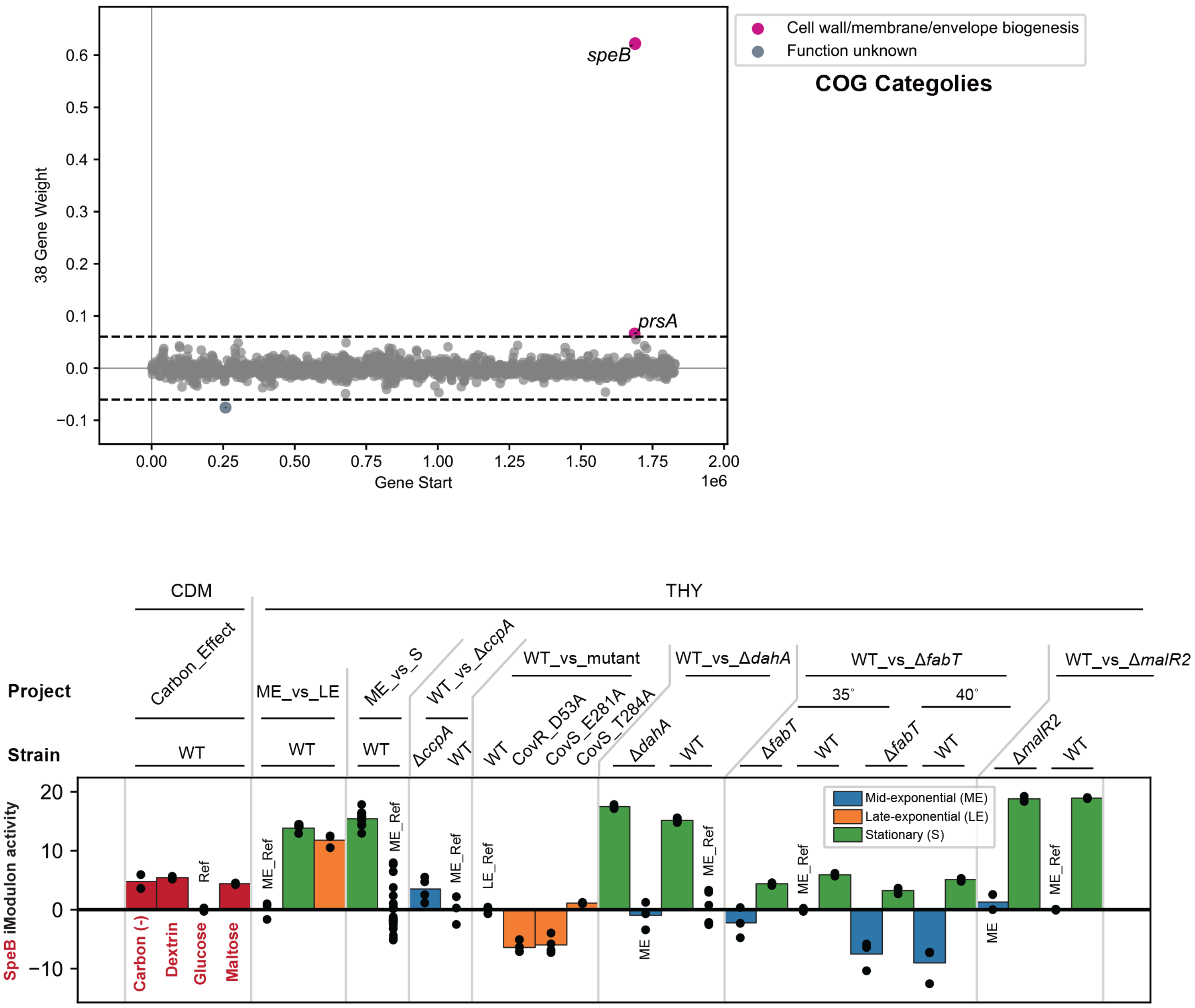

#### Transport-2 iModulon

**Significantly overlapped TRN:**

**Biological function: Multi-purpose metabolism**

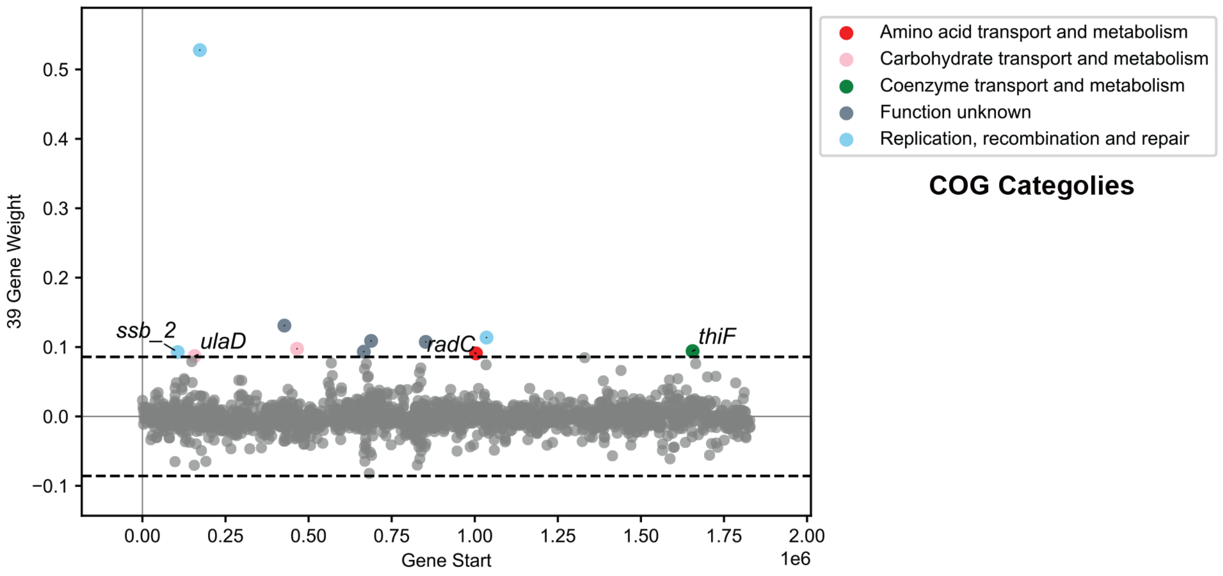


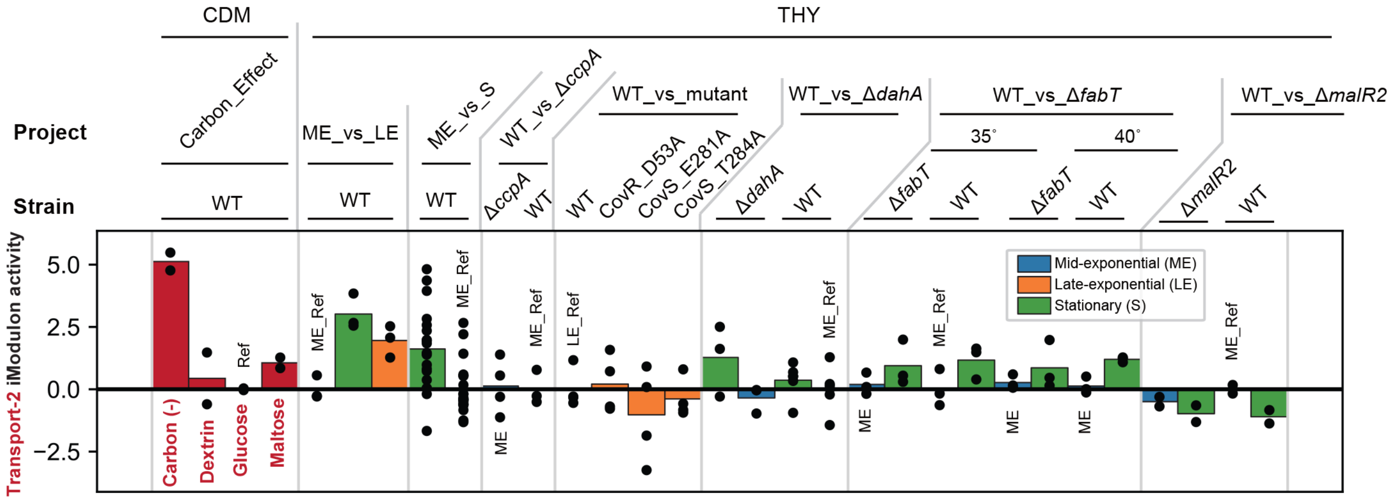

### Unc_12 iModulon

**Significantly overlapped TRN: ---**

**Biological function: General stress**

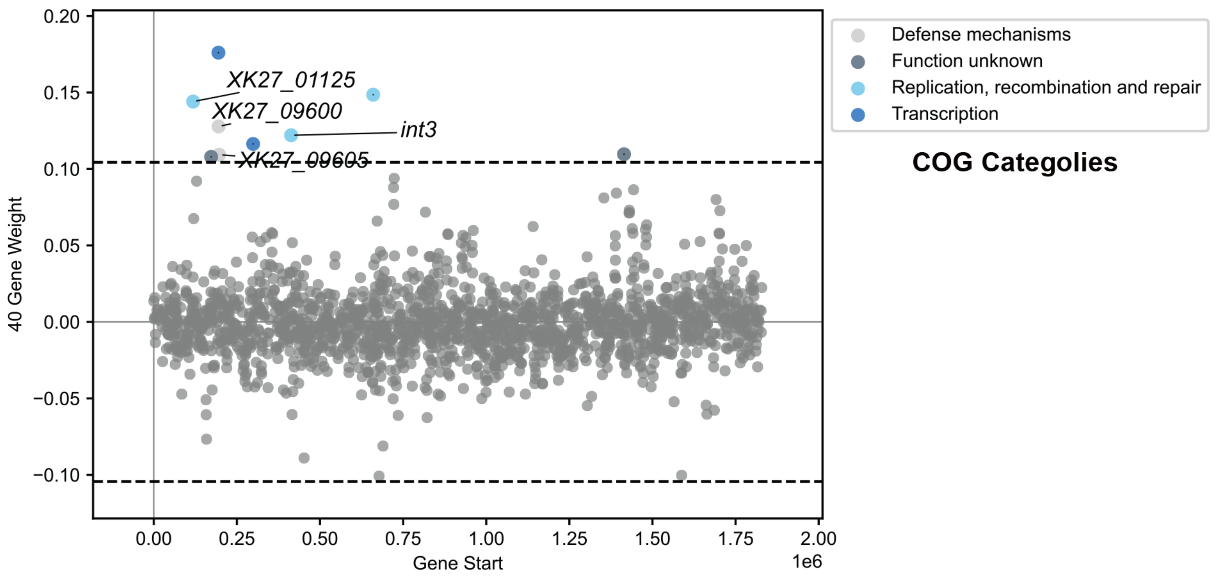


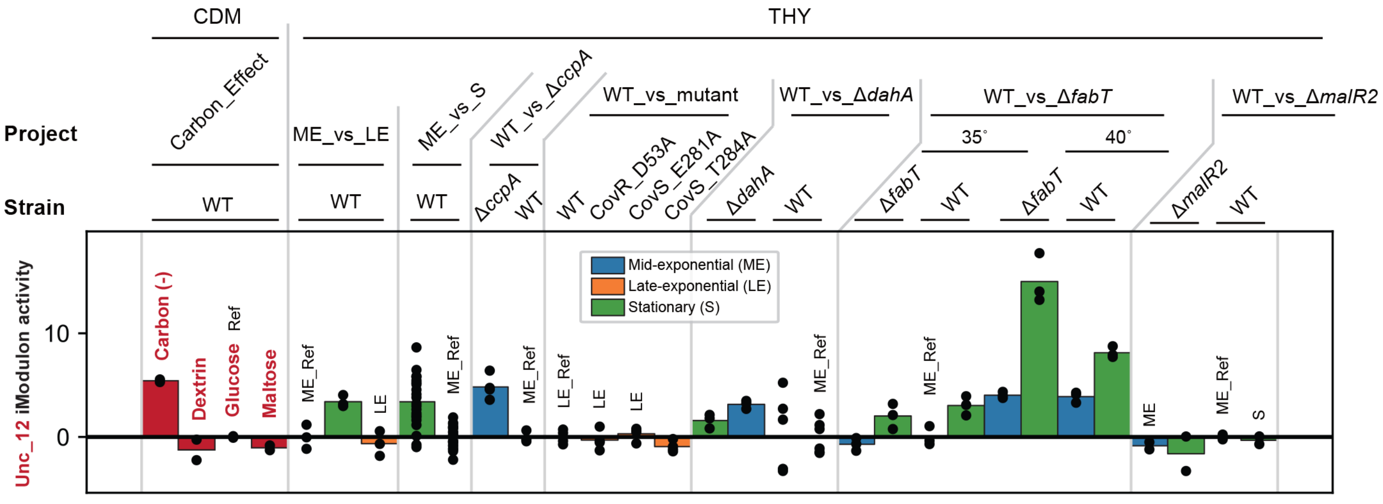

### SLO iModulon

**Significantly overlapped TRN: ---**

**Biological function: Toxin production**

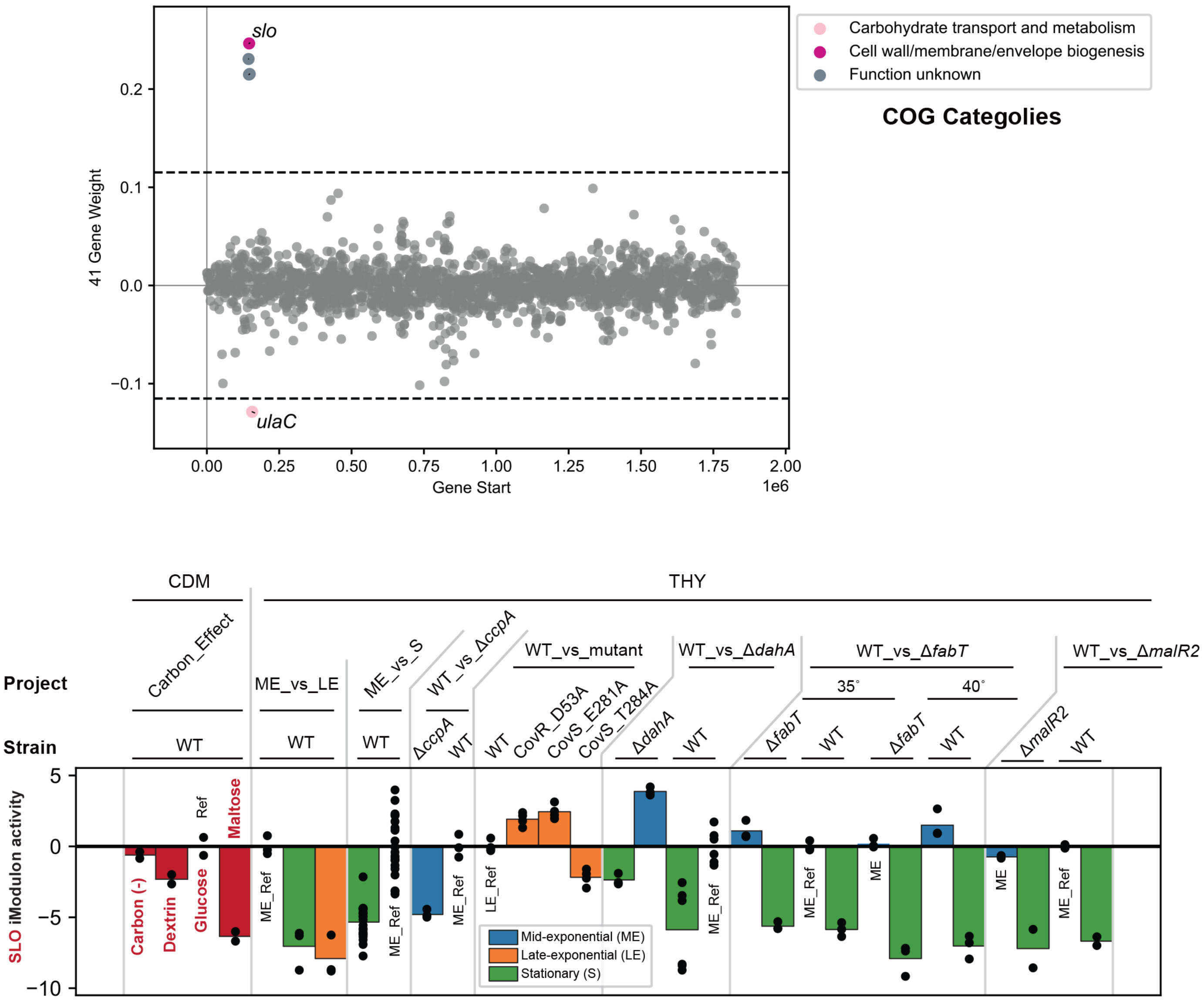
