## Supplementary information 2 for "Elucidation of independently modulated genes in *Streptococcus pyogenes* reveals carbon sources that control its expression of hemolytic toxins"

from 16 RNA-seq data set in this study. ……………………………………….…………6

**1. Supplementary Table**

**Table S1: Primers used in this study**

| **Primer** | **Sequence (5’-3’)** | **Purpose** |
| --- | --- | --- |
| **MalR2KOupF** | CCCAAGCTTTCCTTTTGGAAAATGGTTTATGA | Construction of pHY304-*malR2*KO |
| **MalR2KOupR** | CCAGTGATTTTTTTCTCCATCATTTTGTTATATCCGAATTC | Construction of pHY304-*malR2*KO |
| **MalR2KOdownF** | GAGTGGCAGGGCGGGGCGTAACGCCTGAGAATTAACCTATC | Construction of pHY304-*malR2*KO |
| **MalR2KOdownR** | ATAGTTTAGCGGCCGCTCCCTTGGAACGGATTGTAA | Construction of pHY304-*malR2*KO |
| **MalR2KOcatF** | ATGGAGAAAAAAATCACTGGATATACC | Construction of pHY304-*malR2*KO |
| **MalR2KOcatR** | TTACGCCCCGCCCTGCCACTCATCGCA | Construction of pHY304-*malR2*KO |
| **MalR2KOconfirmF** | GCGAAGGAACTGCTTACGTC | Confirmation of *malR2* deletion |
| **MalR2KOconfirmR** | GCCGAGCTCAGTTTACTTGG | Confirmation of *malR2* deletion |

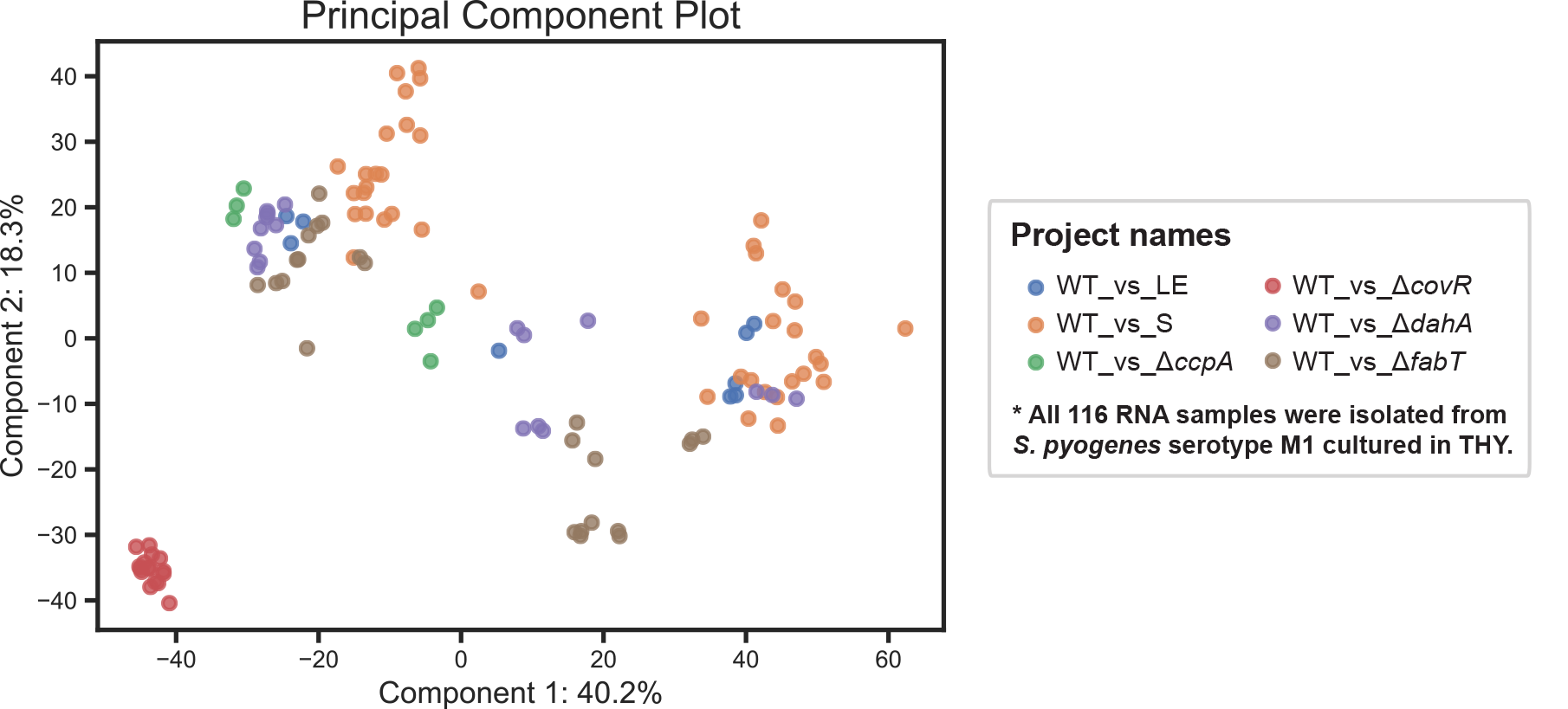
**2. Supplementary Figures**

**Figure S1. Diversity of 116 RNA-seq data sets used for ICA decomposition.** Loadings of the first two principal components (PC). The variation in locations across 116 RNA-seq samples demonstrates the diversity of the compendium.

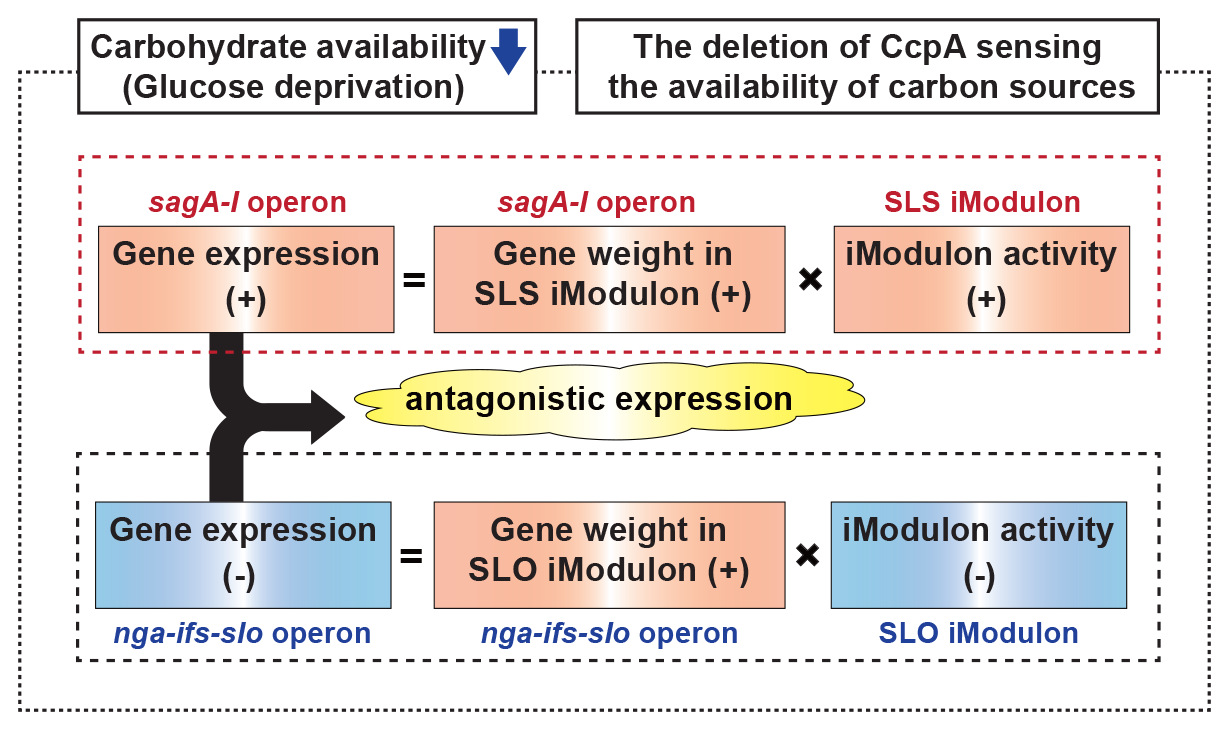

**Figure S2. Conditions that *S. pyogenes* may show the antagonistic expression of *sag A-I* and *nga-ifs-slo* operons.**

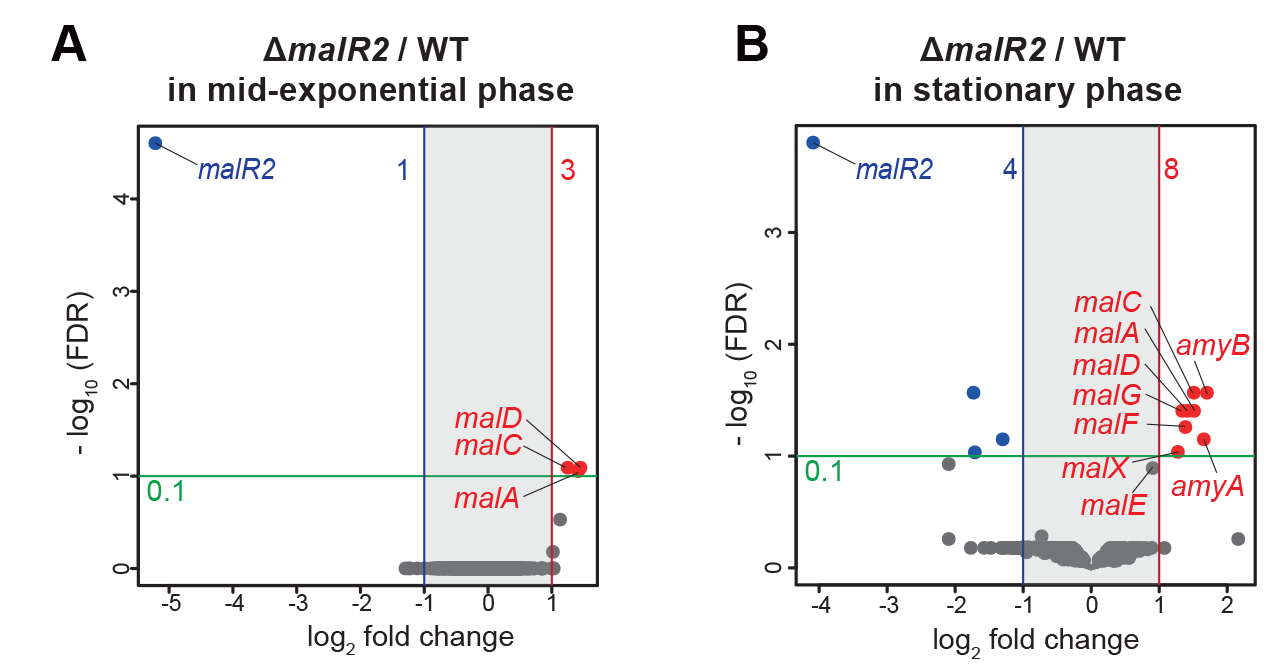

**Figure S3. The actual MalR2 regulon.** Differentially expressed genes (DEGs) from comparisons of the Δ*malR2* mutant and WT strains at (A) mid-exponential, and (B) stationary growth phases in THY broth. Colored circles indicate significantly upregulated (red) and downregulated (blue) genes (absolute log2 fold change, > 1; adjusted P < 0.1).

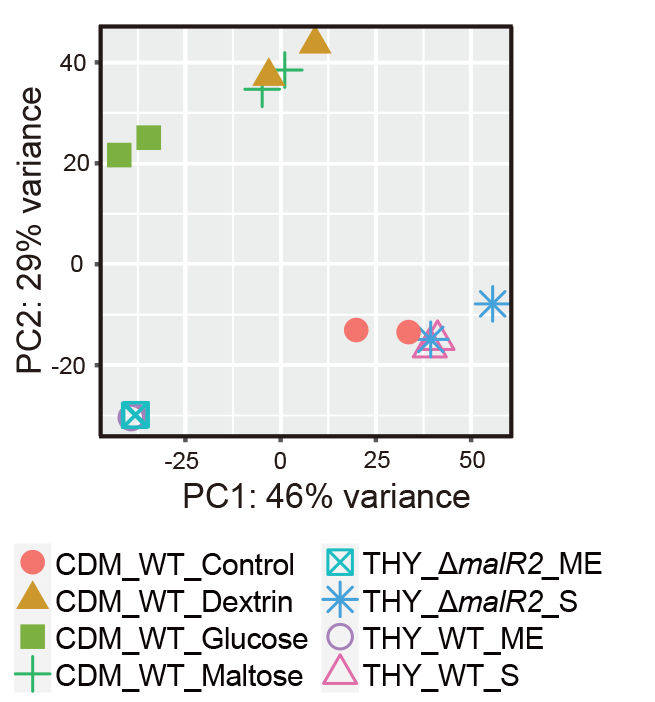

**Figure S4. Principal component analysis (PCA) plot of TPM data from 16 RNA-seq data set in this study.** ME, Mid-Exponential growth phase. S, Stationary growth phase.
